# Striatal BOLD and midfrontal theta power express motivation for action

**DOI:** 10.1101/2020.09.11.292870

**Authors:** Johannes Algermissen, Jennifer C. Swart, René Scheeringa, Roshan Cools, Hanneke E.M. den Ouden

**Author notes:** Contract Address: Montessorilaan 3B, 6525 HR Nijmegen, (J.A.); (H.E.M.d.O.).

## Abstract

Action selection is biased by the valence of anticipated outcomes. To assess mechanisms by which these motivational biases are expressed and controlled, we measured simultaneous EEG-fMRI during a motivational Go/NoGo learning task (N=36), leveraging the temporal resolution of EEG and subcortical access of fMRI. VmPFC BOLD encoded cue valence, importantly predicting trial-by-trial valence-driven response speed differences and EEG theta power around cue onset. In contrast, striatal BOLD encoded selection of active Go responses and correlated with theta power around response time. Within trials, theta power ramped in the fashion of an evidence accumulation signal for the value of making a ‘Go’ response, capturing the faster responding to reward cues. Our findings reveal a dual nature of midfrontal theta power, with early components reflecting the vmPFC contribution to motivational biases, and late components reflecting their striatal translation into behavior, in line with influential recent “value of work” theories of striatal processing.

---

Learning from rewards and punishments allows us to adapt action selection to our environment. At the same time, our responses are also shaped by seemingly automatic action tendencies that appear to be innate or acquired very early in development. A prime example are motivational action biases (also called “Pavlovian” biases), referring to the tendency to invigorate actions when there is a prospect of reward, but to hold back when there is a threat of punishments (Dayan et al. 2006; Guitart-Masip, Duzel, et al. 2014). Such an action ‘prior’ allows for fast responding which is often adaptive, given that reward pursuit typically requires active responses and threat avoidance typically requires action suppression. However, in environments in which these relationships do not hold, the action triggered by the motivational bias can interfere with the normative optimal action (i.e., the action that maximizes rewards and minimizes punishments) and needs to be suppressed. Keeping a balance between hardwired action tendencies and action values flexibly learned from experience can be challenging, and deficits have been linked to psychiatric disorders such as addiction, depression, trauma symptoms, and social anxiety (Garbusow et al. 2016, 2019; Huys et al. 2016; Mkrtchian et al. 2017; Ousdal et al. 2018). Thus, it is important to understand the mechanism by which humans selectively rely on these biases when they are helpful and suppress them when they are not. In this study, we investigate the neural interactions that accompany (un)successful suppression of motivational biases when needed.

Motivational biases have been hypothesized to arise from dopaminergic effects in the basal ganglia (Frank 2005; Collins and Frank 2014). In line with the putative role of the striatum in driving motivational biases, striatal fMRI BOLD signal has been found modulated by cues signaling the prospect of reward (O’Doherty et al. 2003; Tobler et al. 2007; Niv et al. 2012) and dopaminergic medication has been found to modulate these motivational biases (Guitart-Masip, Chowdhury, et al. 2012; Guitart-Masip, Economides, et al. 2014; Swart et al. 2017; van Nuland et al. 2020).

When actions triggered by motivational biases conflict with the action required to obtain a desired outcome, individuals need to detect this motivational conflict and mobilize control mechanisms to suppress biases. This role has traditionally been attributed to interactions between the medial and lateral prefrontal cortex particularly the anterior cingulate cortex (ACC). More recent approaches regard the medial frontal cortex, in particular the ACC, as a central decision hub evaluating whether to recruit cognitive control or not (Shenhav et al. 2013, 2016). Recruiting cognitive control is perceived as costly, but potentially worth the effort to overcome biases and select the optimal action.

Bursts of oscillatory synchronization in the theta range (4-8 Hz) over midfrontal cortex have been proposed as an electrophysiological signature of conflict processing. Recently, we and others have observed this signal also when participants successfully overcame motivational conflict between task-appropriate and task-inappropriate, bias-triggered actions (Cavanagh et al. 2013; Swart et al. 2018; Csifcsák et al. 2019). However, the downstream mechanisms by which midfrontal signals prevent the behavioral expression of motivational biases, putatively driven by the striatum, remain elusive. Previous research has focused on tasks with two active responses (e.g. the Stroop task), where midfrontal theta has been suggested to activate the subthalamic nucleus, which raises the threshold of striatal input needed to elicit an action and thus prevents impulsive actions (Zavala et al. 2014; Frank et al. 2015; Aron et al. 2016; Herz et al. 2016). In contrast, in our task, not only reward-triggered action, but also punishment-triggered inhibition needs to be overcome, which might be achieved by a direct attenuation of the cortical inputs into the striatum implemented via recurrent fronto-striatal loops (Alexander et al. 1986; Mink 1996; Gurney et al. 2001). In the current study, we tested the hypothesis that midfrontal theta power is associated with an attenuation of subcortical signals that encode motivational biases when those need to be suppressed.

To test this hypothesis, participants performed a Motivational Go/NoGo learning task known to elicit motivational biases in humans (Swart et al. 2017, 2018) while simultaneously recording fMRI and scalp EEG. The temporal precision of EEG allowed us to separate cue-induced conflict signals from later, response-locked signals. We tested the specific hypotheses that i) striatal BOLD encodes cue valence, ii) this valence signaling is attenuated when participants successfully overcome motivational biases, and iii) valence signal attenuation correlates with midfrontal theta power. Specifically, using BOLD signal to predict EEG power at different time points allowed us to separate regions involved in (early) valence cue processing vs. (later) response biases.

## Materials and Methods

### Participants

Thirty-six participants (*Mage* = 23.6, age range 19–32; 25 women, all right-handed) performed the motivational Go/NoGo learning task while simultaneous EEG and fMRI were recorded. Sample size was based on previous EEG studies (Cavanagh et al. 2013; Swart et al. 2018) accounting for potential dropout. The study was approved by the local ethics committee (CMO2014/288; Commissie Mensgebonden Onderzoek Arnhem-Nijmegen). All participants provided written informed consent. Exclusion criteria comprised claustrophobia, allergy to gels used for EEG electrode application, hearing aids, impaired vision, colorblindness, history of neurological or psychiatric diseases (including heavy concussions and brain surgery), epilepsy, metal parts in the body, or heart problems.

Participants attended a single three-hour recording session and were compensated for participation (€30). Additionally, they received a performance-dependent bonus (range €0–5, *Mbonus* = €1.28, *SD*bonus = 1.54). The reported behavioral and EEG results are based on all 36 participants. For two participants, fMRI co-registration failed due to excessive orbitofrontal distortion in the T1 image; thus, fMRI results are based on 34 participants (*Mage* = 23.5, age range 19–32; 25 women). fMRI-informed EEG results are based on 29 participants (Mage = 23.4, 21 women): Apart from the two participants for whom co-registration failed, we excluded four further participants who exhibited strong head motion (i.e., at least 5 volumes with relative displacement larger than the voxel size of 2 mm). These four participants also exhibited stronger overall head motion (i.e., mean relative displacement across all volumes), *M* = 0.213, *SD* = 0.084, compared to the other participants, *M* = 0.088, *SD* = 0.040. Head motion is a particular problem in the EEG-fMRI combined analysis, as it will lead to high and spatially uniform correlations between fMRI and EEG data. Indeed, for these participants, regression weights for all regressors were an order of magnitude larger than for the other participants and largely uniform across time and frequency. We repeated behavioral, EEG and fMRI results for the subgroup of these 29 participants in Supplementary Material S01; conclusions were identical unless mentioned otherwise in the main text.

### Motivational Go/NoGo learning task

Participants performed the Motivational Go/NoGo learning task as detailed in (Swart et al. 2018) with trial timings adjusted to the fMRI acquisition (Fig. 1). The task was programmed in MATLAB 2014b (The MathWorks, Natick, MA, United States) / Psychtoolbox-3.0.13. Each trial started with the presentation of one of eight cues (a colored geometric shape) in the center of the display (Fig. 1A). This cue determined the outcome valence (Win reward/ Avoid punishment) and which action (Left Go/ Right Go/ NoGo) was required for the desired outcome (win reward/ avoid punishment). Participants had to learn both the valence of the cue and the required action from trial-and-error (Fig. 1B). For Win cues, participants should aim to win a reward and avoid neutral outcomes, while for Avoid cues, they should aim to achieve neutral outcomes and avoid punishments. Participants could respond by pressing a left button (Left Go), right button (Right Go), or choose not to press (NoGo) while the cue was on screen for 1300 ms. After a pseudorandomly jittered fixation period of 1400– 2600 ms, the outcome was presented for 750 ms in the form of money falling into a can (reward), money falling out of a can (loss / punishment), or just the can (neutral outcome). Outcomes were probabilistic so that the optimal action led to the desired outcome in only 80% of trials, while suboptimal actions led to desired outcomes in 20% of trials (Fig. 1C). Each trial ended with a pseudorandomly jittered inter-trial interval of 1250–2000 ms, resulting in an overall trial length of 4700–6650 ms. Analyses of learning and outcome-based activity in EEG and fMRI will be reported in a separate publication.

**Fig 1.**
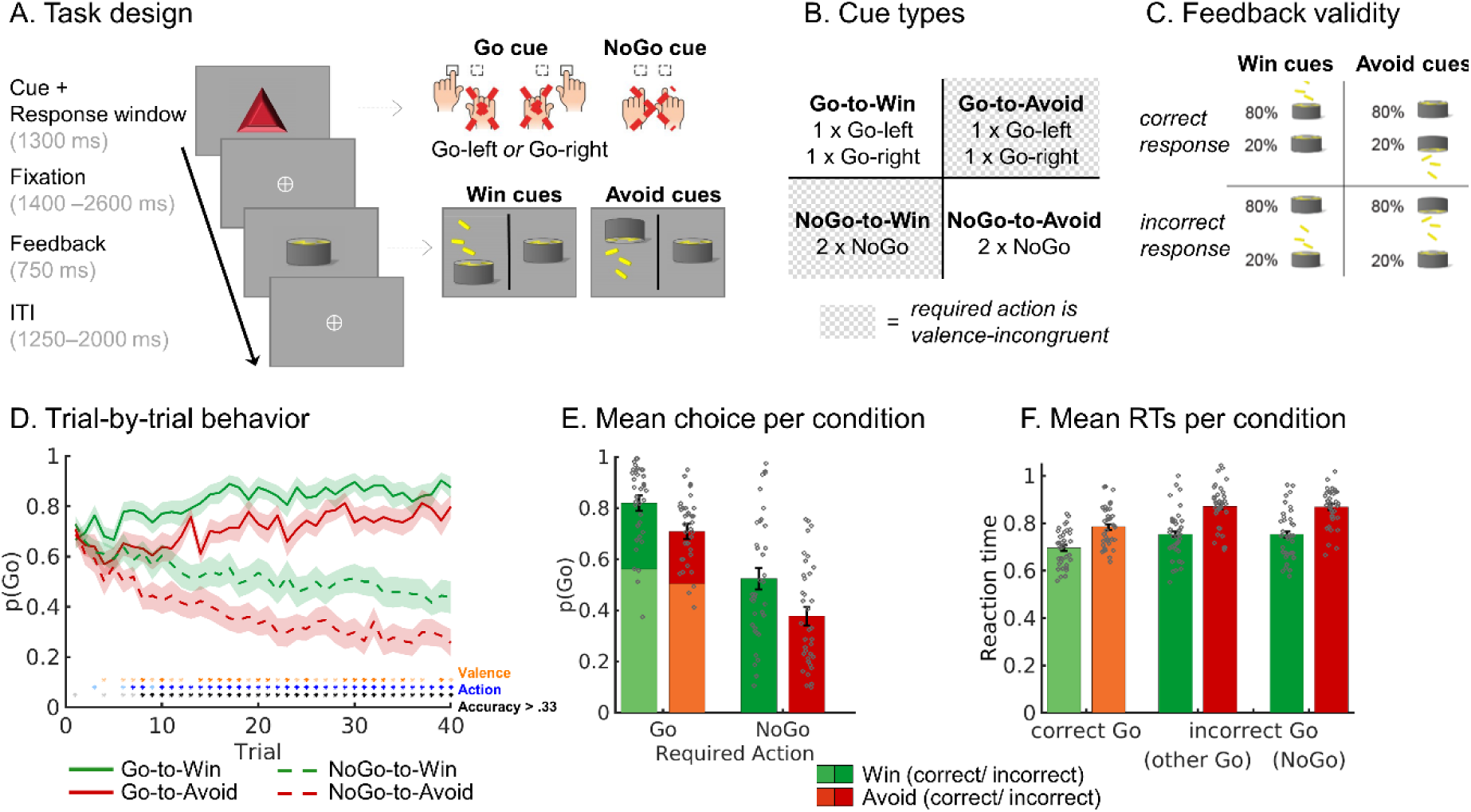
Motivational Go/NoGo learning task and performance. Motivational Go/NoGo learning task design and performance. (A) On each trial, a Win or Avoid cue appears; valence of the cue is not signaled but should be learned. Participants should respond during cue presentation. Response-dependent feedback follows after a jittered interval. Compared to a previous study with the same task (Swart et al. 2018), fixations between cue and feedback were jittered and 700–1900 ms longer and ITIs were 250 ms longer to allow us to disentangle cue- and outcome-related activity in the fMRI signal. Each cue has only one correct action (Go-left, Go-right, or NoGo), which is followed by the desired outcome 80% of the time. For Win cues, actions can lead to rewards or neutral outcomes; for Avoid cues, actions can lead to neutral outcomes or punishments. (B) There are eight different cues, orthogonalizing cue valence (Win versus Avoid) and required action (Go versus NoGo). The motivationally incongruent cues, for which the motivational action tendencies are incongruent with the instrumental requirements, are highlighted in gray. (C) Feedback is probabilistic: Correct actions to Win cues lead to rewards in 80% of cases, but neutral outcomes in 20% of cases. For Avoid cues, correct actions lead to neutral outcomes in 80% of cases, but punishments in 20% of cases. For incorrect actions, these probabilities are reversed. Rewards and punishments are depicted by money falling into/ out of a can. (D) Trial-by-trial proportion of Go actions (±SEM) for Go cues (solid lines) and NoGo cues (dashed lines). D) Trial-by-trial proportion of Go actions (±SEM) for Go cues (solid lines) and NoGo cues (dashed lines). Shadows indicate standard errors for per-condition-per-participant means across participants using the Cousineau-Morey method (Morey 2008). The motivational bias is defined as the tendency to make more Go actions to Win than Avoid cues (i.e., green lines are above red lines). Additionally, participants clearly learn whether to make a Go actions or not (solid lines go up, dashes lines go down). Orange asterisks below indicate trial-by-trial significance of motivational bias, blue asterisks indicate performance accuracy above chance (i.e., correct GoLeft, GoRight or NoGo response) (light color: p < .05 uncorrected; dark color: p < 0.0013; Bonferroni corrected for number of trials). (E) Mean (±SEM) proportion Go actions per cue condition (points are individual participants’ means). Proportion Go actions is higher for Go than NoGo cues, indicative of task learning, and higher for Win than Avoid cues, reflecting the influence of motivational biases on behavior. (F) Mean (±SEM) reaction times for correct and incorrect Go actions, the latter split up in whether the other Go response or the NoGo response would have been correct (points are individual participants’ means). Participants respond faster on correct than on incorrect Go actions and faster to Win than Avoid cues, reflecting the influence of motivational biases on behavior.

Participants received two button boxes in the scanner, one for each hand, and were instructed to use only one of the four keys on each button box. When participants accidentally pressed one of the three other buttons, the text “invalid response” appeared instead of an outcome. For analyses purposes, such invalid button presses were recoded into the valid button press of the respective hand, assuming that participants aimed for the correct key of the respective button box.

Before the actual task, participants underwent a practice session in which they were familiarized first with each condition separately (using practice stimuli) and then practiced all conditions together, using different cues from the actual experiment. They were informed about the probabilistic nature of feedback and that each cue features one optimal action. The actual task comprised 320 trials, split into three blocks of approximately ten minutes with short breaks between blocks. Participants performed the task twice, with a different set of cues, yielding 640 trials in total. Introducing a new set of cues allowed us to prevent ceiling effects in performance and investigate continuous learning throughout the task.

### Behavioral data analysis

For behavioral analyses, all trials with RTs before 1.3 seconds (i.e., cue offset) were treated as Go responses (in line with fMRI and EEG analyses). Button presses were treated as Go responses irrespective of whether the correct button or an incorrect button was pressed. We recoded 43 trials with invalid button presses (i.e., pressing a button that was not instructed as response button) into the correct response button of the respective hand (0.19% of trials; max. 14/640 per participant). For analyses of RTs, 12 trials with RTs smaller than 200 ms (0.08% of trials, max. 5/640 per participant) were excluded as such responses were unlikely to follow from cue-based action selection. Furthermore, 980 trials with RTs larger than 1300 ms (i.e., after cue offset, which was the instructed response time limit; 6.5% of trials, max. 139/640 per participant) were excluded from RT analyses as it was unclear whether such button presses were still intended as responses to the cue or were mere “action slips”. Results did not qualitatively change when including these trials.

We used mixed effects logistic regression (lme4 package in R) to analyze participants’ behavioral responses (Go vs. NoGo). We assessed a main effect of required action (Go/ NoGo), reflecting whether participants learned the task (i.e. showed more Go response to Go than NoGo cues), a main effect of cue valence (Win/ Avoid), reflecting the motivational bias (i.e. more Go response to Win than Avoid cues), and the interaction between valence and required action. The model written in Wilkinson notation was:

*response ∼ cueValence * requiredAction + (cueValence * requiredAction|participant)*

All variables were treated as factors, with sum-to-zero coding. To achieve a maximal random effects structure (Barr et al. 2013), we added random intercepts and random slopes of all three predictors for each participant and further allowed for random correlations between all predictors. The same predictors and random effects structure were used to analyze RTs, for which linear regression was used. *P*-values were computed using likelihood ratio tests (package afex in R). *P*-values smaller than α = 0.05 were considered statistically significant. In all plots, whiskers indicate standard errors, which were computed across participants based on the per-condition-per-participant means using the Cousineau-Morey method (Morey 2008).

### fMRI data acquisition

MRI data were acquired on a 3T Siemens Magnetom Prisma fit MRI scanner. In the scanner, participants’ heads were stabilized with foam pillows, and a strip of adhesive tape was applied to participants’ forehead to provide motion feedback and minimize head movement (Krause et al. 2019). After two localizer scans to position slices, functional images were collected using a whole-brain T2*-weighted sequence (68 axial-oblique slices, TR = 1400 ms, TE = 32 ms, voxel size 2.0 mm isotropic, interslice gap 0 mm, interleaved multiband slice acquisition with acceleration factor 4, FOV 210 mm, flip angle 75°, A/P phase encoding direction) and a 64-channel head coil. This sequence yielded a short TR at high spatial resolution, which allowed us to disentangle BOLD signal related to cue and outcome presentation. The sequence parameters were piloted to find a sequence with minimal signal loss in the striatum. The first seven volumes of each run were automatically discarded.

After task completion, when the EEG cap was removed, an anatomical image was collected using a T1-weighted MP-RAGE sequence (192 sagittal slices per slab, GRAPPA acceleration factor = 2, TI = 1100 ms, TR = 2300 ms, TE = 3.03 ms, FOV 256 mm, voxel size 1.0 mm isotropic, flip angle 8°) for registration and a gradient fieldmap (GRE; TR = 614 ms, TE1 = 4.92 ms, voxel size 2.4 mm isotropic, flip angle 60°) for distortion correction. For one participant, no fieldmap was collected due to time constraints. At the end of each session, an additional diffusion tensor imaging (DTI) data collection took place; results will be reported elsewhere.

### fMRI preprocessing

fMRI data for each of the six blocks per participants were preprocessed using FSL 6.0.0 (Smith et al. 2004). Functional images were cleaned for non-brain tissue (BET; Smith, 2002), segmented, motion-corrected (MC-FLIRT; Jenkinson, Bannister, Brady, & Smith, 2002), and smoothed (FWHM 3 mm). Field maps were used for B0 unwarping and distortion correction in orbitofrontal areas. We used ICA-AROMA (Pruim et al. 2015) to automatically detect and reject independent components in the data that were associated with head motion (non-aggressive denoising option).

To prevent empty regressors on a block level, we concatenated blocks and performed a single first-level GLM per participant. For this purpose, we registered the volumes of all blocks to the middle image (the default registration option in FSL) of the first block of each participant (using MCFLIRT) and then merged files. The first and last 20 seconds of each block did not contain any trial events, such that, when modelling trial events, no carry-over effects from one block to another could occur.

After concatenation and co-registration of the EPI, we performed high-pass filtering with a cutoff of 100 s, and pre-whitening. We then computed the co-registration matrices of EPI images to high-resolution anatomical images (linearly with FLIRT using Boundary-Based Registration) and to MNI152 2mm isotropic standard space (non-linearly with FNIRT using 12 DOF and 10 mm warp resolution; Andersson, Jenkinson, & Smith, 2007).

### ROI selection

We used masks of selected ROIs for three purposes: fMRI GLMs with small volume correction, BOLD-RT correlations, and fMRI-informed EEG analyses. For GLMs with small-volume correction, we used anatomical masks of striatum and ACC/pre-SMA based on our a-priori hypothesis and previous literature. Anatomical masks were based on the Harvard-Oxford atlas, thresholded at 10%. For BOLD-RT correlations and fMRI-informed EEG analyses, we used conjunctions of anatomical masks and relevant functional contrasts. These are follow-up analyses to results observed in the fMRI GLMs and test independent hypotheses. The well-established rationale for using these conjunctive constraints (O’Reilly et al. 2012) is that the anatomical constraints ensure that all voxels are from the same anatomical region, while the functional constraints ensure that voxels reflect the signal of interest. These masks were obtained by thresholding the z-map of the relevant functional contrast at z > 3.1 (cluster-forming threshold) and then combining it with the anatomical mask using the logical AND operation.

We obtained the following anatomical masks by combining sub-masks of the Harvard-Oxford atlas: striatum (bilateral caudate, putamen, and nucleus accumbens), midfrontal cortex (anterior division of the cingulate gyrus and juxtapositional lobule cortex—formerly known as supplementary motor cortex), ACC (anterior division of the cingulate gyrus), left and right motor cortex (precentral and postcentral gyrus), vmPFC (frontal pole, frontal medial cortex, and paracingulate gyrus).

Masks were either used in the group-level GLM for small-volume correction or back-transformed to participants’ native space to extract either parameter estimates from the GLM or the raw BOLD signal time series. All masks are displayed in Supplementary Material S02.

### fMRI analysis

We fitted a first-level GLM to the data of each participant using the fixed-effects model in FSL FEAT. The four task regressors of interest were the four conditions resulting from crossing cue valence (Win/Avoid) and performed action (Go/NoGo irrespective of Left vs. Right Go), all modeled at cue onset. We also included five regressors of no interest: two at cue onset, namely response side (Go left = +1, Go right = -1, NoGo = 0) and errors (i.e. participants chose the incorrect action), and three at outcome onset to control for outcome-related activity, namely outcome onset (intercept of 1 for every outcome), outcome valence (reward = +1, punishment = -1, neutral = 0), and invalid trials (invalid buttons pressed and thus not feedback given). We further added the following nuisance regressors (separate regressors for each block): intercept, six realignment parameters from motion correction, mean cerebrospinal fluid (CSF) signal, mean out-of-brain (OOB) signal, and separate spike regressors to model out each volume on which relative scan-to-scan displacement was more than 2 mm (occurred in 10 participants; in those participants: *M* = 7.40, range 1 – 29). Task regressors were convolved with a double-gamma haemodynamic response function (HRF) and high-pass filtered at 100 s. The full model is also displayed in Supplementary Material S03.

We hypothesized that a main effect of valence in the striatum would be attenuated under motivational conflict. Thus, the positive BOLD response to a Win cue would be reduced when having to make a NoGo response for Win, and the negative BOLD response to an Avoid cue would be less negative for a Go response for Avoid (Figure 3A). This hypothesized conflict mediated attenuation would thus manifest itself as a (weaker) main effect of action in the presence of a (stronger) main effect of valence. Taken together, we predicted that striatal BOLD signal would exhibit both a main effect of valence and a main effect of action. Indeed, previous literature using a simpler version of this task (Guitart-Masip et al. 2011; Guitart-Masip, Chowdhury, et al. 2012; Guitart-Masip, Huys, et al. 2012) has reported a main effect of action in the striatum. In addition to testing the main effects of valence and action, we also computed a congruency contrast. Finally, left vs. right Go responses were contrasted to identify lateralized motor cortex activation as a quality control check. Regressors were specified across both correct and incorrect trials; we report results for regressors on the correct trials only (in line with our EEG analysis approach) in Supplementary Material S3.

First-level participant-specific contrasts were fitted in native space. Co-registration and re-slicing was then applied on each participant’s contrast maps, which were combined at the group-level (using FSL FLAME for mixed effects models; Woolrich et al., 2009; Woolrich, Behrens, Beckmann, Jenkinson, & Smith, 2004) with a cluster-forming threshold of *z* > 3.1 and cluster-level error control at α < .05 (i.e. two one-sided tests with α < .025).

Given our specific hypotheses about the striatum and midfrontal cortex, we additionally tested contrasts using a small-volume correction: For the Valence and Action contrasts, we used an anatomical mask of the striatum, and for the Congruency contrast (i.e. NoGo2Win and Go2Avoid minus Go2Win and NoGo2Avoid), we used a mask of the entire midfrontal cortex (ACC and pre-SMA). While theoretical models predict differences in BOLD signal in ACC, empirical fMRI studies have actually found correlates in more dorsal regions, specifically pre-SMA (Botvinick et al. 2004). Indeed, source reconstruction of midfrontal theta suggested a rather superficial source in pre-SMA (Cohen and Ridderinkhof 2013) and a simultaneous EEG-fMRI study found choice conflict related to pre-SMA BOLD (Frank et al. 2015). Thus, the anatomical mask comprised the entire midfrontal cortex.

### BOLD – behavior correlations

To assess whether regions that encoded cue valence, i.e. information driving motivational biases, predicted reaction times, we performed regression analyses of the trial-by-trial BOLD signal in relevant regions on RTs (for trials where participants made a Go response). The main regions we found to encode cue valence were vmPFC (positively) and ACC (negatively). We computed conjunctions of anatomical vmPFC and ACC masks with the cue valence contrast and back-transformed the resultant masks to each participant’s native space. We then extracted the first eigenvariate of the signal in both ROIs, returning one summary measure of BOLD signal in that ROI per volume. The volume-by-volume signal in each ROI was then highpass-filtered at 128 s and nuisance regressors (6 realignment parameters, CSF, OOB, single volumes with strong motion, same as in the fMRI GLM) were regressed out. Afterwards, the signal was upsampled by factor 10, epoched into trials of 8 s duration (Hauser et al. 2015), and a separate HRF was fitted for each trial (i.e. 57 upsampled datapoints).

We then tested whether BOLD signal in those ROIs correlated with reaction times on a trial-by-trial basis. In order to account for overall differences between trials with Win cues and trials with Avoid cues, we standardized reaction times and BOLD signal separately for Win trials and Avoid trials within each participant such that differences between cue valence conditions were removed. We computed correlations between trial-by-trial reaction times and BOLD signal HRF amplitude for each participant, applied Fisher-*z* transformations to correlations to make them normally distributed, and then tested with a one-sample *t*-test whether correlations were significantly different from zero at a group level.

### EEG data acquisition and preprocessing

EEG data were acquired using 64 channels (BrainCap-MR-3-0 64Ch-Standard; Easycap GmbH; Herrsching, Germany; international 10-20 layout, reference electrode at FCz) at a sampling rate of a 1,000 Hz using MRI-compatible EEG amplifiers (BrainAmp MR plus; Brain Products GmbH, Gilching, Germany). Additional channels for electrocardiogram, heart rate, and respiration were used recorded for MR artifact correction. Recordings were performed with Brain Vision Recorder Software (Brain Products). EEG amplifiers were placed behind the scanner, and cables were attached to the cap once participants were positioned in the scanner. Cables were fixated with sand-filled pillows to reduce artifacts induced through cable movement in the magnetic field. During functional scans, the MR scanner helium pump was switched off to reduce EEG artifacts. A Polhemus FASTRAK device was used to record the exact location of each EEG electrode on the participant’s head relative to three fiducial points. For four participants, no Polhemus data were recorded due to time constraints and technical errors; for these participants, the average channel positions of the remaining 32 participants were used.

EEG data were cleaned from MR scanner and cardioballistic artifacts using BrainVisionAnalyzer (Allen, Josephs, & Turner, 2000). Pre-processing was performed in Fieldtrip (Oostenveld et al. 2011) in MATLAB 2017b by rejecting channels with high residual MR noise (mean 4.8 channels per participant, range 0–13), epoching trials (-1750–2800 ms relative to cue onset, total duration of 4550 ms), re-referencing the channels to their grand average and recovering the reference as channel FCz, band-pass filtering the data in the 0.5– 15 Hz range using a two-pass 4^th^ order Butterworth IIR filter (Fieldtrip default), and finally linear baseline correction based on the 200 ms prior to cue onset. The low-pass filter cut-off of 15 Hz allowed us to dissociate theta from its adjacent bands while filtering out residual high-frequency MR noise. We used ICAs to visually identify and reject independent components related to eye blinks, saccades, head motion, and residual MR artifacts (mean 12.94 components per participant, range 8–19), and afterwards manually rejected trials that were still contaminated by noise (mean 29.6 trials per participant, range 2–112). Finally, we computed a Laplacian filter with the spherical spline method to remove global noise (using the exact electrode positions obtained with the Polhemus FASTRAK), which we also used to interpolate previously rejected channels. This filter attenuates more global signals (deep sources) and noise (heart-beat and muscle artifacts) while accentuating more local effects (superficial sources).

For response-locked analyses, we re-epoched Go actions trials, time-locked to the time of response (RT). For NoGo response trials, we re-epoched the data time-locked to the average RTs (for each participant) of Go actions for that cue valence, as a proxy for ‘latent RTs’ on these trials.

### EEG TF decomposition

Time-frequency decomposition was performed using Hanning tapers between 1–15 Hz in steps of 1 Hz, every 25 ms with 400 ms time windows. We first zero-padded trials to a length of 8 sec. and then performed time-frequency decomposition in steps of 1 Hz by multiplying the Fourier transform of the trial with the Fourier transform of a Hanning taper of 400 ms width, centered around the time point of interest. This procedure results in an effective resolution of Hz (Rayleigh frequency), interpolated in 1 Hz steps, which is more robust to the exact choice of frequency bins. Given that all pre-processing was performed on data epoched into trials, we aimed to exclude the possibility of slow drifts in power over the time course of the experiment. We thus performed baseline correction by fitting a linear model across trials for each channel/frequency combination. This model included trial number as a regressor and the average power in the last 50 ms before cue onset as outcome. The power predicted by this model was then removed from single-trial data. Note that in absence of any drift, this approach amounts to correcting all trials by the grand-mean across trials per frequency in the selected baseline window per participant. Next, we averaged over trials within each condition spanned by of valence (Win/ Avoid) and action (Go/ NoGo; correct trials only). Finally, power was converted to decibel for all analyses to ensure that data across frequencies, time points, electrodes, and participants were on the same scale.

In line with (Swart et al. 2018), we restricted our analyses to correct trials, i.e. trials where required and performed action matched. We assumed that on correct incongruent (Go2Avoid and NoGo2Win) trials, participants successfully detected and resolved conflict, while no such processes were required on congruent (Go2Win and NoGo2Avoid) trials. Although the same processes might be initiated on incorrect trials, though unsuccessfully, these trials are potentially confounded by error-related synchronization in the theta range (Cavanagh, Zambrano-Vazquez, et al. 2012), which makes the interpretation of any effects in the theta range less straightforward. To be consistent with fMRI results, we report results across both correct and incorrect trials in Supplementary Materials S06 and S10.

### EEG data analysis

All analyses were performed on the average signal of the a-priori selected channels Fz, FCz, and Cz based on previous findings (Swart et al. 2018). We performed non-parametric cluster-based permutation tests (Maris and Oostenveld 2007) as implemented in Fieldtrip for the selected electrodes in the theta range (4–8 Hz) during cue presentation (0–1300 ms). This procedure is suited to reject the null hypothesis of exchangeability of two experimental conditions, but not suited to exactly determine when or where differences occur (Sassenhagen and Draschkow 2019). Our interpretations of when and where conditions differed in power are thus based on visual inspection of the signal time courses.

Given our a-priori hypothesis of midfrontal theta power reflecting conflict, we performed the test contrasting bias-incongruent than bias-congruent actions to the theta range (4–8Hz). Furthermore, since visual inspection of the condition-specific time courses of theta power suggested major differences in theta power between Go and NoGo responses, we additionally performed an exploratory test contrasting Go and NoGo responses. Given its exploratory nature, this test was performed on broadband power (1–15 Hz).

### fMRI-informed EEG analysis

The sluggish nature of the BOLD signal makes it difficult to determine when exactly different brain regions become active. In contrast, EEG provides much higher temporal resolution. Identifying distinct EEG correlates of the BOLD signal in different regions could thus reveal when these regions become active (Hauser et al. 2015). Furthermore, using the BOLD signal from different regions in a multiple linear regression allows to control for variance that is shared among regions (e.g. changes in global signal; variance due to task regressors) and test which region is the best unique predictor of a certain EEG signal. In such an analysis, any correlation between EEG and BOLD signal from a certain region reflects an association above and beyond those induced by task conditions.

To link BOLD signal from distinct regions to time-frequency power, we applied the same approach as for BOLD-RT correlation analyses and fitted a trial-by-trial hemodynamic response function (HRF) to the BOLD signal in selected ROIs. We then used the trial-by-trial HRF amplitudes to predict TF power at each time-frequency-channel bin (Hauser et al. 2015). For BOLD signal extraction, we specified six ROIs using a combination of functional and anatomical constraints based on our fMRI GLM results: vmPFC (valence contrast), ACC (valence and action contrast), left and right motor cortex (response side contrast, which captured lateralized motor activity better than the action contrast), and striatum (action contrast). For ACC, we obtained two separate masks (valence and action contrast), which strongly overlapped. Analyses included only one of those masks at a time; conclusions were identical with either mask. The resultant masks were back-transformed to each participant’s native space and the first eigenvariate of the signal in each ROI was extracted, returning one summary measure of BOLD signal in that ROI per volume. We applied the same high-pass filter, nuisance regression, upsampling procedure, and trial-by-trial HRF estimation as in BOLD-RT correlation analyses. The trial-wise HRF amplitude estimates for each ROI were used as regressors in a multiple linear regression to predict the TF power for each 3D time-frequency-channel bin across trials, resulting in a 3D map of regression weights (*b*-map) for each ROI. In these regressions, we also added behavioral regressors for the main effects of required action, valence, and their interaction as covariates of no interest to account for task-related variance in EEG power. In all analyses, predictors and outcomes were demeaned so that any intercept was zero. Finally, participants’ *b*-maps were Fisher-*z* transformed (which makes the sampling distribution of correlation coefficients approximately normal and allows to combine them across participants).

Finally, to test whether BOLD signal in certain ROIs uniquely predicted variance in TF power, we performed cluster-based one-sample permutation *t*-tests across participants (Hunt et al. 2013). We performed these tests on the mean regression weights of the channels that exhibited condition differences in the EEG-only analyses (FCz and Cz; Fz was dropped because it did not show significant power differences between conditions) in the range of 0– 1300 ms (i.e. duration of cue presentation), 1–15 Hz. We first obtained a null distribution of maximal cluster mass statistics from 10000 permutations. In each permutation, the sign of the *b*-map of a random subset of participants was flipped. Then, a separate *t*-test for each time-frequency bin (bins of 25 ms, 1 Hz) across participants was computed. The resulting *t*-map was thresholded at |*t*| > 2, from which we computed the maximal cluster mass statistic (i.e. sum of all *t*-values) of any cluster (i.e. adjacent voxels above threshold). We next computed a *t*-map for the real data, from which we identified the cluster with largest cluster mass statistic. The corresponding *p*-value was computed as the number of permutations with larger maximal cluster mass than the maximal cluster mass of the real data, and considered significant for a p-value of < .05.

## Results

Thirty-six healthy participants performed a motivational Go/NoGo learning task. In this task, they needed to learn by trial and error which response (left Go/ right Go/NoGo) to make in order to gain rewards (“Win” cues) or avoid losses (“Avoid” cues); see Fig 1. We simultaneously measured EEG and fMRI while participants performed this task.

### Task performance

Participants successfully learned the task, as they performed significantly more Go actions to Go cues than NoGo cues (Required action: χ^2^(1) = 32.01, *p* < .001; Fig. 1D-E). Furthermore, participants showed a motivational bias, as they performed more Go actions for Win cues than Avoid cues (Valence: χ^2^(1) = 23.70, *p* < .001). The interaction of Required action x Valence was not significant (χ^2^(1) = 0.20, *p* = .658), suggesting that motivational biases occurred similarly for Go and NoGo cues.

When making a (Go) action, participants responded faster to Go cues (averaged over correct and incorrect responses) than to NoGo cues (where responses were by definition incorrect) (Required action: χ^2^(1) = 25.64, *p* < .001). Furthermore, a motivational bias was present also in RTs, with significantly faster actions to Win than Avoid cues (Valence (χ^2^(1) = 44.58, *p* < .001). Again, the interaction was not significant (χ^2^(1) = 1.51, *p* = .219; see Fig. 1F).

### fMRI

#### Valence

We hypothesized higher BOLD signal in the striatum for Win compared with Avoid cues (Fig. 2A). There were no significant clusters in the striatum in a whole-brain corrected analysis. When restricting our analyses to an anatomical mask of the striatum, there were two significant clusters (for a complete list of all significant clusters and *p*-values, see Supplementary Material S04): BOLD signal in left posterior putamen was significantly higher for Win than Avoid cues (Fig. 2E; no longer significant when excluding the five participants that were excluded from the final EEG-fMRI analysis; Supplementary Material S01). In contrast, BOLD signal in the bilateral medial caudate nucleus was, surprisingly, higher for Avoid than Win cues (Fig. 2F). While the effect in left putamen appeared robust over the time course of the task, the effect in medial caudate was strongest at the beginning of the task, but then disappeared towards the end of the task (see Supplementary Material S05). At a whole-brain level cluster correction, the largest cluster of higher BOLD signal for Win than Avoid cues was observed in the ventromedial prefrontal cortex (vmPFC). Positive valence coding was also observed in bilateral dorsolateral prefrontal cortex (dlPFC), bilateral ventrolateral prefrontal cortex (vlPFC), posterior cingulate cortex (PCC), bilateral amygdala, and bilateral hippocampus (Fig. 2A). Conversely, BOLD was higher for Avoid cues in dorsal ACC and bilateral insula. Repeating analyses on correct trials only yielded identical results in the whole-brain analyses, while effects posterior putamen and medial caudate with small-volume correction were not significant anymore (see Supplementary Material S06).

**Fig. 2.**
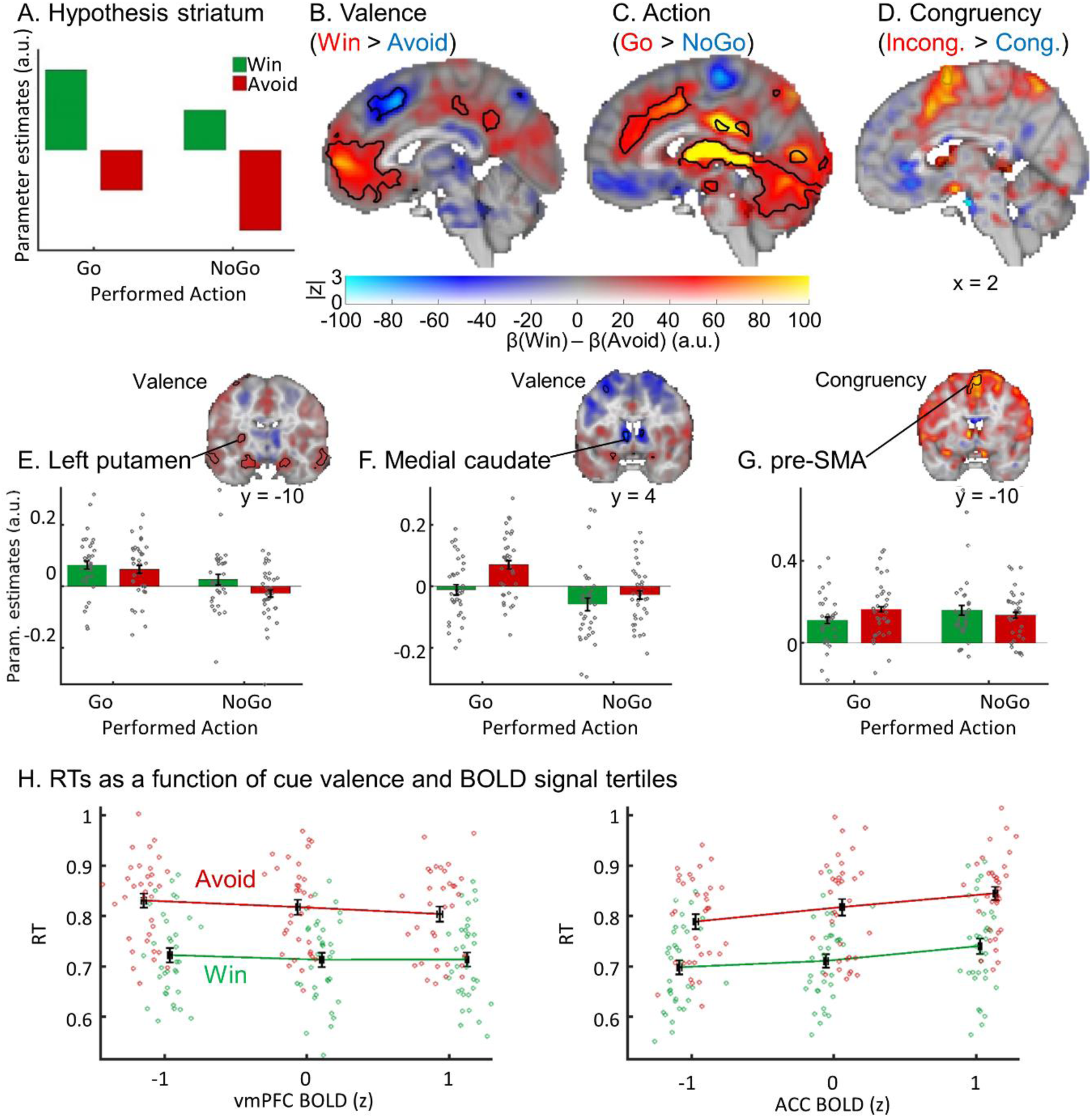
fMRI results. BOLD signal as a function of cue valence, performed action, and congruency. (A) We hypothesized striatal BOLD to encode cue valence (main effect of valence), with an attenuation of this valence signal when actions incongruent to the bias-triggered actions were performed (main effect of action). (B) BOLD signal was significantly higher for *Win* compared to *Avoid* cues in ventromedial prefrontal cortex (vmPFC; whole brain corrected) and left putamen (small-volume corrected), but higher for *Avoid* compared to *Win* cues in ACC and medial caudate (small-volume corrected). (C) BOLD signal was significantly higher for *Go* compared with *NoGo* responses in the entire striatum as well as ACC, thalamus, and cerebellum (all whole-brain corrected). (D) BOLD signal was significantly higher for bias-incongruent actions than bias-congruent actions in pre-SMA (small-volume corrected). B-D. BOLD effects displayed using a dual-coding visualization with color indicating the parameter estimates and opacity the associated *z*-statistics. Contours indicate statistically significant clusters (p < .05), either small-volume corrected (striatal and SMA contours explicitly linked to a bar plot) or whole-brain corrected (all other contours). (E-G) Mean beta weights per task condition (x-axis) per participant (individual grey dots) in significant clusters in left putamen, medial caudate and pre-SMA (significant in small-volume correction). (E) Left posterior putamen encoded valence positively (higher BOLD for Win than Avoid cues), but was dominated by an encoding of the performed action (higher BOLD for Go than NoGo responses). (F) Medial caudate encoded valence negatively (higher BOLD for Avoid than Win cues), again predominantly showing a main effect of action. (G) BOLD signal in pre-SMA was higher for bias-incongruent than bias-congruent actions (small-volume corrected). (H) Reaction times (RTs) as a function of cue valence and BOLD signal tertiles (z-standardized) per participant (individual dots; x-location relative to all other participants). RTs were significantly predicted by BOLD signal in vmPFC (positively) as well as by BOLD signal in ACC and striatum (negatively). BOLD-RT correlations were independent of cue valence. Lines connect the means of RT tertiles. Error bars (±SEM) for both BOLD (vertically) and RTs (horizontally) are very narrow.

#### RTs

We reasoned that any region translating cue valence into motivational biases in behavior should also predict the speed-up of RTs for Win compared with Avoid cues. Our finding that vmPFC and ACC BOLD reflected cue valence is in line with a wealth of previous literature (Haber and Knutson 2010; Bartra et al. 2013), raising the question whether signals from these two regions impact action selection in a way that gives rise to motivational biases. We reasoned that if such signals have a causal effect on behavior, they should predict RT differences, i.e. both overall RT differences between Win and Avoid cues, but also RT differences within each valence condition. To assess whether BOLD signal in vmPFC and ACC indeed related to the speed of selected actions, we computed correlations of reaction times with the trial-by-trial deconvolved BOLD signal in the identified vmPFC and ACC clusters. To account for overall differences between Win and Avoid cues, we standardized RTs and BOLD signal separately for Win and Avoid cues, removing the overall difference between both conditions. There was a strong negative association for vmPFC, *t*(33) = -4.11, *p* < 0.001, *d* = - 0.71, with higher vmPFC BOLD predicting faster reaction times, and a strong positive association for ACC, *t*(33) = 7.83, *p* < .001, *d* = 1.34, with higher ACC BOLD predicting slower reaction times (Fig. 2H). These results are consistent with vmPFC and ACC, both signaling cue valence, influencing the speed of downstream actions and thus contributing to motivational biases in reaction times. However, we cannot infer a causal role of these regions from the observed correlation.

#### Action

We further hypothesized that the valence signal in the striatum would be modulated by the congruency between valence and action, such that increased striatal BOLD signal for Win cues would be dampened when (bias-incongruent) NoGo responses were required, while decreased striatal activity for Avoid cues should be elevated when (bias-incongruent) Go actions were required (Fig. 2A). This interaction effect between valence and congruency is equivalent to a main effect of action.

Striatal BOLD (bilateral caudate nucleus, putamen, and nucleus accumbens) was indeed significantly higher for Go than NoGo responses (Fig. 2C). This effect was absent on the first block, but strongly emerged over time (see Supplementary Material S05). However, in absence of a clear effect of valence on striatal BOLD, this main effect of action might not reflect an attenuation of valence signaling. Rather, the striatum appears to predominantly encode action itself–even in the left posterior putamen, which significantly encoded valence (Fig. 2E). This finding replicates previous studies using a different version of the Motivational Go/NoGo task (Guitart-Masip et al. 2011; Guitart-Masip, Chowdhury, et al. 2012; Guitart-Masip, Huys, et al. 2012). Other regions that responded more strongly to Go versus NoGo responses included the ACC, thalamus and bilateral cerebellum. For a complete list of significant clusters, see Supplementary Material S04. Repeating analyses on correct trials only yielded identical results (see Supplementary Material S06).

#### Valence x Action interaction (Congruency)

Based on prior work, we expected increased BOLD for bias-incongruent compared to bias-congruent actions in midfrontal cortex, the putative cortical source of midfrontal theta oscillations. At a whole-brain cluster level significance correction, there were no clusters in which BOLD signal differed between bias-congruent and -incongruent actions. When restricting the analysis to a mask comprising ACC and pre-SMA, there was a cluster in pre-SMA (Fig. 2D, G; not significant when excluding the five more participants excluded in the final EEG-fMRI analysis; see Supplementary Material S01; not significant in the GLM on correct trials only, see Supplementary Material S06). This finding is in line with source reconstruction studies of midfrontal theta (Cohen and Ridderinkhof 2013) and EEG-fMRI findings observing choice conflict-related activity in pre-SMA (Frank et al. 2015). The effect of conflict in pre-SMA was robust over time (see Supplementary Material S05).

### EEG

#### Action x Valence interaction (Congruency)

We used a non-parametric, cluster-based permutation test to test whether time-frequency (TF) power in the theta band (4–8 Hz) was significantly higher on motivational incongruent (Go2Avoid, NoGo2Win) than congruent (Go2Win, NoGo2Avoid) trials over midfrontal channels (Fz, FCz, Cz). We indeed found that theta power was higher on incongruent than congruent trials (*p* = .023), most strongly around 175–325 ms after cue onset. However, this difference occurred markedly earlier than in our previous study (450–650 ms) (Swart et al. 2018) and visual inspection of the time-frequency plot showed that the peak of this cluster was located rather in the alpha band (8–12 Hz), leaking into the upper theta range (Fig. 3E-F), and restricted to an early, transient increase in alpha power over midfrontal channels. This congruency effect was not present in evoked activity (ERPs; see Supplementary Material S07) and indeed remained unaltered after the subtraction of evoked activity (see Supplementary Material S08). Furthermore, this change in alpha power occurred selectively on incongruent trials on which participants made a correct response, rather than nonspecifically on all incongruent or on all correct trials (see Supplementary Material S09). In other words, power increased selectively when biases were successfully suppressed (Swart et al. 2018). In sum, we found an electrophysiological correlate of conflict, which was however different in time and frequency range from our previous finding (Swart et al. 2018).

**Fig. 3.**
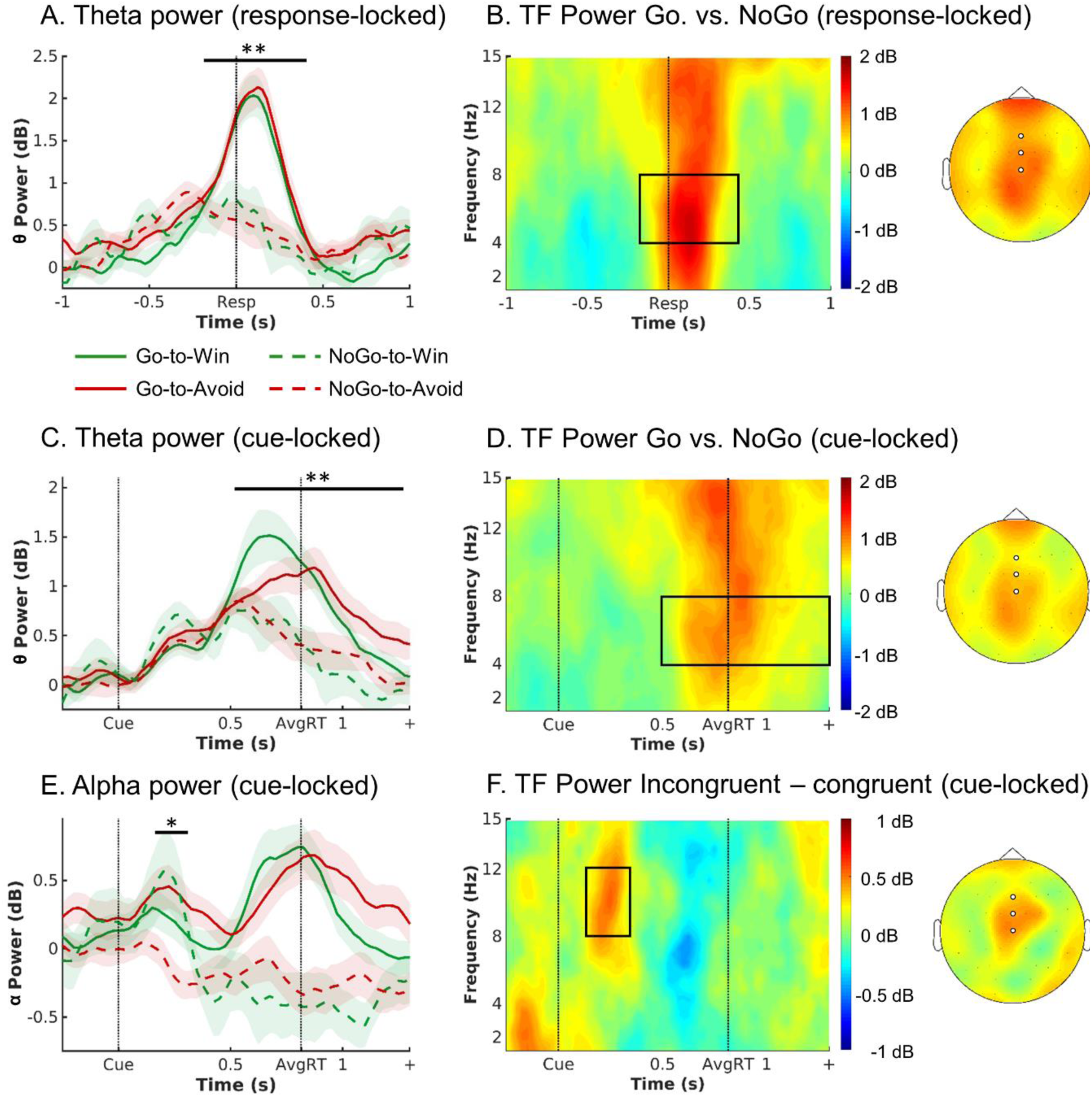
EEG results. EEG time-frequency power as a function of cue valence and action. (A) Response-locked within trial time course of average theta power (4–8 Hz) over midfrontal electrodes (Fz/ FCz/ Cz) per cue condition (correct-trials only). Theta increased in all conditions relative to pre-cue levels, but to a higher level for Go than NoGo trials. There were no differences in theta peak height or latency between Go2Win and Go2Avoid trials. (B) Left: Response-locked time-frequency power over midfrontal electrodes for Go minus NoGo trials. Go trials featured higher broadband TF power than NoGo trials. The broadband power increase for Go compared to NoGo trials is strongest in the theta range. Right: Topoplot for Go minus NoGo trials. The difference is strongest at FZ and FCz electrodes. (C-D) Cue-locked within trial time course and time-frequency power. Theta increased in all conditions relative to pre-cue levels, but to a higher level for Go than NoGo trials, with earlier peaks for Go2Win than Go2Avoid trials. (E) Trial time course of average alpha power (8–13 Hz) over midfrontal electrodes per cue condition (correct trials only; cue-locked). Alpha power transiently increases for both incongruent conditions in an early time window (around 175–325 ms). (F) Left: Time-frequency plot displaying that the transient power increase was focused on the alpha band (8 – 13 Hz), leaking into the upper theta band. Right: Topoplot of alpha power displaying that this incongruency effect was restricted to midfrontal electrodes (highlighted by white disks). * *p* < 0.05. ** *p* < 0.01. Shaded errorbars indicate (±SEM). Box in TF plots indicates the time frequency window where *t*-values > 2.

#### Action and Valence main effects

We observed a modulation of time-frequency power by valence-action congruency in the alpha range shortly after cue onset (around 175–325 ms). We next performed permutation tests to explore whether time-frequency power was further modulated at any later time point by the individual task factors rather than their interaction (i.e., action or valence). Broadband power (1–15 Hz) was significantly higher on trials with Go actions than NoGo responses (cue-locked: *p* = .002; response-locked: *p* = .002): This difference between Go and NoGo responses occurred as a broadband-signal from 1–15 Hz, but peaked in the beta band (cue-locked) and theta band (response-locked; Fig. 3B and D). The topographies exhibited a bimodal distribution with peaks both at frontopolar (FPz) and central (FCz, Cz, CPz) electrodes (Fig. 3B and 3D). As visual inspection of Fig. 3C shows, theta power increased in all conditions until 500 ms post cue onset and then bifurcated depending on the action: For NoGo responses, power decreased, while for Go actions, power kept rising and peaked at the time of the response. This resulted in higher broadband power for Go versus NoGo responses for about 550–1300 ms after cue onset (see Fig. 3C-D; around -200–400 ms when response-locked, see Fig. 3A-B; same held in analyses across both correct and incorrect trials, see Supplementary Material S10). When looking at the cue-locked signal, the signal peaked earlier and higher for Go actions to Win than to Avoid cues, in line with faster reaction times on Go2Win than Go2Avoid trials. When testing for differences in broadband power between Win cues and Avoid cues, broadband power was indeed higher for Avoid than Win cues around 825–1300 ms cue-locked (*p* = .002). When comparing power time courses for Go responses to Win and to Avoid cues selectively in the theta range, theta power was higher for Win than Avoid cues around 550–700 ms, *p* = 0.028, but then higher for Avoid than Win cues around 925–1300 ms, *p* = .002. This difference in latency and peak height of the ramping signal was not present in the response-locked signal, and the respective test of Win vs. Avoid cues not significant (*p* = .110; Fig. 3A and Supplementary Material S11).

The occurrence of this signal close to response execution across a broadband frequency range raised the question whether it reflects (a) a decision process (incorporating decision parameters like cue valence and action values) or rather (b) a generic motor signal occurring for any manual response, or even (c) a signal artifact of head motion in the scanner. Regarding the latter, a number of control analyses indicated that the theta increase was likely not reducible to a motor artifact: first, results remained unchanged when accounting for fMRI-realignment parameters (reflecting head motion) using a linear regression approach (following Fellner et al. 2016; see Supplementary Material S12), and second, relative to the pre-trial baseline, increases were clearly focused on the theta band (see Supplementary Material S13), started already 300 ms after cue onset, and even occurred on NoGo trials (i.e. where no overt response was executed). Furthermore, the theta signal was unlikely to reflect a generic motor response, as it was modulated by task demands: Theta (but not broadband) power was higher for left, non-dominant hand compared to the right, dominant hand (in line with reaction time findings; see Supplementary Material S14), and higher for correct than incorrect responses (see Supplementary Material S11). Taken together, we cannot exclude the possibility that motor execution processes or signal artifacts contributed to the observed differences in theta power; however, differences in theta are likely not reducible to such processes, and do at least in part reflect pre-response, decision-related processes.

Next, if midfrontal theta power reflects a decision process, we asked whether this signal bore resemble to evidence accumulation processes described in perceptual decision-making before (Gold and Shadlen 2007; O’Connell et al. 2012). In such processes, a response is elicited once an accumulation signal reaches a certain threshold. This observation was particularly the case for the theta band (Fig. 3C) rather than any other band (see alpha band in Fig. 3E). Three further tests corroborated this interpretation: First, the peak of the ramping theta signal in the cue-locked data predicted reaction times within participants (see Supplementary Material S11), while differences in peak height and latency were absent in the response-locked data. Thus, faster accumulation of evidence in favor of a ‘Go’ response for ‘Win’ cues could be the driving mechanism for a motivational bias. This change in accumulation rate would then be putatively driven by a neural region encoding cue valence, such as vmPFC or ACC. Second, the “threshold” that theta needed to reach to elicit a response was higher for responses of the left (non-dominant) than the right (dominant) hand (in line with reaction times findings; Supplementary Material S14), putatively reflecting response competition in which the dominant hand needs to be overruled by raising response thresholds. Third, peak theta was lower for incorrect than correct responses (Supplementary Material S11), consistent with the idea that incorrect responses might (sometimes) reflect premature responding due to random fluctuations in response thresholds (O’Connell et al. 2012). These three observations are consistent with an interpretation of the ramping theta signal as reflecting an evidence accumulation process for Go-related evidence.

### fMRI-informed EEG analysis

Given that we observed action encoding in both BOLD (ACC, striatum) and midfrontal EEG power (theta), we next tested when differences in activity in these regions occurred by correlating the BOLD signal from those regions with midfrontal EEG time-frequency power. We extracted trial-by-trial BOLD signal from clusters encoding valence (vmPFC, ACC), action (striatum, ACC) and response hand (left and right motor cortex) and used these signals in a multiple linear regression to predict midfrontal time-frequency power.

First, trial-by-trial analyses revealed that striatal BOLD correlated positively with theta power around the time of response, over and above task and condition effects (*p* = .037 cluster-corrected; Fig. 4A). Conversely, and perhaps surprisingly, there was no such correlation between midfrontal theta power and ACC BOLD (mask based on action contrast: *p* = .268 cluster-corrected; mask based on valence contrast: *p* = .592). Supplementary analyses yielded correlations between motor cortex BOLD and midfrontal beta power (Jurkiewicz et al. 2006; Ritter et al. 2009), corroborating the overall ability of our approach in detecting well established BOLD-EEG associations (Supplementary Material S15). Note that because the design matrix included all ROI timeseries as well as task regressors, these correlations only reflect variance uniquely explained by a specific ROI, over and above task effects in these regressors.

**Fig. 4.**
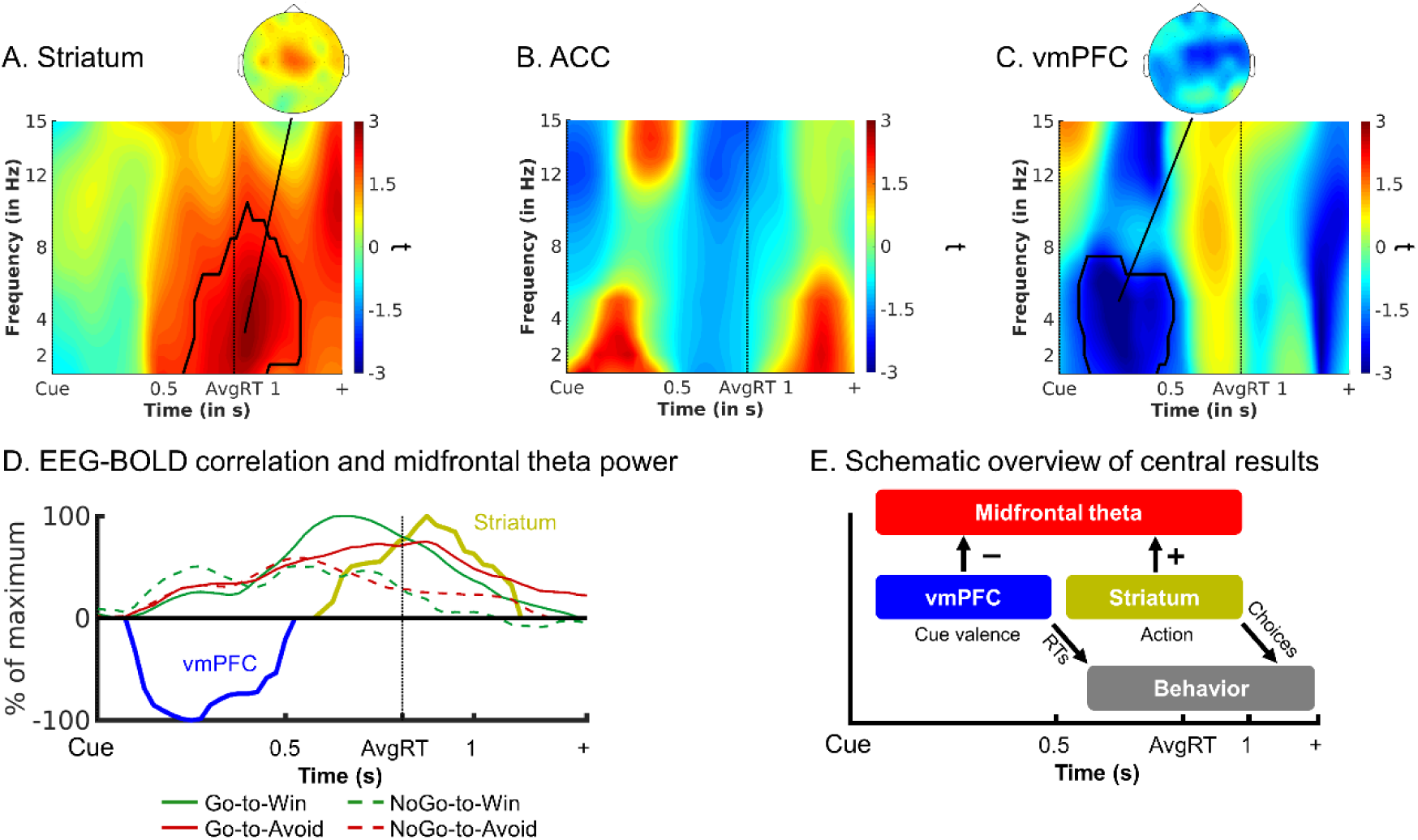
fMRI-informed EEG results. Uniquely explained variance in EEG time-frequency power over midfrontal electrodes (FCz/Cz) by BOLD signal from (A) whole striatum, (B) ACC, and (C) vmPFC to the average Group-level *t*-maps display the modulation of the EEG time-frequency power by trial-by-trial BOLD signal in the selected ROIs. Striatal (but not ACC) BOLD correlates most strongly with theta/delta power around the time of response. vmPFC BOLD correlates with broadband (peak: theta) power soon after cue onset. Areas surrounded by a black edge indicate clusters of |*t*| > 2 with *p* < .05 (cluster-corrected). Topoplots indicate the topography of the respective cluster. Note that there are no significant clusters in ACC, but given our a-priori hypothesis regarding the relation between ACC BOLD and midfrontal theta, for completeness, we include this visualization. (D) Time course of vmPFC and striatal BOLD correlations with theta power (*t*-values from clusters above threshold extracted and summed over frequencies), normalized to the peak of the time course of each region, overlaid with theta power for each valence x action condition. vmPFC-theta correlations emerge when theta is still similar for each condition, while striatum-theta correlations emerge when theta rises more strongly for Go than NoGo responses. (E) Schematic overview of our main EEG-fMRI results: Both vmPFC and striatum modulate midfrontal theta power (striatum likely indirectly via motor areas, see Supplementary Material S17). BOLD signal in both regions predicts the amplitude of theta power—the vmPFC early and negatively, the striatum late and positively. We speculate that the vmPFC encodes cue valence and sends this information to the striatum, where valence information biases the motivation for active responses in recurrent fronto-striatal loops and thus gives rise for motivational biases in behavioral responses and reaction times.

Next, to investigate whether vmPFC BOLD, which encoded cue valence rather than action, contributes to action selection in fronto-striatal circuits, we also assessed an association between midfrontal EEG power and BOLD in the vmPFC. Trial-by-trial deconvolved BOLD signal from the valence cluster in vmPFC correlated negatively with broadband power (*p* = .043, cluster-corrected) in an early cluster (around 2–15 Hz, peak in the upper theta range around 6–7 Hz, 0–0.4 sec., Fig. 4C). Regression of the time-domain EEG voltage on vmPFC BOLD yielded a negative association of vmPFC BOLD with a left frontal P2 component, which however showed a different topography and was unlikely to explain the negative vmPFC-theta correlations over midfrontal electrodes (see Supplementary Material S16).

Finally, we performed complementary EEG-informed fMRI analyses using the trial-by-trial midfrontal alpha and theta signals identified in EEG-only analyses as regressors on top of the task regressors. While fMRI-informed EEG analyses allow to test which region is the best predictor of a certain EEG signal (competing with BOLD signal from other regions), this EEG-informed fMRI analysis allows to assess whether networks of several regions might reflect trial-by-trial changes in power (although none of them might do so uniquely). Alpha power correlated negatively with dlPFC and SMG BOLD, while theta power correlated positively with BOLD signal in bilateral pre-SMA, ACC, motor cortices, operculum, putamen, and cerebellum, corroborating the association between theta and motor regions, including the striatum (see Supplementary Material S17). See Supplementary Material S17 for a methodological discussion on these seemingly contrasting findings.

In sum, the fMRI-informed EEG analyses show that the amplitude of the ramping theta signal at the time of response correlates positively with striatal BOLD signal. In contrast, theta power early after cue onset correlates negatively with vmPFC activity. Finally, ACC activity in the cluster encoding Go vs. NoGo responses was not significantly linked to theta power, while other parts of ACC and pre-SMA (and further motor regions) were in fact related to trial-by-trial theta power. Taken together, we observed an early negative correlation of theta with vmPFC BOLD, which encoded cue valence, and a late positive correlation of theta (ramping up to the response) with striatal BOLD, which encoded the selected action.

### Summary of the main results

In sum, we observed motivational biases in both choice and reaction time data. Cue valence, which drives these biases, led to differences in BOLD signal in vmPFC (higher for Win cues) and ACC (higher for Avoid cues). These vmPFC and ACC BOLD responses to cue valence also predicted reaction times on a trial-by-trial basis. Striatal BOLD did not respond to cue valence, but predominantly reflect the action participants selected (higher for Go than NoGo). Motivational conflict was associated with higher BOLD in midfrontal cortex, though only weakly, and with early transient midfrontal alpha power. In contrast, midfrontal theta power reflected the selected action (higher for Go than NoGo) around the time of responses. Finally, trial-by-trial vmPFC BOLD correlated negatively with theta power early after cue onset, while striatal BOLD correlated positively with theta power around the time of responses.

## Discussion

The main aim of this study was to investigate the interaction of striatal and midfrontal processes during the suppression of motivational biases, using combined EEG-fMRI. We and others have previously found elevated theta power when such biases were successfully suppressed (Cavanagh et al. 2013; Swart et al. 2018). Furthermore, computational models of basal ganglia loops posited the origin of this bias in the striatum (Frank 2005; Collins and Frank 2014). We thus hypothesized that theta power would correlate with attenuated striatal valence coding during bias-incongruent actions. However, we found that both striatal and theta signals were strongly dominated by action per se, rather than by a combination of valence and action, as was predicted for the striatum, or by valence-action congruency, as was predicted for the midfrontal cortex. Most importantly, we found that the time course of theta power exhibited several features of a process accumulating evidence whether to select an active Go response or not. Interestingly, valence-driven vmPFC BOLD signal uniquely predicted variability in mid-frontal theta immediately following cue presentation, while action-driven striatal BOLD responses uniquely predicted variability in midfrontal theta around the time of the response. Taken together, these results suggest that striatum and vmPFC may act in concert to evaluate the value of performing an active Go response, i.e., the “value of work”.

### Striatal BOLD reflects motivation for action

We found that striatal BOLD signal was strongly dominated by the action participants performed (Go vs. NoGo) rather than cue valence. This finding replicates previous studies using a very similar task (Guitart-Masip et al. 2011; Guitart-Masip, Chowdhury, et al. 2012; Guitart-Masip, Huys, et al. 2012) and highlights the role of the striatum in value-based behavioral activation and invigoration (Taylor and Robbins 1986, 1984; Robbins and Everitt 1992, 2007; Niv et al. 2007; Salamone and Correa 2012; Howe and Dombeck 2016; Syed et al. 2016; Coddington and Dudman 2018, 2019; da Silva et al. 2018), yet appears to be at odds with the role of the striatum in reward expectation (Doya 2000; Dayan and Daw 2008; Collins and Frank 2014). A recent theory aimed to reconcile these roles by proposing that striatal (dopamine) signals do not reflect the value of anticipated outcomes per se, but rather the value of performing an action to obtain this outcome, i.e. the “value of work” (Berke 2018). Recent empirical work in rodents has indeed shown selective ramping of striatal dopamine signals for active responses approaching a goal state (Hamid et al. 2016, 2019; Syed et al. 2016; Mohebi et al. 2019). In light of these findings, it seems plausible that the striatum evaluates whether there are sufficient incentives to overcome a NoGo default and to instead take action to achieve a valuable goal. Any signal that reflects such an evolving value of Go should ramp over the trial time course and peak when a response is elicited, like an evidence accumulation process. While the BOLD response is too sluggish for capturing such fast within-trial signals, EEG can provide insight.

### Late midfrontal theta power reflects striatal activity

In this study, midfrontal theta power, like striatal BOLD, was modulated by whether participants made an active Go response or not. Theta power bifurcated for Go and NoGo responses, peaking around the time of the response. Striatal BOLD and midfrontal theta signals were strongly linked, such that trial-by-trial fluctuations in striatal (rather than ACC or motor cortex) BOLD was the best predictor of trial-by-trial fluctuations in theta power around the time of response.

The observed link between striatal BOLD and midfrontal theta may seem surprising given that previous EEG source localization studies of conflict-related midfrontal theta power modeled a source in ACC or pre-SMA (Hanslmayr et al. 2008; Cohen and Ridderinkhof 2013), and previous resting-state EEG-fMRI studies reported negative correlations of frontal theta with regions of the default-mode network (Scheeringa et al. 2008, 2009). Our study might fill a blind spot in these literatures: by recording EEG-fMRI during a decision task, we show that theta power increases commonly observed in such tasks might reflect subcortical action selection processes. Arguably, the striatum is far away from the scalp and thus unlikely to be the direct neural source of midfrontal theta oscillations. It is possible that striatal action selection processes modulate activity in parts of midfrontal (or motor) cortex, reflected in the amplitude of theta power over the scalp (see also our EEG-informed fMRI analyses in Supplementary Material S17). This finding suggests that scalp EEG can give insights into evolving action selection in the striatum, which is not visible in resting-state recordings, but can only be studied using appropriate task designs.

In contrast to our study, previous findings have reported elevated theta power mostly in situations of cognitive conflict (Cavanagh and Frank 2014; Cohen 2014), including our own EEG study using the same task (Swart et al. 2018). Importantly, however, these studies usually observe strong theta rises for *any* action, with conflict-induced theta constituting a minor increase on top of this much larger rise (Cohen and Cavanagh 2011; Swart et al. 2018). While conflict-induced theta was absent in our data, action-induced theta was strongly present— which might be especially visible in Go/NoGo tasks such as in this study, but concealed in other paradigms that only feature active responses. We speculate that both phenomena are related and reflect the evolving value of making a Go action: theta power rises prior to Go actions, but even further in situations of cognitive conflict. If response thresholds in striatal pathways are elevated during conflict, theta may not reflect a cortical top-down “trigger” that drives threshold elevation, but rather the extra bits of accumulated evidence in the striatum that follow from such elevated response thresholds. This alternative account of midfrontal theta power modulation provides a putative unifying explanation for both action- and conflict-induced theta increases.

### Early vmPFC valence signals shape action selection

So far, we have suggested that striatal BOLD and theta power signals reflect how evidence for action is accumulated, putatively reflecting the value of work (i.e., the physical effort of taking action). This interpretation leaves open what drives this evidence accumulation—and does so differently for reward and punishment prospects, leading to the observed expression of motivational biases in behavior. Any neural “source” of these biases should show differential activity in response to Win and Avoid cues. While valence coding was weak and spatially heterogeneous in the striatum, it clearly emerged in vmPFC (positively) and ACC (negatively). Particularly the vmPFC appears to be a likely candidate source of motivational biases given that a wealth of previous studies (Haber and Knutson 2010; Bartra et al. 2013) has shown vmPFC BOLD to encode the expected outcomes. In behavior, cue valence affected both the probability and the speed of making a Go response, as people responded more often and faster to Win cues than to Avoid cues. A follow-up trial-by-trial analysis showed that the vmPFC is likely involved in eliciting this motivational bias, as fluctuations in vmPFC signal predicted response times also within each valence condition. This finding is consistent with the idea that valence information in this region feeds into fronto-striatal loops and gives rise to motivational biases in behavior. Of note, the region of ACC encoding cue valence did not significantly correlate with midfrontal theta power, even though trial-by-trial ACC BOLD did correlate with RTs. Taken together, in line with past theories of recurrent fronto-striatal loops (Alexander et al. 1986; Mink 1996; Middleton and Strick 2000; Gurney et al. 2001), our result suggest that vmPFC encodes cue valence at an early time point and then biases the motivation for active responses, i.e. the value of work (Hamid et al. 2016; Berke 2018), in the striatum.

vmPFC BOLD correlated negatively with midfrontal theta power very early after cue onset (Fig. 4C), consistent with previous EEG-fMRI findings (Scheeringa et al. 2008, 2009; Hauser et al. 2015) as well as electrophysiological data in humans (Harris et al. 2011; Hunt et al. 2012) and animals (Van Wingerden et al. 2010; Vinck et al. 2010; Seo et al. 2012; Knudsen and Wallis 2020). The vmPFC shows a more positive BOLD response to Win (compared to Avoid) cues. The observed negative vmPFC-theta correlations are in line with previous findings showing that both vmPFC and midfrontal theta power encode valence, though with opposite signs: vmPFC BOLD is typically higher for positive than negative events, while the opposite holds for midfrontal theta (and midfrontal BOLD signal) (Shackman et al. 2011; Cavanagh, Figueroa, et al. 2012; Cavanagh, Zambrano-Vazquez, et al. 2012; Braem et al. 2017). Our results indicate that vmPFC encodes cue valence very soon after this information becomes available, indexed in midfrontal theta power.

In contrast to the negative vmPFC-theta correlation immediately following cue onset, action-related theta and positive striatum-theta correlations occurred later, around the time of response (Cohen 2014). Although both vmPFC and striatal BOLD correlated with power in the same frequency band, correlations may well reflect different neural processes. Compared to vmPFC-theta correlations, striatum-theta correlations and in particular action-related theta exhibited a more centroparietal (rather than midfrontal) topography of rather short duration (for a discussion of the burst-like modulations of ongoing theta oscillations, see Cohen 2014). Of note, timing and topography of action-related theta in the EEG-only analysis were similar to the “centroparietal positivity”/ P300, which has been suggested to reflect perceptual evidence accumulation (O’Connell et al. 2012; Kelly and O’Connell 2013; Philiastides et al. 2014; Twomey et al. 2015). However, this action-related theta signal was not visible in cue-locked ERP analyses and thus apparently not phase-locked (see Supplementary Material S08). Taken together, early vmPFC-theta correlations likely reflect cue valence processing, while late striatum-theta correlations likely reflect the motivation for a final action.

### Caveats and open questions

A priori, we expected elevated midfrontal theta power in situations of motivational incongruency between biases and required actions. Instead of theta power, we observed a transient increase in midfrontal alpha power, which specifically occurred on incongruent trials on which participants successfully overcame biases. Trial-by-trial midfrontal alpha power was negatively correlated with BOLD signal in dorsolateral prefrontal cortex and supramarginal gyrus. While we are not aware of previous literature reporting elevated frontal alpha in conflict situations, the observed associations with BOLD in regions of the fronto-parietal attention network might point at midfrontal alpha reflecting an unspecific mechanism of focused attention and increased task engagement (Brier et al. 2010; Harris et al. 2013; Helfrich et al. 2018) which might help to retrieve and focus on learned stimulus-response associations (Buschman et al. 2012). Of note, while heightened attention is typically associated with decreased (posterior) alpha, there have been findings of increased alpha, as well (Van Der Meij et al. 2016). Nonetheless, future research is needed to understand the role midfrontal alpha might play in overcoming motivational conflict.

The absence of elevated theta power in situations of cognitive conflict could perhaps be due to relatively low performance in the current study compared to previous studies using the same task (Swart et al. 2017, 2018). This reduced performance is likely due to the fMRI environment and associated necessary task changes (longer and jittered response-outcome intervals), and may explain the inconsistency with previous findings (Swart et al. 2018). If participants learned the required action of a cue less well, they would be less aware of conflict between motivational bias and action requirements. Furthermore, reduced performance resulted in relatively fewer trials on which participants successfully overcame motivational biases, leaving less statistical power to test neural hypotheses on bias suppression. This might explain why we did not observe conflict-related midfrontal theta increases and why evidence for increased BOLD signal in pre-SMA during conflict was rather weak. Thus, our findings do not undermine the role of theta in motivational conflict, but rather highlight a putatively complementary role of theta in the motivation of actions.

Finally, the response-locked nature of the observed theta power difference raises the question whether this finding might simply reflect motor execution. Control analyses indicated that the signal was modulated by response hand and accuracy, which can be expected from a signal that reflects decision variables such as response conflict or threshold variability, but not from a signal that reflects simple motor execution or even a signal artifact. Moreover, the observation that striatal BOLD was the best predictor of trial-by-trial midfrontal theta power (rather than BOLD in motor cortices) again speaks for theta reflecting a pre-response, decision-related processes.

## Conclusion

In sum, participants in this simultaneous EEG-fMRI study exhibited strong motivational biases and relatively poor instrumental learning in a motivational Go-NoGo learning task. This feature likely has rendered our set-up suboptimal for isolating the predicted fronto-striatal mechanisms involved in suppressing motivational biases. However, the presence of these strong (relatively uncontrolled) motivational biases enabled us to further dissect the mechanisms of bias expression. Specifically, the finding that, despite strong valence effects on behavior, striatal BOLD indexed the selected action (rather than cue valence) indicates that the striatum is unlikely to play a role in generating the motivational bias. Rather, striatal BOLD might selectively reflect the motivation to show an active Go response. The finding of strong cue valence signaling in the vmPFC, which also predicted reaction times on a trial-by-trial basis, suggests that the motivational bias might instead arise from the vmPFC. The negative association of this vmPFC signal with early midfrontal theta suggests that the vmPFC processes valence information very early after cue onset and may subsequently shape action selection. One putative mechanism through which the vmPFC could shape action selection is the modulation of the rate of evidence accumulation towards a Go response. Besides the negative correlation of vmPFC BOLD with midfrontal theta power early after cue onset, the positive correlation of striatal BOLD with late midfrontal theta power concurs with a complementary role for the striatum in the eventual decision to execute an active response. Together, these findings suggest a dual nature of midfrontal theta power, with early components reflecting valence processing in the vmPFC and late components reflecting motivation for action in the striatum. Taken together, our results are in line with “value of work” theories of the role of fronto-striatal loops in the evaluation of whether to perform an active response.

## Funding

This work was supported by the Netherlands Organization for Scientific Research (NWO) under a research talent (grant number 406-14-028 to J.C.S.), a VENI grant (grant number 451-12-021 to R.S.), a VICI grant (grant number 453-14-005 to R.C.), an Ammodo KNAW Award 2017 (to R.C.), a VIDI grant (grant number 452-17-016 to H.E.M.d.O.), and by the James S. McDonnell Foundation under a James McDonnell scholar award to (R.C.).

## Acknowledgments

We thank Emma van Dijk for assistance with data collection, Michael J. Frank for helpful discussions, Tobias U. Hauser and Laurence Hunt for sharing code for fMRI-informed EEG analyses, and the weekly Donders M/EEG meeting for discussions of these results and many helpful suggestions.

## Competing Interests

All authors declare that no competing interests exist.

## Data availability

All data, code and materials necessary to reproduce the experiments, analyses, and results are available on the Donders Repository under the following URL: https://data.donders.ru.nl/login/reviewer-113095877/El7ZhUKAkr_qz0yK-sl86OtoILxMChDaEmwsBlRxjsc The URL provided should only be shared with reviewers. The data will only be publicly available after manuscript acceptance, in which case researchers can access the data after signing a data use agreement. In line with requirements of the Ethics Committee and the Radboud University security officer, potentially identifying data (such as imaging data) can only be shared to identifiable researchers, hence the requirement for registration and for requesting access. Neither authors nor data steward is involved in granting access to external researchers, this is only based on the complete registration of the researcher and follows a “click-through” procedure.

## Supplementary Materials

### S01: Behavioral, fMRI, and EEG analyses with only the 30 participants included in EEG-fMRI analyses

We repeated the behavioral, fMRI, and EEG analyses reported in the main text while excluding the six participants that were also not included in the fMRI-inspired EEG analyses in the main text: two participants due to fMRI co-registration failure, which were also not included in the fMRI-only analyses, and five further participants due to large outliers on the *b*-maps in the fMRI-inspired EEG analyses.

In this subgroup, similar to the entire sample, participants performed significantly more Go responses to Go cues than NoGo cues (Required action: χ^2^(1) = 25.77, *p* < .001; see Fig. S1A-B). Furthermore, participants showed a motivational bias, as they performed more Go actions for Win cues than Avoid cues (Valence: χ^2^(1) = 18.87, *p* < .001). The interaction of Valence x Required Action was not significant (χ^2^(1) = 0.13, *p* = .910). Similarly, when making a (Go) response, participants responded faster when this was correct (Go cues) than when this was incorrect (NoGo cues) (Required action: χ^2^(1) = 21.02, *p* < .001). Furthermore, a motivational bias was present also in RTs, with significantly faster responses to Win than Avoid cues (Valence: χ^2^(1) = 37.31, *p* < .001). Again, the interaction was not significant (χ^2^(1) = 2.25, *p* = .134; see Fig. S1C). In sum, all behavioral results also held in this subsample.

**Figure S1A.**
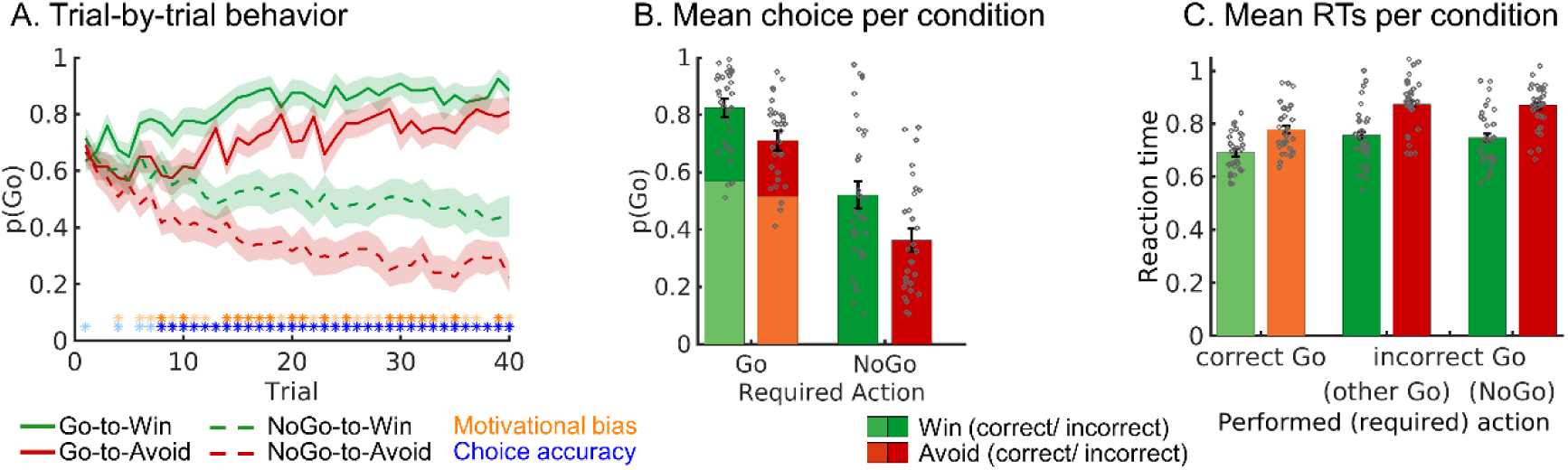
Motivational Go/NoGo learning task performance in the subgroup of 29 participants included in the fMRI-inspired EEG analyses. (A) Trial-by-trial proportion of Go responses (±SEM) for Go cues (solid lines) and NoGo cues (dashed lines). Shadows indicate standard errors for per-condition-per-participant means across participants using the Cousineau-Morey method (Morey 2008). The motivational bias is defined as the tendency to make more Go actions to Win than Avoid cues (i.e., green lines are above red lines). Additionally, participants clearly learn whether to make a Go actions or not (solid lines go up, dashes lines go down). Orange asterisks below indicate trial-by-trial significance of motivational bias, blue asterisks indicate performance accuracy above chance (i.e. correct GoLeft, GoRight or NoGo response) (light color: p < .05 uncorrected; dark color: p < 0.0013; Bonferroni corrected for number of trials). (B) Mean (±SEM) proportion Go responses per cue condition (points are individual participants’ means). Proportion Go responses is higher for Go than NoGo cues, indicative of task learning, and higher for Win than Avoid cues, reflecting the influence of motivational biases on behavior. (C) Mean (±SEM) reaction times for correct and incorrect Go responses, the latter split up in whether the other Go response or the NoGo response would have been correct (points are individual participants’ means). Participants respond faster on correct than on incorrect Go responses and faster to Win than Avoid cues, reflecting the influence of motivational biases on behavior.

In our fMRI analyses, when testing for differences in BOLD signal between Win and Avoid cues, there were again no significant clusters in the striatum in a whole-brain corrected analysis. When restricting our analyses to an anatomical mask of the striatum, BOLD signal in both left (*z*max = 4.35, *p* = .00485, MNI peak coordinates: xyz = [-8 4 4]) and right medial caudate nucleus (*z*max = 3.98, *p* = .0121, xyz = [12 6 6]) was higher for Avoid than Win cues as reported in the main text (Fig. S2C), while differences in left posterior putamen reported in the main text were not significant any more (Fig. S2B). Furthermore, at a whole-brain level cluster correction, BOLD signal was again higher for Win compared to Avoid cues in vmPFC (*z*max = 5.09, *p* = 2.24e-16, xyz = [-6 44 2]), dlPFC (*z*max = 5.01, *p* = .000142, xyz = [16 48 48]), PCC (*z*max = 4.34, *p* = .000297, xyz = [6 -44 32]), and left amygdala/ hippocampus (*z*max = 4.36, *p* = .0162, xyz = [-20 -2 -22]). There were additional clusters in left vlPFC (zmax = 4.30, *p* = .00702, xyz = [28 36 -10]), left middle temporal gyrus (zmax = 3.76, *p* = .014, xyz = [-62 -18 -12]),) right middle temporal gyrus (zmax = 3.92, *p* = .014, xyz = [62 -18 -6]), left angular gyrus (*z*max = 4.99, *p* = 5.07e-6, xyz = [-44 -56 20]), and right amygdala/ hippocampus (*z*max = 4.52, *p* = .0223, xyz = [20 -6 -20]) not featured in the results in the main text. Conversely, in line with results reported in the main text, BOLD was higher for Avoid stimuli in dorsal ACC (*z*max = 4.12, *p* = 1.15e-05, xyz = [2 36 46]), left insula (*z*max = 4.47, *p* = .0014, xyz = [-28 22 0]), left superior frontal gyrus (*z*max = 4.12, *p* = .00398, xyz = [-22 -4 56]), right insula (*z*max = 4.07, *p* = .00148, xyz = [32 26 0]), left vlPFC (*z*max = 4.47, *p* = .00697, xyz = [-32 62 8]), right superior frontal gyrus (*z*max = 3.99, *p* = .00588, xyz = [22 -4 52]), and right precuneous (*z*max = 4.60, *p* = .00156, xyz = [8 -64 54]), in line with results reported in the main text (see Fig. S2D). In addition, we observed clusters in left frontal pole (*z*max = 4.67, *p* = .00702, xyz = [-32 62 8]) and left angular gyrus (*z*max = 3.98, *p* = .0162, xyz = [-38 -56 48]).

When correlating RTs and BOLD signal, there was again a significantly negative correlation between RTs and vmPFC BOLD, *t*(29) = -3.89, p < 0.001, *d* = -0.71, and a significantly positive correlation between RTs and ACC BOLD, *t*(29) = 7.41, *p* < 0.001, *d* = 1.35 (see Fig. S2E).

When testing for differences in BOLD signal between responses, we observed again significantly higher BOLD signal for *Go* than *NoGo* action in the entire striatum (bilateral caudate nucleus, putamen, and nucleus accumbens), thalamus, and bilateral cerebellum (*z*max = 7.05, *p* = 0, xyz = [-12 -24 10]), ACC (*z*max = 6.87, *p* = 9.04e-05, xyz = [0 8 42]), left motor cortex (zmax = 4.59, *p* = .00915, xyz = [-54 -22 -24], Fig. S2F), in line with results reported in the main text. Furthermore, there was an additional (separate) cluster in left frontal pole (*z*max = 4.23, *p* = .0124, xyz = [-28 42 6]. Again, there were not clusters with higher BOLD signal for NoGo than Go responses.

When testing for differences in BOLD signal between bias-incongruent and bias-congruent actions, there were no clusters at a whole-brain cluster level significance correction, and—different from results report in the main text—also not when restricting the analysis to a mask comprising ACC and pre-SMA (Fig. SG-H).

**Figure S1B.**
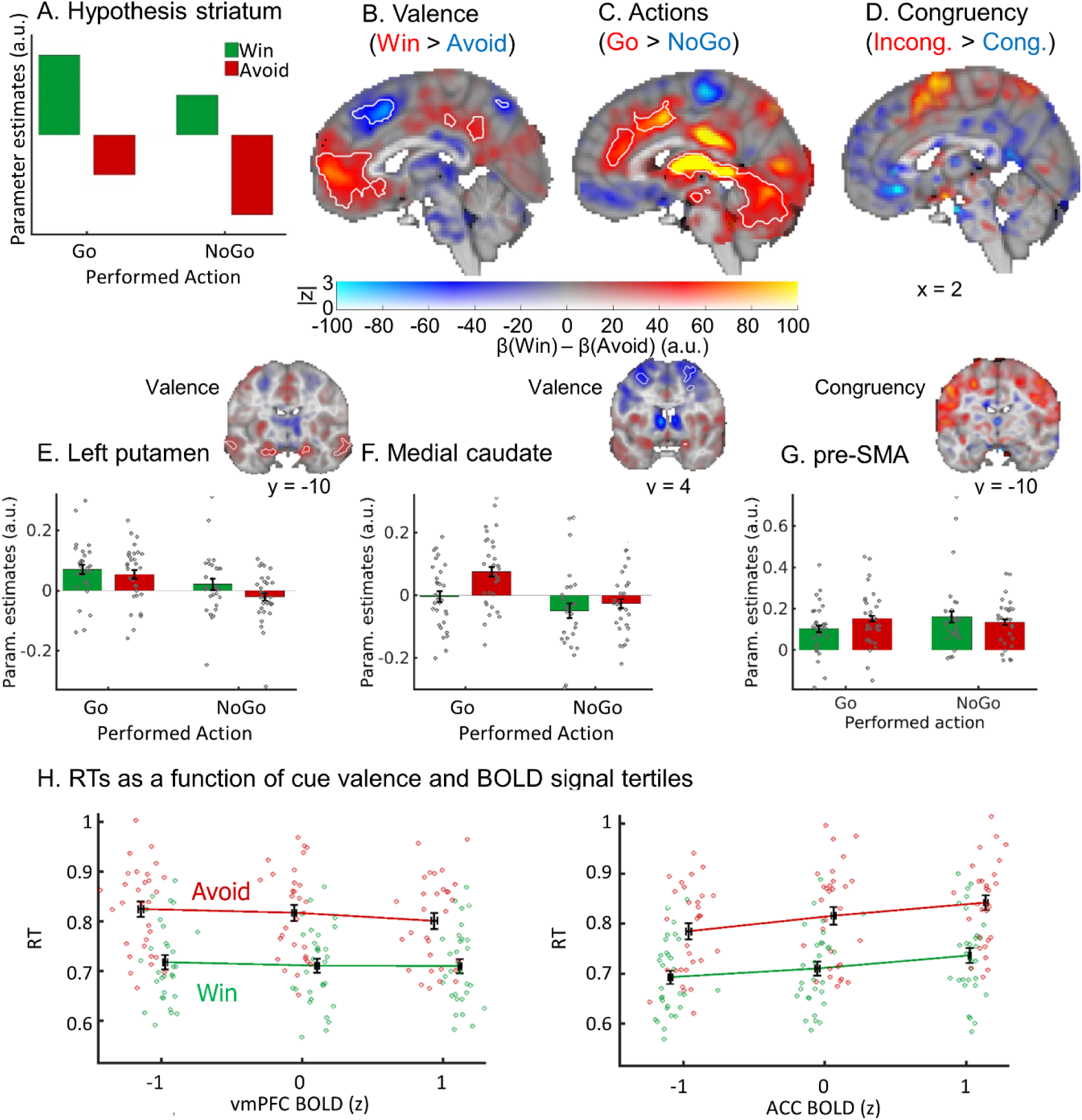
BOLD signal as a function of cue valence, performed action, and congruency in the subgroup of 29 participants included in the fMRI-inspired EEG analyses. (A) We hypothesized striatal BOLD to encode cue valence (strong main effect of valence), with an attenuation of this valence signal when actions incongruent with bias-triggered actions were performed (weak main effect of action). (B) BOLD signal was significantly higher for *Win* compared to *Avoid* cues in ventromedial prefrontal cortex (vmPFC; whole brain corrected) and left putamen (small-volume corrected), but higher for *Avoid* compared to *Win* cues in ACC and medial caudate (small-volume corrected). (C) BOLD signal was significantly higher for *Go* compared to *NoGo* actions in the entire striatum as well as ACC, thalamus, and cerebellum (all whole-brain corrected). (D) Based on the plot, it appears that BOLD signal was higher for bias-incongruent actions than bias-congruent actions in pre-SMA, but contrary to the results reported in the main text, this was not significant. B-BOLD effects displayed using a dual-coding data visualization approach with color indicating the parameter estimates and opacity the associated *z*-statistics. White contours indicate statistically significant clusters (p < .05), either small-volume corrected (orange) or whole-brain corrected (black). (E-G) Mean beta weights per task condition (x-axis) per participant (individual grey dots) in significant clusters in left putamen, medial caudate and pre-SMA (significant in small-volume correction). (E) It appears that left posterior putamen encoded valence positively (higher BOLD for Win than Avoid cues), but contrary to results reported in the main text, this was not significant. (F) Medial caudate encoded valence negatively (higher BOLD for Avoid than Win cues). (G) Extracted BOLD signal from pre-SMA to illustrate (non-significant) congruency effects. (H) Reaction times (RTs) as a function of cue valence and BOLD signal tertiles (z-standardized) per participant (individual dots; x-location relative to all other participants). RTs were significantly predicted by BOLD signal in vmPFC (positively) as well as by BOLD signal in ACC and striatum (negatively). BOLD-RT correlations were independent of cue valence.

In our EEG analyses, there was no significant difference between incongruent than congruent trials in the theta band (*p* = .236). In the alpha band, it was marginally significant (*p* = .052), most strongly around 200–325 ms after cue onset (Fig. S3E-F). A permutation test on the broadband (1–15 Hz) TF power yielded a significant result (*p* = 0.046).

Furthermore, broadband power (1–15 Hz) was significantly higher on trials with Go actions than NoGo actions (cue-locked: *p* = .002, around 550–1300 ms after cue onset, see Fig. S3A-B; response-locked: *p* = .002, around -150–425 ms relative to responses, see Fig. S3C-D). Overall, all EEG results also held in this subsample.

Taken together, behavioral and EEG analyses yielded identical conclusions as results reported in the main text. In the fMRI analyses, differences between Win and Avoid cues in left posterior putamen and differences in pre-SMA between bias-incongruent and bias-congruent actions were not significant, while all other results were still significant and yielded identical conclusions as results reported in the main text.

**Figure S1C.**
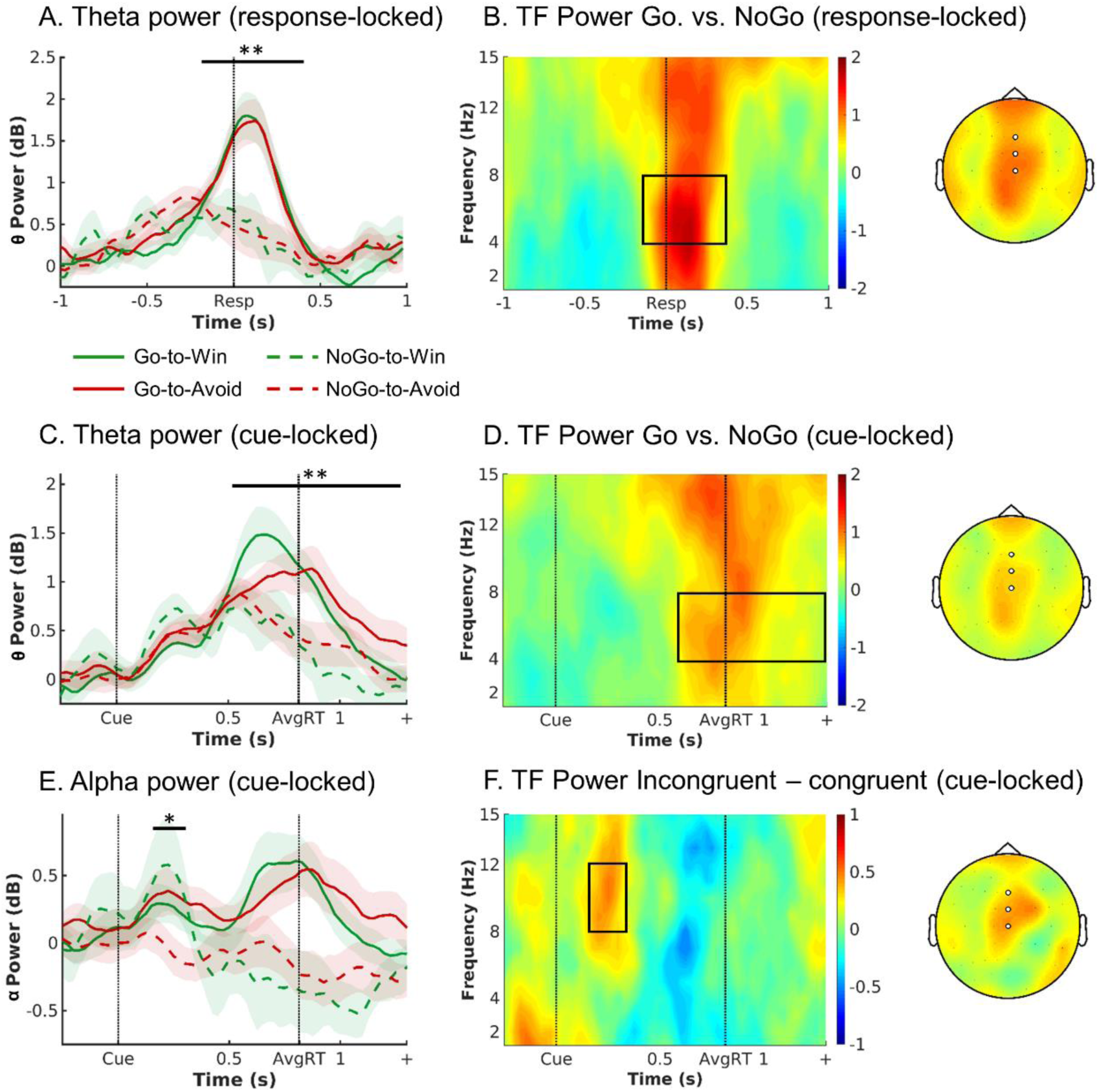
EEG time-frequency power as a function of cue valence and action. (A) Response-locked within trial time course of average theta power (4–8 Hz) over midfrontal electrodes (Fz/ FCz/ Cz) per cue condition (correct-trials only). Theta increased in all conditions relative to pre-cue levels, but to a higher level for Go than NoGo trials. There were no differences in theta peak height or latency between Go2Win and Go2Avoid trials. (B) Left: Response-locked time-frequency power over midfrontal electrodes for Go minus NoGo trials. Go trials featured higher broadband TF power than NoGo trials. The broadband power increase for Go compared to NoGo trials is strongest in the theta range. Right: Topoplot for Go minus NoGo trials. The difference is strongest at FZ and FCz electrodes. (C-D) Cue-locked within trial time course and time-frequency power. Theta increased in all conditions relative to pre-cue levels, but to a higher level for Go than NoGo trials, with earlier peaks for Go2Win than Go2Avoid trials. (E) Trial time course of average alpha power (8–13 Hz) over midfrontal electrodes per cue condition (correct trials only; cue-locked). Alpha power transiently increases for both incongruent conditions in an early time window (around 175–325 ms). (F) Left: Time-frequency plot displaying that the transient power increase was focused on the alpha band (8 – 13 Hz), leaking into the upper theta band. Right: Topoplot of alpha power displaying that this incongruency effect was restricted to midfrontal electrodes (highlighted by white disks). * p < 0.05. ** p < 0.01. Shaded errorbars indicate (±SEM). Box in TF plots indicates the time frequency window where t-values > 2.

### S02: Anatomical masks (for small-volume corrected analyses) and conjunctions of anatomical and functional masks (for fMRI-informed EEG analyses)

**Figure S03A.**
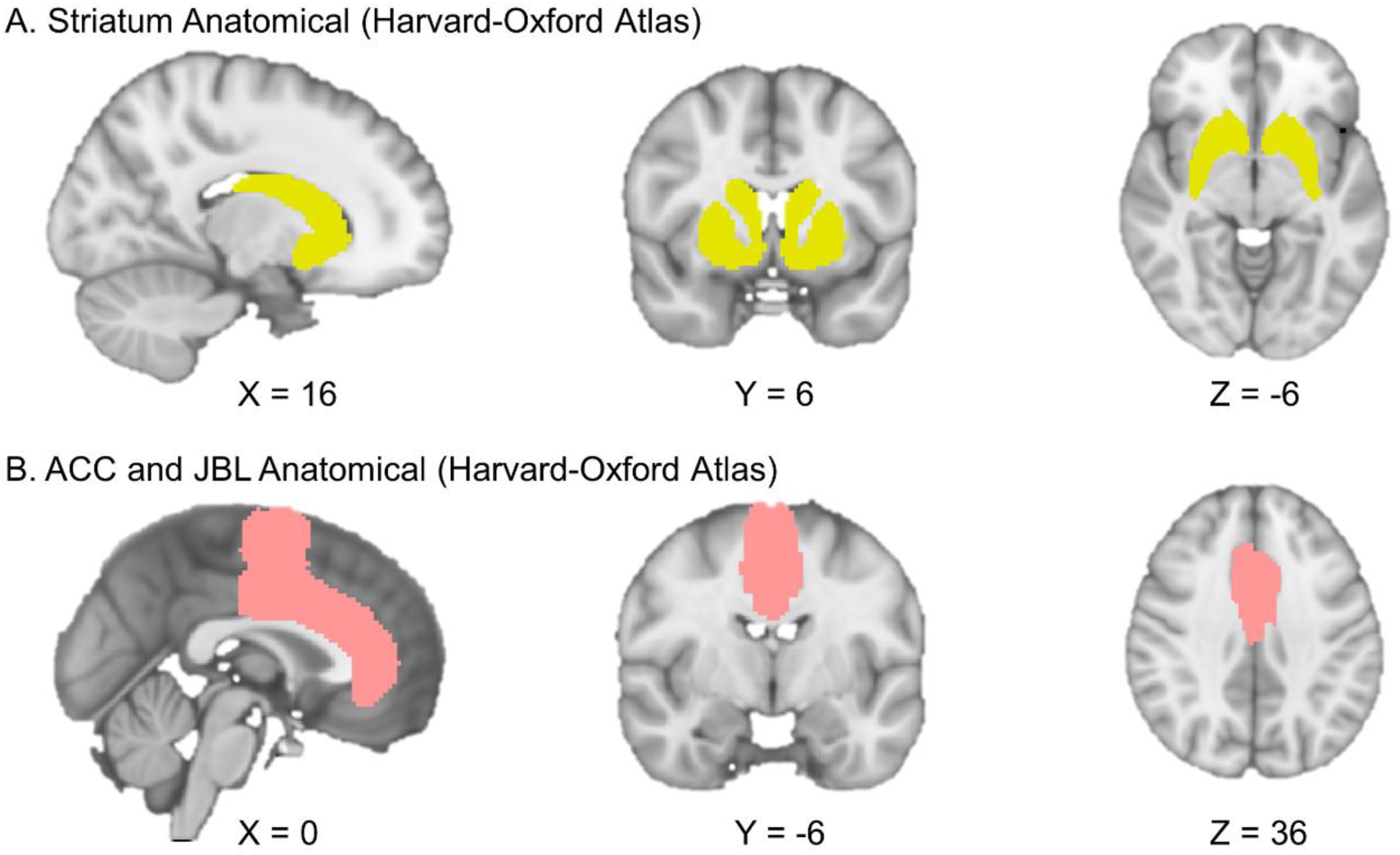
Anatomical masks of (A) striatum (yellow, conjunction of bilateral nucleus accumbens, caudate, and putamen) and (B) midfrontal cortex (pink, cingulate cortex anterior and juxtapositional lobule cortex) used for small-volume corrected GLM analyses. All masks were extracted from the probabilistic Harvard-Oxford Atlas, thresholded at 10%. Note that images are in radiological orientation (i.e., left brain hemisphere presented on the right and vice versa).

**Figure S03B.**
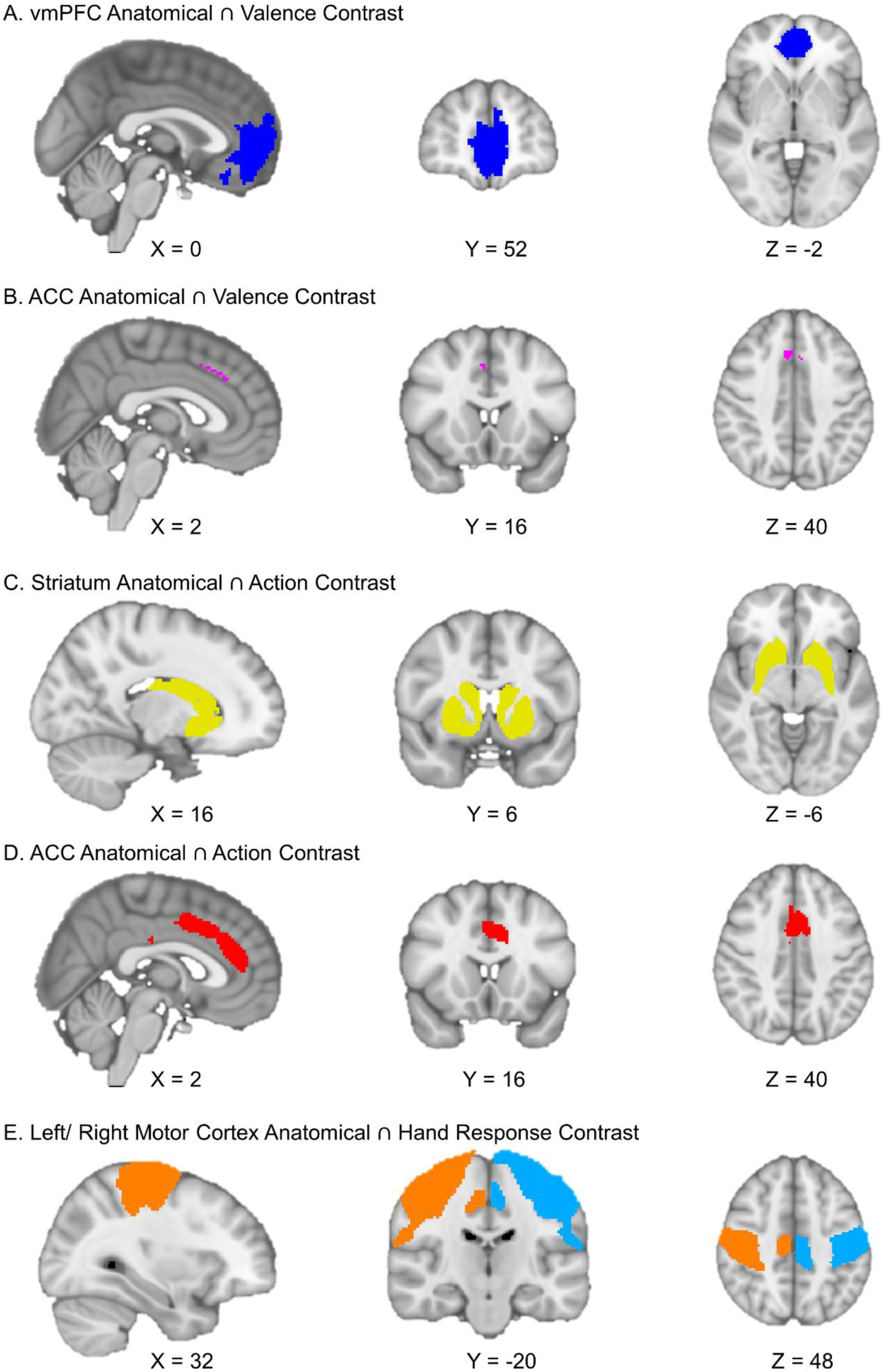
Conjunctions of anatomical masks (based on the Harvard-Oxford Atlas) and functional contrasts from fMRI GLM analyses (Valence, Action, and Hand Response contrasts) used for fMRI-informed EEG analyses: (A) vmPFC valence contrast (dark blue, conjunction of frontal pole, frontal medial cortex, and paracingulate gyrus), (B) ACC valence contrast striatum (purple, cingulate cortex anterior), (C) striatum action contrast (yellow, conjunction of bilateral nucleus accumbens, caudate, and putamen), (D) ACC action contrast (red, cingulate cortex anterior), and left (light blue) and right (orange) motor cortices hand response contrast (conjunction of precentral gyrus and postcentral gyrus) used for fMRI-informed EEG analyses. All anatomical masks were extracted from the probabilistic Harvard-Oxford Atlas, thresholded at 10%. Note that images are in radiological orientation (i.e., left brain hemisphere presented on the right and vice versa).

### S03: Regressors and contrasts in fMRI analyses

Regressors:

- Win2GoOnset: for every trial with Win cue and Go action, at cue onset, duration 1, value +1
- Win2NoGoOnset: for every trial with Win cue and NoGo action, at cue onset, duration 1, value +1
- Avoid2GoOnset: for every trial with Avoid cue and Go action, at cue onset, duration 1, value +1
- Avoid2NoGoOnset: for every trial with Avoid cue and NoGo action, at cue onset, duration 1, value +1
- Handedness: for every trial, at cue onset, value +1 for left hand response, 0 for NoGo response, -1 for right hand response
- Error: for every trial, at cue onset, value +1 for incorrect response, 0 for correct response
- utcomeOnset: for every trial, at outcome onset, value +1 for every trial
- utcomeValence: for every trial, value +1 for positive outcome (reward, no punishment), -1 for negative outcome (no reward, punishment)
- InvalidOutcome: for trials where uninstructed button was pressed, at outcome onset, value 1

Nuisance Regressors:

- Realignment Parameters (obtained from co-registration)
- white-matter signal
- out-of-brain signal
- regressor for each volume where relative displacement > 2mm

| Regressor | 1 | 2 | 3 | 4 | 5 | 6 | 7 | 8 | 9 |
| --- | --- | --- | --- | --- | --- | --- | --- | --- | --- |
| Contrast | Win2GoOnset | Win2NoGoOnset | Avoid2GoOnset | Avoid2NoGoOnset | Handedness | Error | OutcomeOnset | OutcomeValence | InvalidOutcome |
| 1) Valence | 1 | 1 | -1 | -1 |  |  |  |  |  |
| 2) Executed Action | 1 | -1 | 1 | -1 |  |  |  |  |  |
| 3) Conflict | -1 | 1 | 1 | -1 |  |  |  |  |  |
| 4) Hand |  |  |  |  | 1 |  |  |  |  |

### S04: Significant BOLD clusters in the valence, action, and congruency contrasts

**Table S04.**
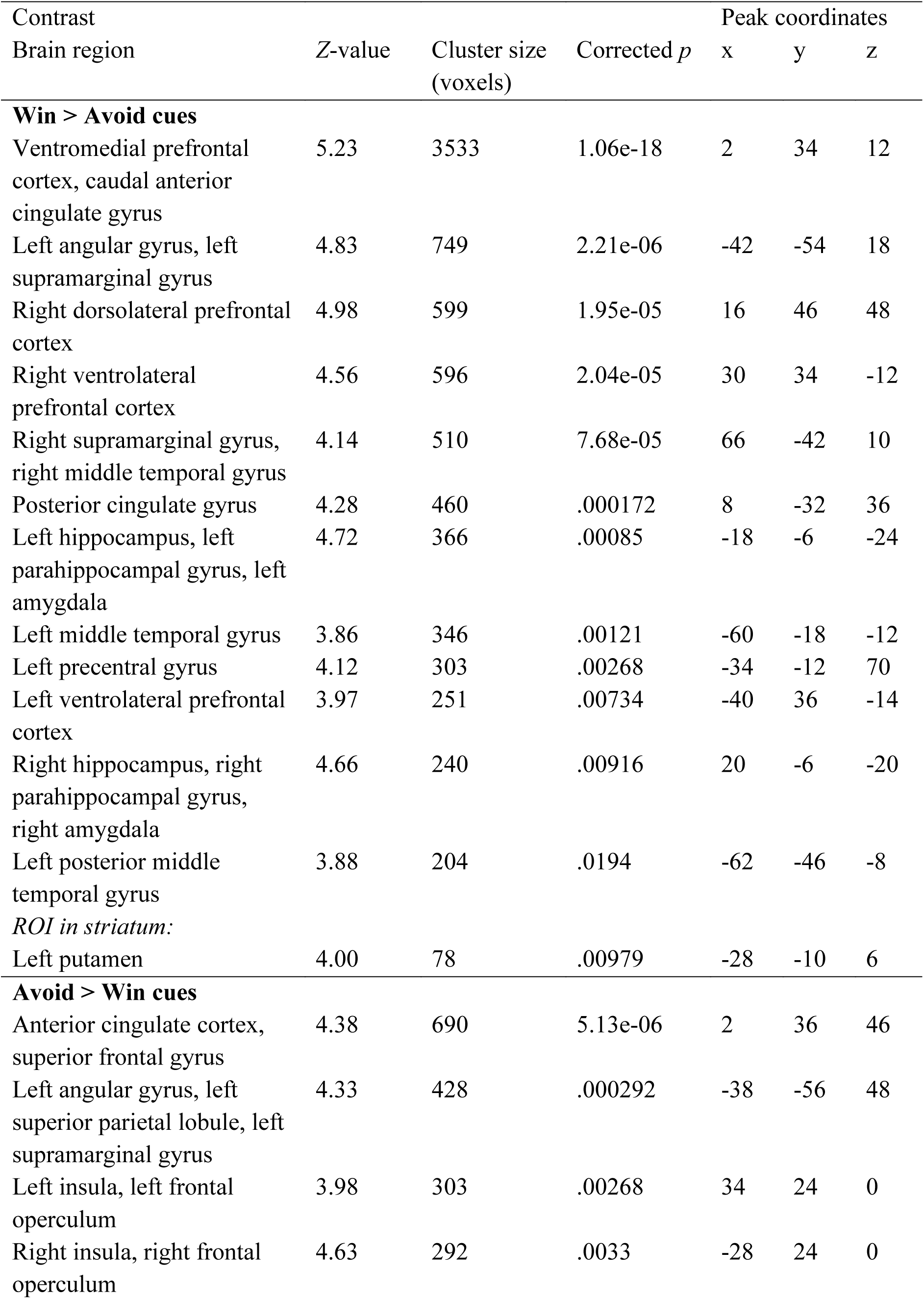
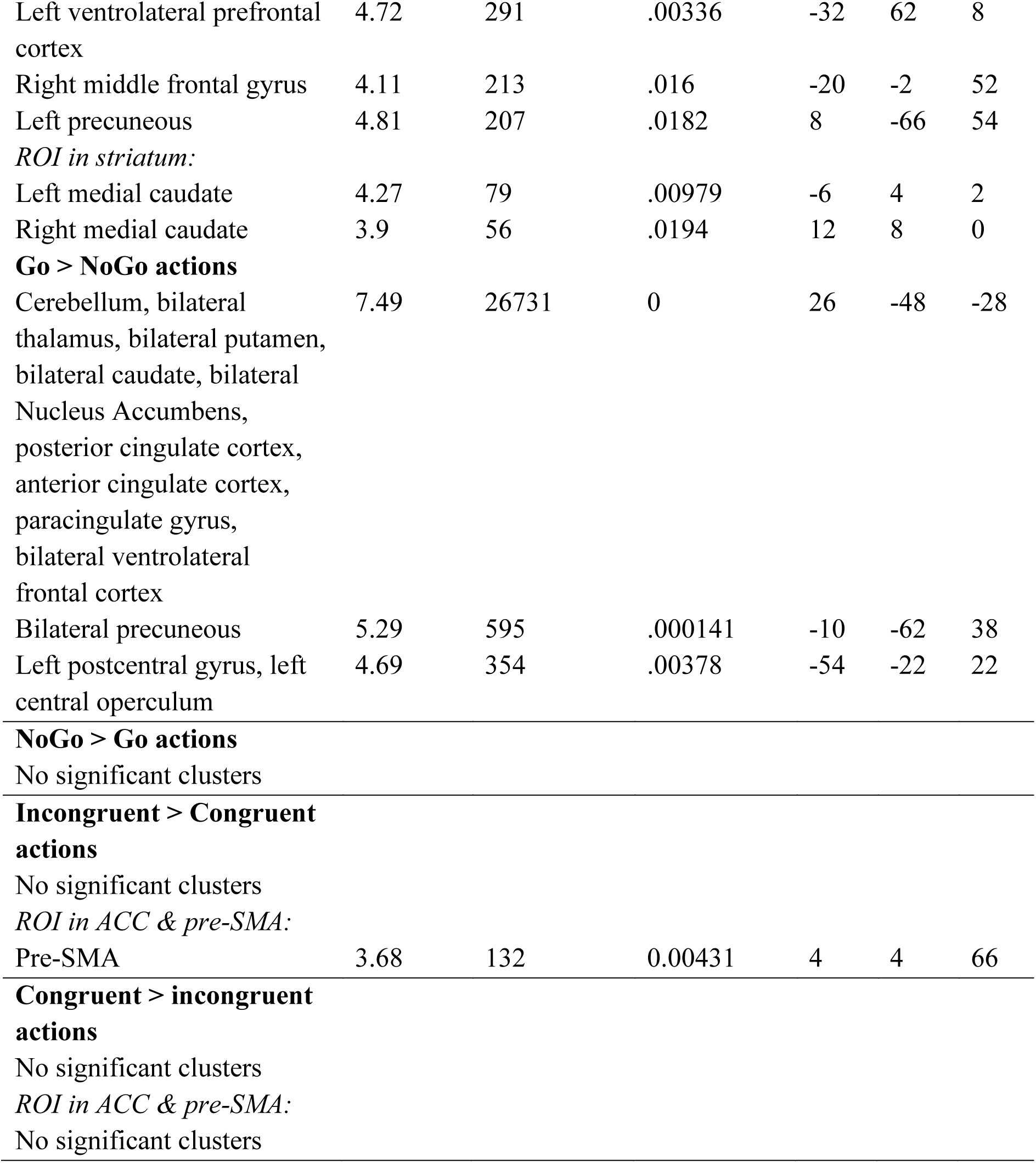
Significant clusters in the valence, action and congruency contrasts in the fMRI GLM.

### S05: Changes in activity over time

After identifying BOLD correlates of cue valence, performed action, and motivational conflict in the whole-brain and small-volume-corrected GLM analyses reported in the main text, we were interested in whether these effects change over the time course of the experiment. For this purpose, we extracted the first eigenvariate of the BOLD signal from the significant clusters above threshold (see Fig. 2; for masks, see S2), fitted an HRF to each trial to obtain the trial-by-trial HRF amplitude (identical procedure to BOLD-RT correlations and fMRI-informed EEG analyses), and analyzed these amplitude as a function of the respective behavioral variable (cue valence, performed action, or motivational conflict), trial number, and their interaction, using mixed-effects linear regression.

Specifically, for vmPFC, ACC, left putamen and medial caudate signal, we fitted the following model (Wilkinson notation):

*BOLD ∼ cueValence * trialNumber + (cueValence * trialNumber|participant)*

vmPFC signal was strongly modulated by cue valence, χ^2^(1) = 31.313, *p* < .001, with higher signal for Win than Avoid cues. In addition, the main effect of trial number was marginally significant, χ^2^(1) = 3.351, *p* = .067, with signal tending to increase over time. The interaction between valence and trial number was marginally significant as well, χ^2^(1) = 2.959, *p* = .085: The valence effect tended to decrease over time, driven by signal increasing for Avoid cues while staying at a constant high level for Win cues.

ACC signal was also strongly modulated by valence, χ^2^(1) = 15.213, *p* < .001, with higher BOLD signal for Avoid than Win cues. There also was a significant main effect of trial number, χ^2^(1) = 6.491, *p* = .011, with signal decreasing over time. The interaction between valence and trial number was marginally significant, χ^2^(1) = 2.935, *p* = .087: The valence effect tended to decrease over time, driven by signal decreasing for Avoid cues while staying at a constant low level for Win cues.

Signal in left putamen strongly encoded cue valence, χ^2^(1) = 16.949, *p* < .001, with higher signal for Win than Avoid cues. The main effect of trial number was not significant, χ^2^(1) = 1.265, *p* = .261, and neither was the interaction between valence and trial number, χ^2^(1) = 1.544, *p* = .214.

Signal in medial caudate strongly encoded cue valence, χ^2^(1) = 17.330, *p* < .001, with higher signal for Avoid than Win cues. The effect of trial number was just significant, χ^2^(1) = 3.874, *p* = .049, with signal decreasing over time. The interaction between valence and trial number was marginally significant, χ^2^(1) = 3.769, *p* = .052: The valence effect tended to decrease over time, driven by signal decreasing for Avoid cues while staying at a constant low level for Win cues.

For striatal and ACC signal (different mask than for the valence signal reported above), we fitted the following model (Wilkinson notation):

*BOLD ∼ performedAction * trialNumber + (performedAction * trialNumber| participant)*

For striatal signal, the main effect of action was not significant, *t*(31.14) = 0.031, *p* =

.975, while the effect of trial number was strongly significant, *t*(217.09) = -2.773, *p* = .006, with signal decreasing over time. The interaction was marginally significant, *t*(54.61) = 1.736, *p* = .088, driven by signal increasing for Go actions, but decreasing for NoGo actions, such that the action effect in striatum (higher signal for Go than NoGo actions) only emerged over time (because models using likelihood ratio tests failed to converge, *p*-values in this model are instead based on *t*-tests using Satterthwaite’s method as implemented in the R package *lmerTest*).

For ACC signal, the main effect of action was not significant, χ^2^(1) = 0.270, *p* = .603, while the main effect of trial number was significant, χ^2^(1) = 5.342, *p* = .021, reflecting overall decreasing signal over time. The interaction between action and trial number was not significant, χ^2^(1) = 0.038, *p* = .845. This inconsistency with our results in the whole-brain GLM analyses might reflect differential weighting of outliers and block-wise signal in FSL’s FEAT vs. lme4’s mixed effects models.

For pre-SMA signal, we fitted the following model (Wilkinson notation):

*BOLD ∼ conflict * trialNumber + (conflict * trialNumber|participant)*

For pre-SMA signal, there was a significant main effect of conflict, χ^2^(1) = 5.064, *p* = .024, with higher BOLD for bias-incongruent than -congruent action, and a significant negative effect of trial number, χ^2^(1) = 10.530, *p* = .001, with signal decreasing over time. The interaction between conflict and trial number was not significant, χ^2^(1) = 0.142, *p* = .706.

**Figure S05.**
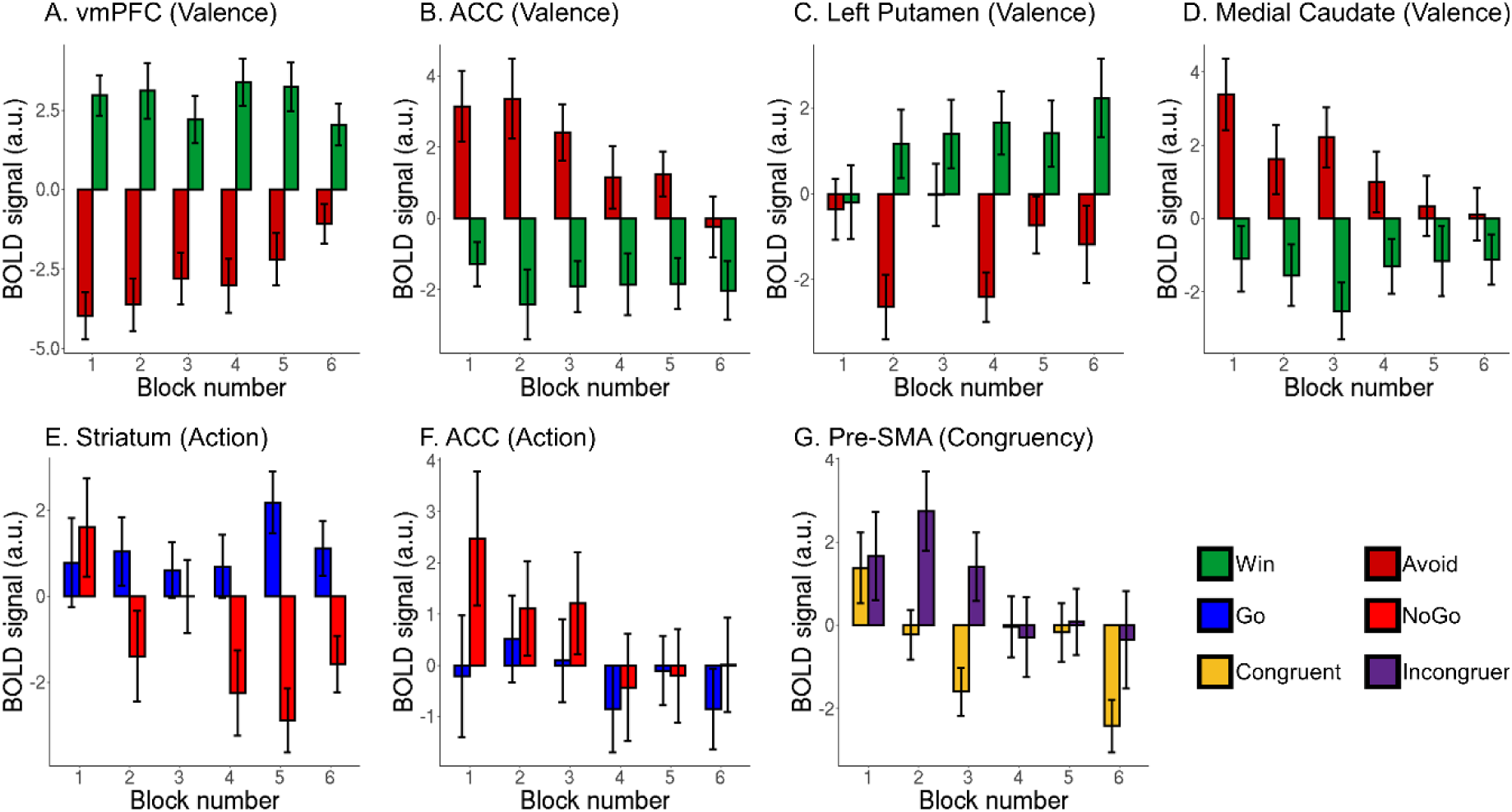
BOLD signal in significant clusters identified with whole-brain and small-volume-corrected GLM analyses (see main text, Figure 2) as a function of behavioral variables (cue valence, executed action, motivational conflict) and block number. Bar represent means, whiskers present standard errors (per condition over participants, computed via the Cousineau-Morey method). (A) vmPFC encoded cue valence (Win > Avoid), but this effect tended to decrease over time. (B) ACC encoded cue valence (Avoid > Win), but this effect tended to decrease over time. (C) Left putamen encoded cue valence (Win > Avoid). (D) Medial caudate encoded cue valence (Avoid > Win), but this effect tended to decrease over time. (E) Striatum encoded the performed action (Go > NoGo), but this effect only emerged over time. (F) In contrast to whole-brain GLM analyses, ACC did not significantly encode the performed action. (G) Pre-SMA encoded motivational conflict (congruency, incongruent > congruent).

### S06: fMRI results for correct trials only

EEG and fMRI research have different analytical procedures of dealing with differences between correct and incorrect trials: While fMRI research typically uses multiple linear regression (GLMs), which allows to model error trials by a designated regressor, EEG research typically tests for differences between (categorical) conditions with a (mass-univariate) *t*-test approach. Because we used both approaches in the main text, here, for consistency, we also report fMRI results for regressors defined for correct trials only. Note that this analysis uses less trials than the one featured in the main text and thus has lower statistical power.

We fitted a GLM with eight task regressors, namely the four conditions resulting from crossing cue valence (Win/Avoid) and performed action (Go/NoGo irrespective of Left vs. Right Go) separately for correct and incorrect trials. We again added four regressors of no interest, namely response side (Go left = +1, Go right = -1, NoGo = 0), outcome onset (intercept of 1 for every outcome), outcome valence (reward = +1, punishment = -1, neutral = 0), and invalid trials (invalid buttons pressed and thus not feedback given). Note that compared to the GLM reported in the main text, we did not add an error regressor. This GLM failed to converge for one participant, leaving 33 participants in the group-level analysis.

When comparing BOLD signal between trials with Win cues and with Avoid cues, in the whole-brain corrected analysis, we again observed higher BOLD for Win cues in vmPFC (*z*max = 5.20, *p* = 1.3e-9, xyz = [0 40 2]), as well as left superior lateral occipital cortex (*z*max = 3.59, *p* = .00325, xyz = [-56 -64 30]), and left medial temporal gyrus (*z*max = 3.63, *p* = .00474, xyz = [-70 -14 -14]; Fig. S4D). Conversely, BOLD signal was higher for Avoid cues in left supramarginal gyrus (*z*max = 3.91, *p* = .00235, xyz = [-36 -48 34]) and left ventrolateral prefrontal cortex (*z*max = 2.34, *p* = .00453, xyz = [-26 58 4]). Note that higher activity in ACC is clearly visible in Fig. S4D, but not statistically significant. Furthermore, analyses using small-volume correction on an anatomical mask of the striatum yielded no clusters of differential BOLD activity, also not in the regions reported in the main text, i.e. in left putamen (Fig. S4B) nor medial caudate (Fig. S4C). Overall, whole-brain results on correct trials only were similar to the results across both correct and incorrect trials reported in the main text, but weaker, suggesting that restricting analyses to correct trials only resulted in a considerable loss in statistical power.

When comparing trials with Go vs. NoGo actions, we observed higher BOLD signal for Go than NoGo actions in clusters in bilateral cerebellum, thalamus, striatum, and ACC (*z*max = 7.01, *p* = 0, xyz = [-30 -50 -30]), right ventrolateral prefrontal cortex (*z*max = 4.38, *p* = 1.79e-07, xyz = [34 48 6]), precuneous (*z*max = 5.21, *p* = 1.97e-05, xyz = [-8 -64 38]), left operculum (*z*max = 4.72, *p* = .000216, xyz = [-52 -22 18]), right supramarginal gyrus (*z*max = 4.99, *p* = .000351, xyz = [-40 -50 36]), left precentral gyrus (*z*max = 4.15, *p* = .00186, xyz = [-44 -18 62]), and right precentral gyrus (*z*max = 3.98, *p* = .00758, xyz = [46 -18 66]; Fig. S4E). This finding is in line with results across both correct and incorrect trials reported in the main text. Conversely, BOLD signal was higher for NoGo than Go trials in left inferior frontal gyrus (*z*max = 4.43, *p* = .0128, xyz = [-58 26 20]), a finding not observed across both correct and incorrect trials reported in the main text.

Finally, when comparing both incongruent and congruent trials, there were again no significant clusters in a whole-brain corrected analysis (Fig. S4F), and also not in an analysis using small-volume correction on midfrontal cortex (Fig. S4G). Again, this null result might be due to a considerate loss in power compared to the results across both correct and incorrect trials reported in the main text.

**Figure S06.**
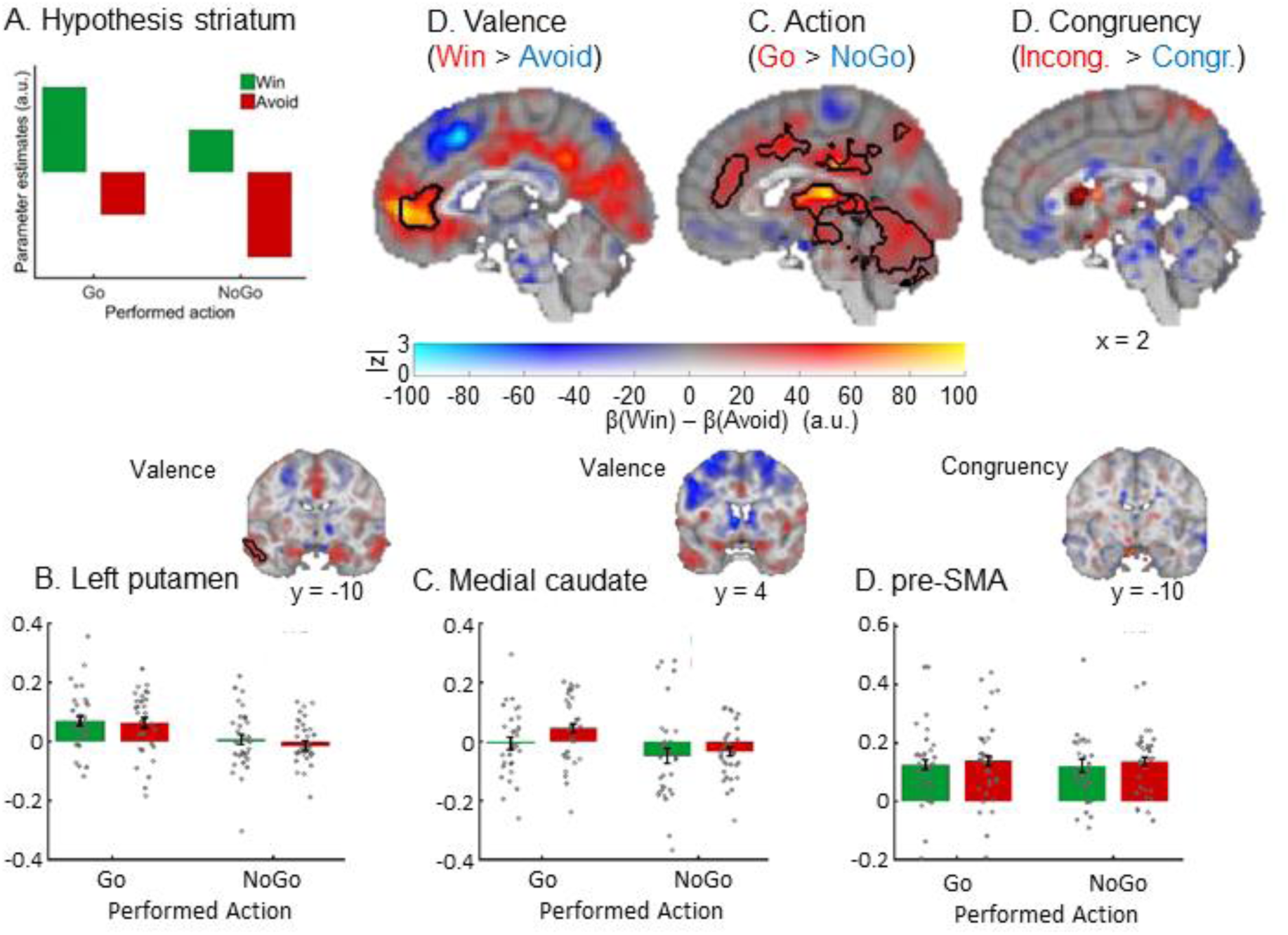
BOLD signal as a function of cue valence, performed action, and congruency for correct trials only. (A) We hypothesized striatal BOLD to encode cue valence (main effect of valence), with an attenuation of this valence signal when actions incongruent to the bias-triggered actions were performed (main effect of action). (B) BOLD signal was significantly higher for *Win* compared to *Avoid* cues in ventromedial prefrontal cortex (vmPFC; whole brain corrected), but in contrast to analyses across both correct and incorrect trials reported in the main text, BOLD was not significantly higher for *Avoid* compared to *Win* cues in ACC. (C) BOLD signal was significantly higher for *Go* compared with *NoGo* actions in the entire striatum as well as ACC, thalamus, and cerebellum (all whole-brain corrected). (D) BOLD signal was not significantly different between bias-incongruent actions (Go actions to Avoid cues and NoGo actions to Win cues) and bias-congruent actions (Go actions to Win cues and NoGo actions to Avoid cues), also not in the cluster in pre-SMA reported in the main text (small-volume corrected). B-D. BOLD effects displayed using a dual-coding data visualization approach with color indicating the parameter estimates and opacity the associated *z*-statistics. Contours indicate statistically significant clusters (p < .05), either small-volume corrected (striatal and SMA contours explicitly linked to a bar plot) or whole-brain corrected (all other contours). (E) Numerically, left posterior putamen seemed to encode valence positively (higher BOLD for Win than Avoid cues), but in contrast to analyses across both correct and incorrect trials reported in the main text, this was not significant. (F) Numerically, medial caudate seemed to encode valence negatively (higher BOLD for Avoid than Win cues), but in contrast to analyses across both correct and incorrect trials reported in the main text, this was not significant. (G) Extracted BOLD signal from pre-SMA to illustrate (the lack of) congruency effects.

### S07: ERPs as function of action and valence

Given that the observed phasic alpha increase occurred soon after stimulus onset and much earlier than the theta effect in our previous study (Swart et al., 2018)—although more similar to the timing reported by Cavanagh and colleagues (2013)—we investigated whether conditions differed in evoked rather and induced activity.

First, we again selected correct trials only, computed average ERPs for each condition per participant, and then tested for significant differences between the ERPs for incongruent and congruent trials using permutation tests on the average signal over midfrontal channels (Fz/ FCz/ Cz) in the time period of 0–700 ms post-cue (where evoked potentials occurred in the condition-averaged plot). We found no significant clusters in which the ERPs differed (no clusters above threshold; see Fig. S5A and D). Visual inspection yielded an inconsistent picture such that, if anything, N1, N2 and P3 components tended to be slightly stronger on incongruent trials, while P2 components tended to be stronger on congruent trials. Numerically, when comparing all four conditions, the P2 seemed to be highest and the N2 lowest on Go2Avoid trials, which the opposite was the case for NoGo2Win trials (see Fig. S6). Such opposite findings cannot explain why both conditions showed an increase in alpha power (see Fig. 3E in the main text), suggesting that the observed alpha power findings are not reducible to evoked activity.

Next, in line with the analyses in time-frequency space, we analyzed ERPs as a function of executed action and cue valence, contrasting trials with Go vs. NoGo actions and trials with Win vs. Avoid actions using permutation tests over the average signal of midfrontal electrodes (Fz/ FCz/ Cz). We found that ERPs differed significantly for Go vs. NoGo responses (*p* = .008) around 200–350 ms after cue onset, reflecting higher P2 (and lower N2) components for Go compared to NoGo responses (Fig. S5B). The peak of the topography of this effect was over left and central frontal electrodes (Fig. S5E). When contrasting ERPs for Win vs. Avoid cues, we only obtained a marginally significant *p*-value of .068, which was driven by higher signal 420–470 ms after cue onset (Fig. S5C and F). This difference occurred over midfrontal electrodes at the moment that the evoked signal rose towards the P3, but did not reflect differences in any of the component peaks.

**Figure S07A.**
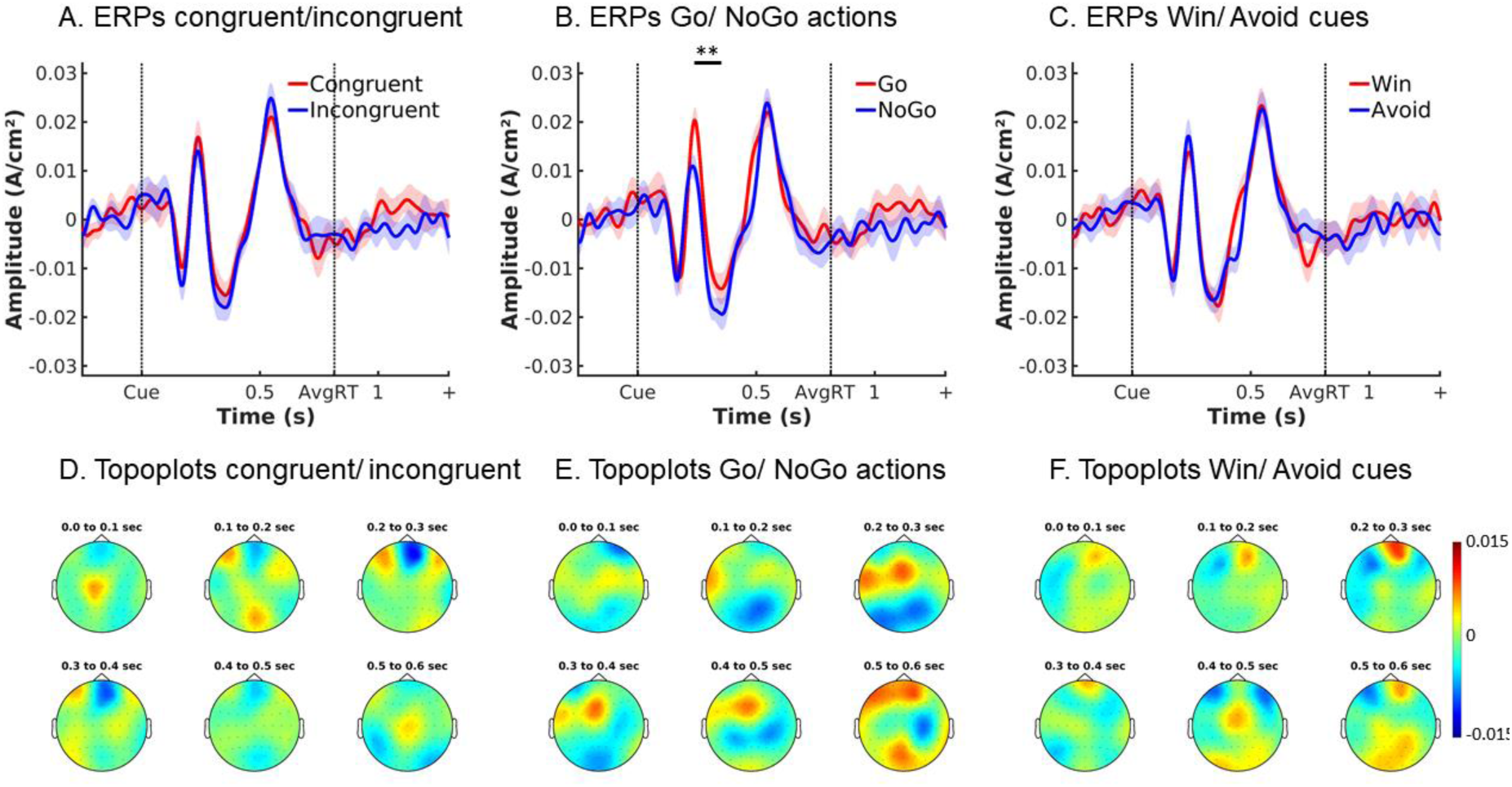
ERPs (±SEM) as a function of congruency, action, and cue valence over midfrontal electrodes (Fz/FCz/Cz; correct trials only). (A) There was no difference in midfrontal between congruent and incongruent trials, showing that the transient alpha effect observed on incongruent trials (main text Fig. 3E-F) was not reducible to evoked activity. (B) The frontal P2 component was stronger for Go compared to NoGo actions (and N2 respectively weaker). ** *p* < 0.01. (C) There was no difference between ERPs on Win and Avoid cues—apart from a small difference when the signal rises towards the P3 peak. (D-F) Topoplots displaying differences in ERPs between (D) congruency, (E) action, and (F) valence conditions in steps of 100 ms from 0 to 600 ms.

**Figure S07B.**
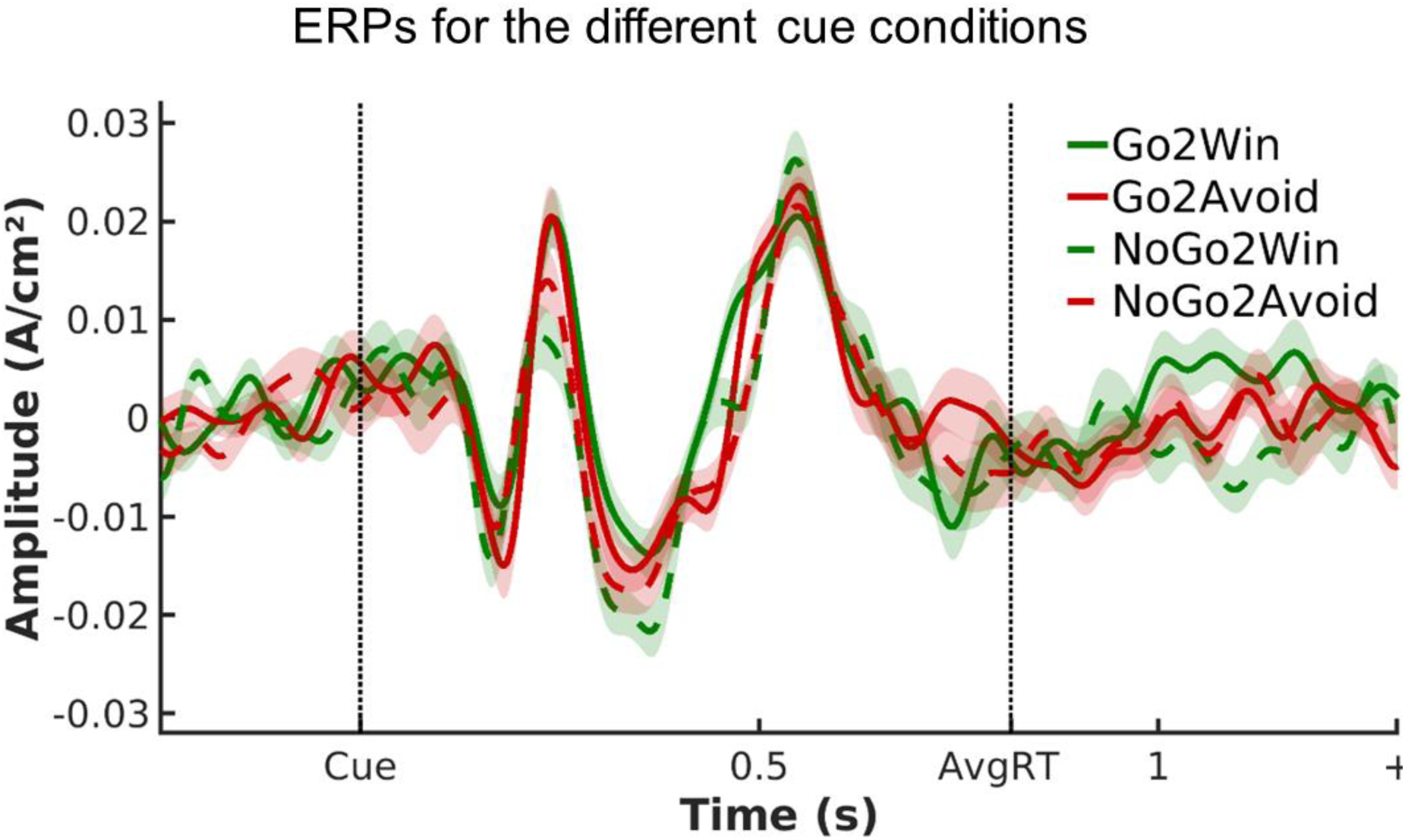
ERPs (±SEM) as a function of cue valence and action (correct trials only) over midfrontal electrodes (Fz/FCz/Cz).

### S08: Conflict-related alpha power after ERPs are subtracted

To test whether the observed earlier phasic alpha increase for incongruent compared to congruent conditions was attributable to evoked rather than induced activity, we removed evoked components from our data (correct trials only) by computing the average ERP for each condition per participant and subtracting it from the trial-by-trial data before performing time-frequency decomposition (Cohen & Donner, 2013). A permutation test on the alpha band yielded the same early phasic alpha increase for incongruent compared to congruent actions (*p* = .024; see Fig. S7) as reported in the main text (see Fig. 3E-F), suggesting that early alpha increase reflected induced rather than evoked activity.

**Figure S08.**
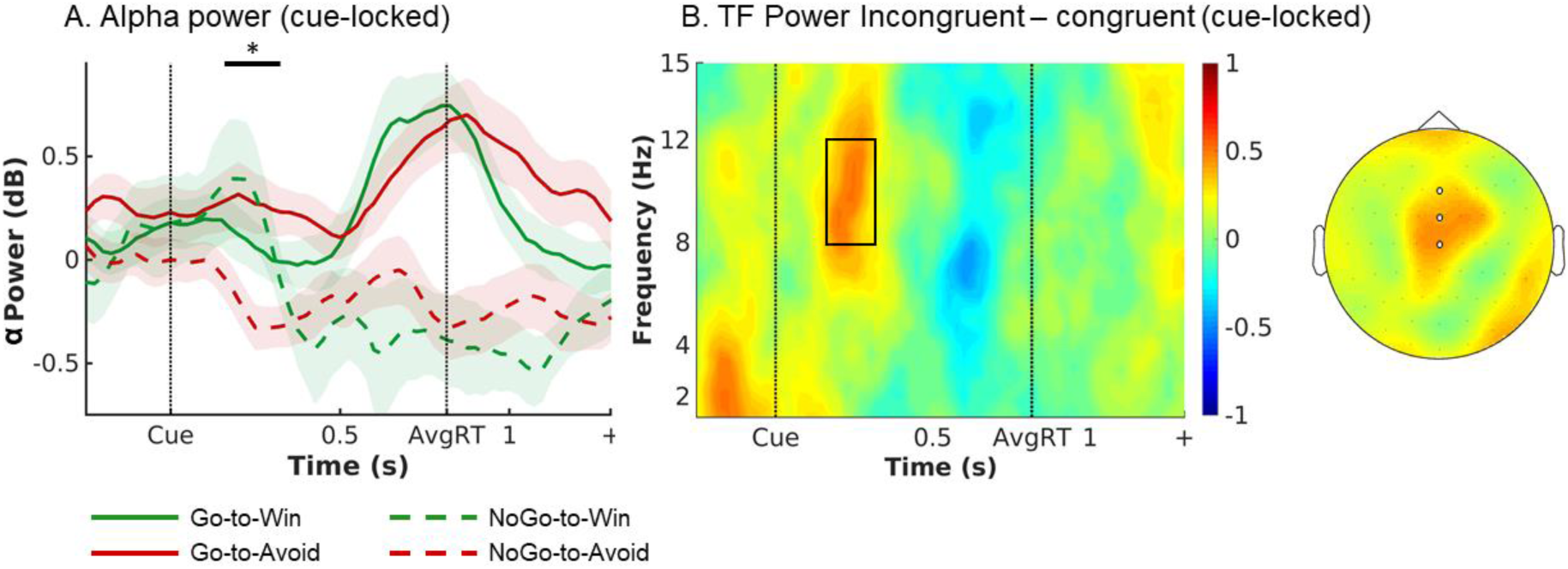
EEG alpha power with stimulus-locked ERPs subtracted. (A) Trial time course of average (±SEM) alpha power (8–13 Hz) over midfrontal electrodes (Fz/FCz/Cz) per cue condition (correct-trials only; stimulus-locked). Alpha power transiently increases for both incongruent conditions in an early time window of around 175–325 ms. The time window where the tested data shows *t*-values > 2 is indicated by the box. * *p* < 0.05. (B) Left: Time-frequency plot displaying that the transient power increase was focused on the alpha band, leaking into upper theta. Right: Topoplot of alpha power displaying that this incongruency effect was restricted to midfrontal electrodes (highlighted by white disks).

### S09: Alpha signal as a function of cue valence, required action, and correctness

Given that we did not expected motivational conflict to be encoded in an early phasic signal in the alpha band, we conducted follow-up analyses. If this alpha signal reflected conflict detection that was causally involved in suppressing motivational biases, it should occur only when incongruent trials where met with the correct response, but be attenuated or even absent when those trials where met with an incorrect response, i.e. when participants failed to detect and/or overcome biases (Swart et al., 2018). Furthermore, the signal should occur only on incongruent trials, but not congruent trials, reflecting conflict detection mechanisms that are selectively recruited on incongruent trials rather than (possibly attentional) mechanisms improving accuracy more globally.

For this purpose, instead of global permutation tests across time and frequencies, we extracted average oscillatory power in a focal window of 175–325 ms after cue onset in the range of 8–13 Hz, averaged over midfrontal electrodes (Fz/ FCz/ Cz), for each participant, and performed repeated-measures ANOVAs with the independent variables valence (Win/ Avoid), required action (Go/ NoGo), and accuracy (correct/ incorrect; see also Swart et al., 2018).

The RM-ANOVA yielded a significant main effect of valence, *F*(1, 35) = 6.930, *p* = .013, η^2^ = 0.007, a significant two-way interaction between valence and action, *F*(1, 35) = 8.368, *p* = .006, η^2^ = 0.008, but also a significant three-way interaction between valence, action, and accuracy, *F*(1, 35) = 5.103, *p* = .03, η^2^ = 0.005 (see Fig. S8). The main effect of accuracy was not significant, *F*(1, 35) = 2.02, *p* = .164, η^2^ = 0.003, suggesting that the observed alpha effect did not reflect an (attentional) process that was overall conducive to higher accuracy. For correct trials, we found the expected two-way interaction between valence and action, *F*(1, 35) = 9.582, *p* = .004, η^2^ = 0.023, in absence of significant main effects, reflecting that alpha power was indeed higher on correct incongruent trials than correct congruent trials, *t*(35) = 3.096, *p* = .004, *d* = 0.397. This reproduces the result of the permutation test from the main text. In contrast, for incorrect trials, we found only a significant main effect of valence, *F*(1, 35) = 8.637, *p* = .006, η^2^ = 0.011, reflecting overall higher alpha for Win than Avoid cues, *t*(35) = 2.939, *p* = .006, *d* = 0.490, but no significant interaction between valence and action, *F*(1, 35) = 0.239, *p* = .628, η^2^ < 0.001. Incorrect incongruent trials did not lead to significantly higher alpha power than incorrect congruent trials, *t*(35) = 0.489, *p* = .628, *d* = 0.081.

These additional findings are in line with increased alpha power reflecting a conflict detection mechanism that is selectively recruited on incongruent trials on which biases are successfully overcome.

**Figure S09.**
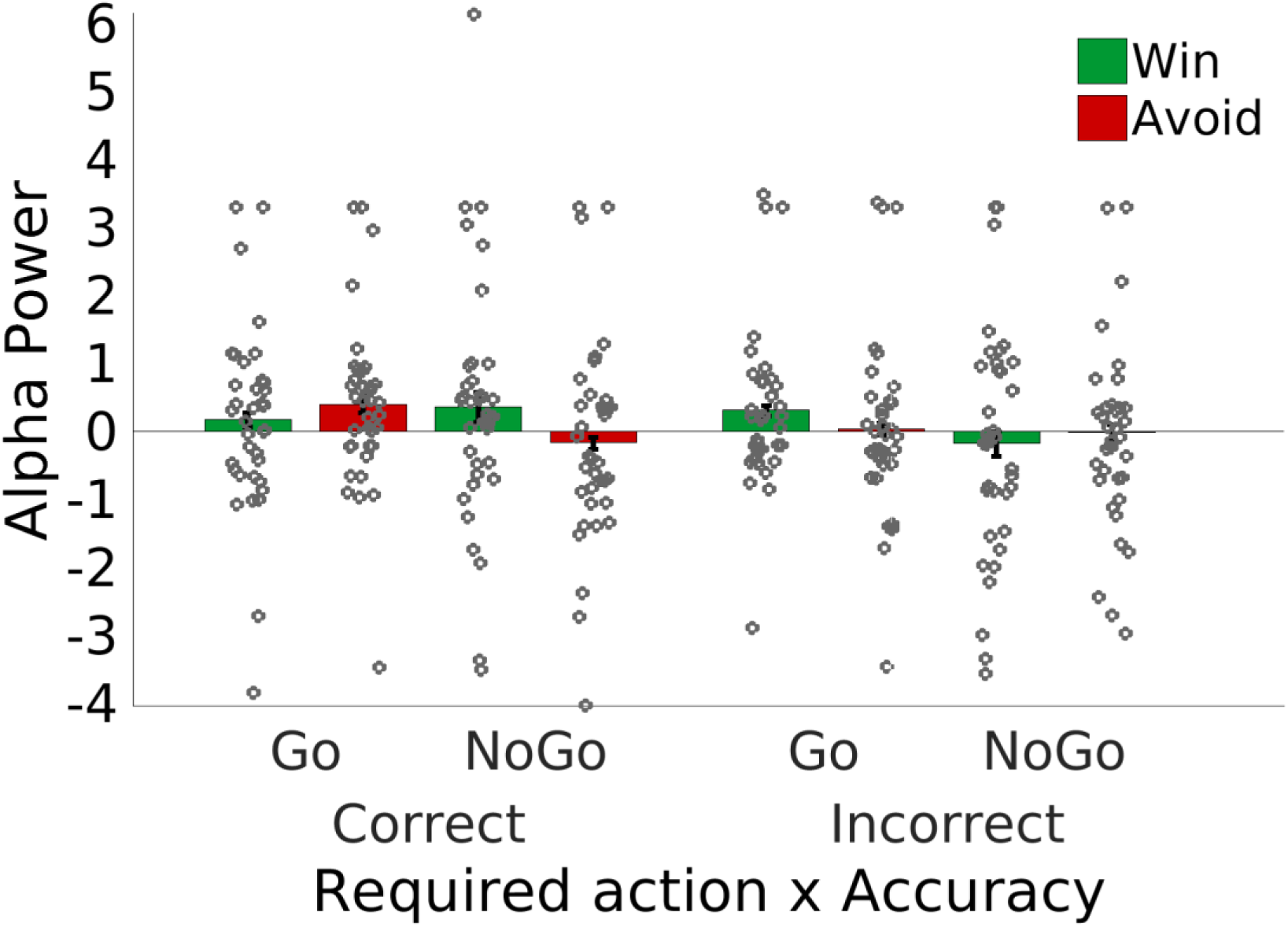
Alpha power (±SEM) as a function of cue valence, performed action, and accuracy over midfrontal electrodes (Fz/FCz/Cz). Alpha power selectively increased for correct bias-incongruent actions (correct Go2Avoid and NoGo2Win). Points are individual participant data points.

### S10: EEG TF power as a function of action and valence across correct and incorrect trials

While EEG results reported in the main text only include correct trials in order to avoid contamination by error-related activity, fMRI results include all trials while explicitly modeling error trials with a designated regressor. To match this fMRI analysis approach, we here report EEG analyses including both correct and incorrect trials, as well.

Results were highly similar to those of the correct trials only reported in the main text<: Broadband power (1–15 Hz) was again significantly higher on trials with Go actions than NoGo actions (cue-locked*: p* = .006; response-locked: *p* = .004): This difference between Go and NoGo actions occurred as a broadband-signal from 1–15 Hz, but peaked in the theta band (Fig. S9B and D). The topographies exhibited a bimodal distribution with peaks both at frontopolar (FPz) and central (FCz, Cz, CPz) electrodes (Fig. S9B and D). As visual inspection of Fig. S9C shows, theta power increased in all conditions until 500 ms post cue onset and then bifurcated depending on the action: For NoGo actions, power decreased, while for Go actions, power kept rising and peaked at the time of the response. This resulted in higher broadband power for Go versus NoGo actions for about 575– 1300 ms after cue onset (see Fig. S8C; around -150–475 ms when response-locked, see Fig. S9A). When looking at the cue-locked signal, the signal peaked earlier and higher for Go actions to Win than to Avoid cues on correct trials, but not on incorrect; hence, when testing for differences in broadband power between Win cues and Avoid cues, broadband power was not significantly different between Win and Avoid cues. This difference in latency and peak height of the ramping signal was not present in the response-locked signal, and the respective test of Win vs. Avoid cues not significant either.

**Figure S10.**
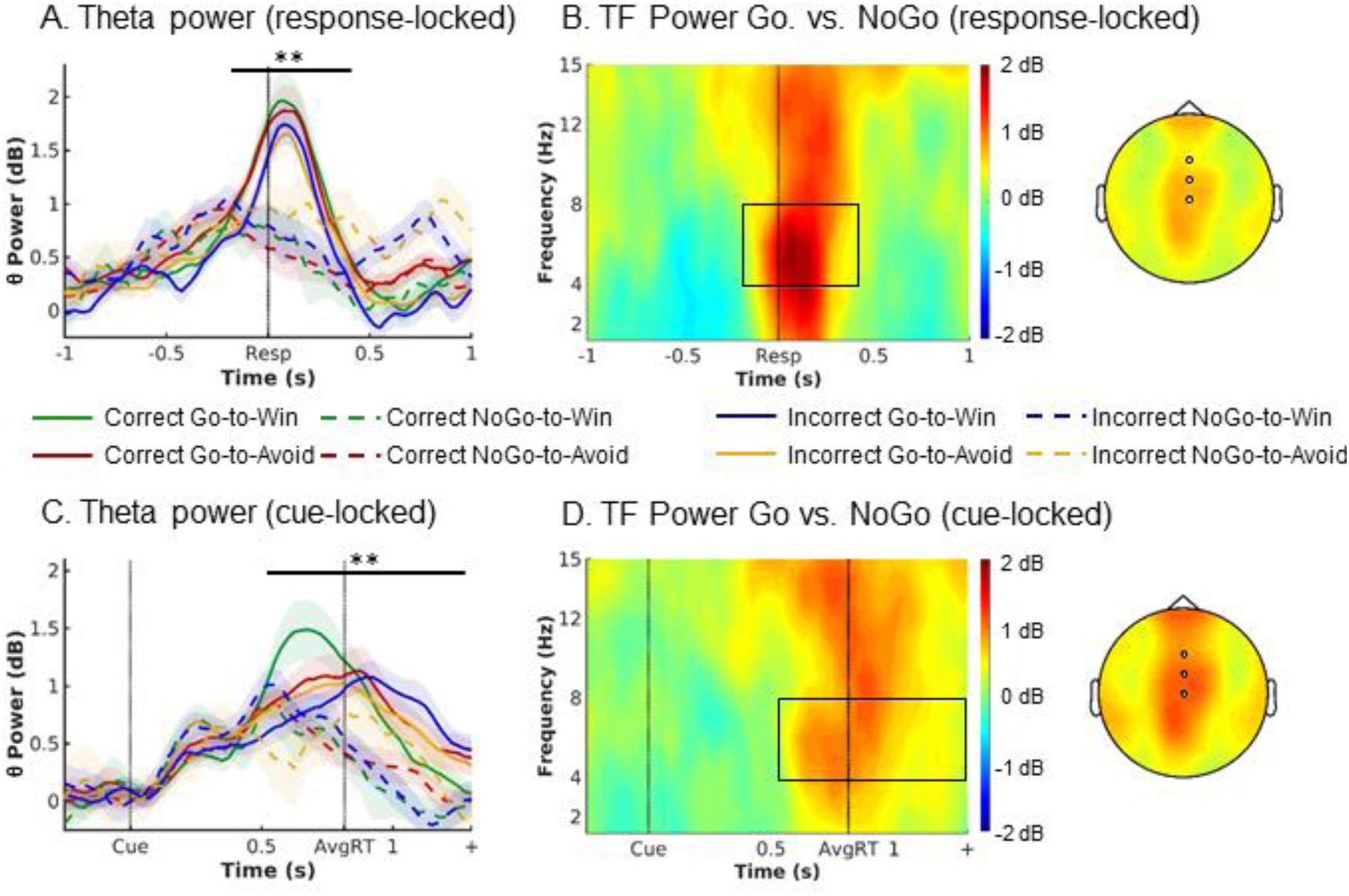
EEG time-frequency power as a function of cue valence and action for both correct and incorrect trials. (A) Response-locked within trial time course of average theta power (4–8 Hz) over midfrontal electrodes (Fz/ FCz/ Cz) per cue condition (correct-trials only). Theta increased in all conditions relative to pre-cue levels, but to a higher level for Go than NoGo trials. There were no differences in theta peak height or latency between Go2Win and Go2Avoid trials. (B) Left: Response-locked time-frequency power over midfrontal electrodes for Go minus NoGo trials. Go trials featured higher broadband TF power than NoGo trials. The broadband power increase for Go compared to NoGo trials is strongest in the theta range. Right: Topoplot for Go minus NoGo trials. The difference is strongest at FZ and FCz electrodes. (C-D) Cue-locked within trial time course and time-frequency power. Theta increased in all conditions relative to pre-cue levels, but to a higher level for Go than NoGo trials, with earlier peaks for Go2Win than Go2Avoid trials. * *p* < 0.05. ** *p* < 0.01. Shaded errorbars indicate (±SEM). Box in TF plots indicates the time frequency window where *t*-values > 2.

### S11: Plots and tests of the evidence accumulation hypothesis

Previous research has suggested theta oscillations to reflect midfrontal mechanisms that lead to an elevation of the response threshold in basal-ganglia action selection mechanisms (Cavanagh et al., 2011; Cavanagh & Frank, 2014; Cohen, 2014; Frank et al., 2015). However, an alternative interpretation of past findings could be that theta reflects the subcortical action selection process itself (Bland & Oddie, 2001; Caplan et al., 2003; DeCoteau et al., 2007a, 2007b; Womelsdorf, Vinck, Leung, & Everling, 2010). This reasoning would explain why in motor tasks, over the entire trial time course, theta oscillations strongly increase in any task condition, even in absence of conflict (Cohen & Cavanagh, 2011; Swart et al., 2018). In conflict situations warranting elevated response thresholds, this process continues beyond normal levels and evolves for an extended time period, leading to the typical “conflict-related theta” reported in the literature.

In fact, several characteristics of the theta signal we observed resembled an accumulating evidence process as evidenced by additional tests of systematic different in peak height and latency of the signal (following O’Connell et al., 2012). In our case, features of the theta signal would be consistent with evidence selectively accumulated for making a Go action:

First, such a process should rise early (when Go is still a considered option) in all trials, but deactivate when the final response is NoGo, while it should keep rising when the final response is Go. This prediction is in line with our observations (see Fig. 3C main text).

Second, we found the theta signal to scale with reaction times, such that the signal peaked earlier on trials with earlier response times. This link would be expected when a signal causes the a response, such that the latency of the signal peak determines reaction times (O’Connell et al., 2012). To test this hypothesis in our data, we split up each participant’s trials with Go actions (correct trials only) into three equally sized bins of fast, medium, and slow reaction times (tertials), separately for Win and Avoid trials (to account for inherent differences in reaction times between Win and Avoid trials). We then computed the average stimulus-locked signal in the theta range for each bin and determined the time point between 0.3 (fastest responses) and 1.3 s (slowest possible responses) at which the signal (first) reached its peak. We then used one-tailed paired-samples *t*-tests to test whether the signal peaked earlier in bins with faster reaction times, separately for Win and Avoid trials. Overall, the signal peaked earlier for Win trials (M = 0.697, SD = 0.175) than Avoid trials (M = 0.787, SD = 0.264), *t*(35) = 2.335, *p* = 0.013, *d* = 0.39. This difference was selective for the theta band (Fig. S12). For Win trials, indeed, the signal peaked significantly earlier for faster than for medium reaction times, *t*(35) = 1.745, *p* = 0.045, *d* = 0.291, and significantly earlier for medium than late reaction times, *t*(35) = 2.577, *p* = 0.007, *d* = 0.430 (see Fig. S11A). For Avoid trials, the signal only peaked marginally significantly earlier for faster than for medium reaction times, *t*(35) = 1.662, *p* = 0.053, *d* = 0.277, but significantly earlier for medium than late reaction times, *t*(35) = 5.220, *p* < 0.001, *d* = 0.870. Conclusions were identical when using non-parametric permutation tests instead of *t*-tests. In sum, the peak latency of the stimulus-locked theta signal scaled with reaction times, as expected for a signal triggering actions.

Third, when response-locked, differences in peak latency and height between cue valence conditions and reaction time bins disappeared, in line the assumption of a fixed threshold that evidence must reach in order to trigger action release (O’Connell et al., 2012). To investigate systematic differences in peak latency, we computed the average response-locked signal in the theta range for each bin for each participant and determined the time point between 0.5 s before and 0.5 s after the response at which the signal (first) reached its peak. We again compared bins within each valence condition using two-tailed *t*-tests. For Win trials, there were no significant differences in peak latency between fast and medium reaction times, *t*(35) = -0.784, *p* = 0.438, *d* = -0.131, medium and slow reaction times, *t*(35) = 0.896, *p* = 0.376, *d* = 0.149, or fast and slow reaction times, *t*(35) = 0.135, *p* = 0.894, *d* = 0.023 (see Fig. S11B). Similarly for Avoid trials, there were no significant differences in peak latency between fast and medium reaction times, *t*(35) = 0.014, *p* = 0.988, d = 0.002, medium and slow reaction times, *t*(35) = 0.347, *p* = 0.730, *d* = 0.058, or fast and slow reaction times, *t*(35) = 0.245, *p* = 0.808, *d* = 0.041. Conclusions were identical when using non-parametric permutation tests instead of *t*-tests. To investigate significant differences in peak height, we extracted the height of the theta signal at the peak latency within each bin for each participant. We again compared bins within each valence condition using two-tailed *t*-tests. For Win trials, were no significant differences in peak height between fast and medium reaction times, *t*(35) = -1.138, *p* = 0.264, *d* = -0.190, medium and slow reaction times, *t*(35) = 1.003, *p* = 0.322, *d* = 0.167, or fast and slow reaction times, *t*(35) = 0.206, *p* = 0.838, *d* = 0.034 (see Fig. S11B). Similarly for Avoid trials, there were no significant differences in peak height between fast and medium reaction times, *t*(35) = -0.017, *p* = 0.986, *d* = -0.003, medium and slow reaction times, *t*(35) = 0.195, *p* = 0.846, *d* = 0.033, or fast and slow reaction times, *t*(35) = 0.195, *p* = 0. 846, *d* = 0.033. Conclusions were identical when using non-parametric permutation tests instead of *t*-tests. Taken together, there were no significant differences in peak latency and height between different reaction times, in line with a fixed response threshold independent of response time or cue valence.

Fourth, previous research on EEG correlates of evidence accumulation in perceptual decision making has found that incorrect responses were elicited at systematically lower thresholds than correct responses (O’Connell et al., 2012), suggesting that trial-by-trial variation in the response threshold can cause erroneous action releases. To test this hypothesis in our data, we computed the response-locked theta signal separately for correct and incorrect Go actions on Win and Avoid trials, and then computed the peak height for each participant. We used one-tailed *t*-tests to test whether peak height was lower for incorrect than correct trials. This was indeed the case both on Win trials, *t*(35) = 2.558, *p* = 0.008, *d* = 0.426, and Avoid trials, *t*(35) = 2.729, *p* = 0.005, *d* = 0.455 (Fig. S11C). As an alternative, we performed a cluster-based permutation test contrasting correct and incorrect responses. Both were indeed significantly different, p = 0.047, most dominantly from 125 ms before until 25 ms after around responses. In conclusion, we found evidence in line with the hypothesis that false-positive action releases occur at a systematically lower response threshold than true-positive action releases.

Fifth, our interpretation of theta as reflecting evidence accumulation is in line with previous research that has found perceptual evidence to be reflected in the theta band (van Vugt, Simen, Nystrom, Holmes, & Cohen, 2012). Also, a recent study observed both perceptual and value-based evidence encoded in the gamma band in topographies very similar to the one we found, with peaks in both frontopolar and centroparietal electrodes (Polanía, Krajbich, Grueschow, & Ruff, 2014). It is possible that feedforward activity encoded in the gamma band is nested in lower-frequency theta cycles reflecting top-down integration (Canolty et al., 2006; Landau, Schreyer, van Pelt, & Fries, 2015; Maris, van Vugt, & Kahana, 2011). In conclusion, our results are in line with theta power reflecting an evidence accumulation process for deciding whether to perform an active Go response.

**Figure S11A.**
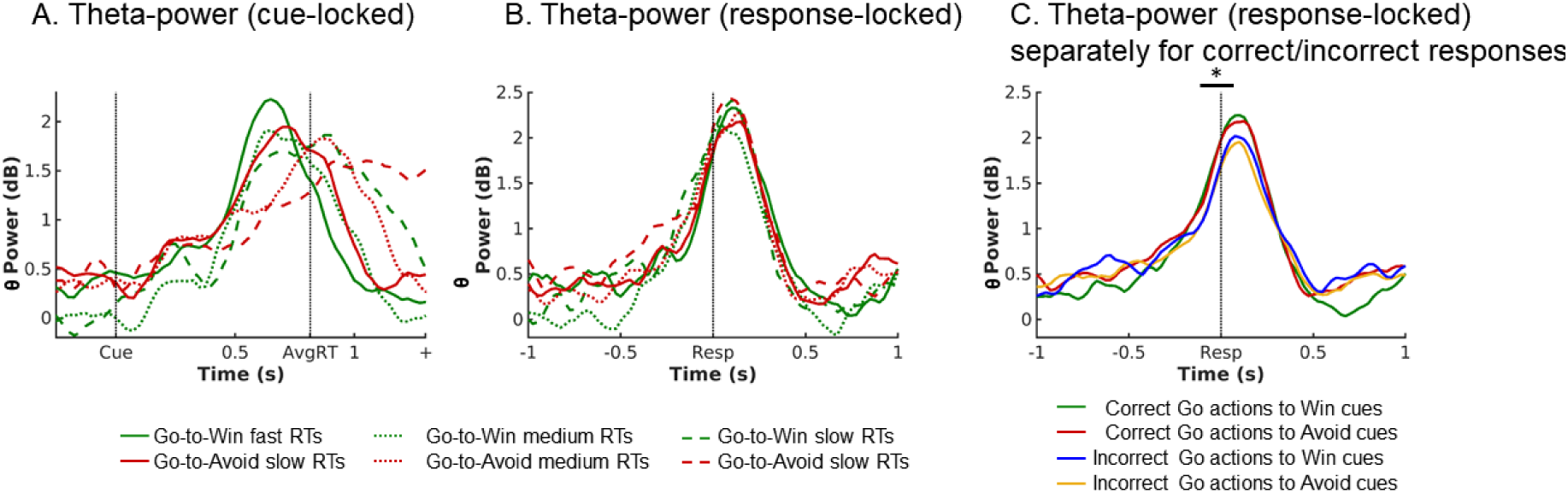
Plots displaying features of the theta signal akin to an evidence accumulation process for active Go responses. (A) The stimulus-locked theta signal split into fast, medium, and slow reaction time bins separately for Win and Avoid trials. The signal peaks systematically earlier for earlier reaction times. (B) The same signal response-locked. Differences in peak height and latency between reaction time bins are absent. (C) Correct and incorrect Go actions separately for Win and Avoid trials. The theta peaks at significantly lower levels for incorrect compared to correct actions.

**Figure S11B.**
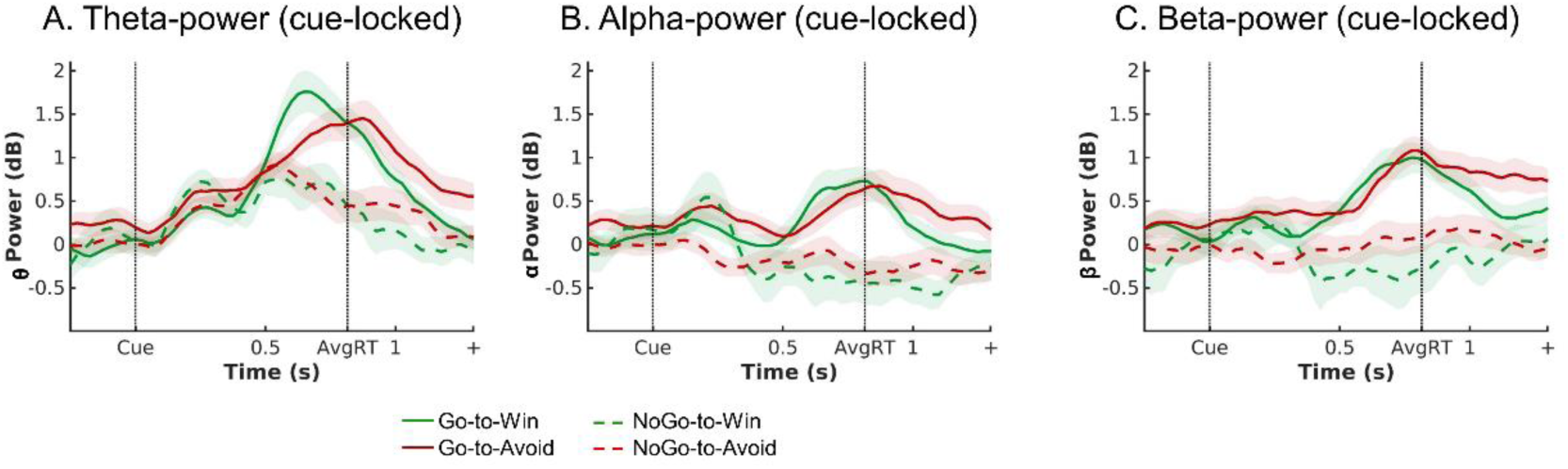
Plots displaying theta (A), alpha (B), and beta (C) power over midfrontal channels (Fz/ FCz/ Cz) split per cue-valence and performed action (correct trials only to avoid contamination error processing). Signal rise is strongest in the theta band. Also, the positively accelerating shape typically observed in evidence accumulation signals (Donner, Siegel, Fries, & Engel, 2009; O’Connell et al., 2012) is only observed in the theta band. Finally, differences in peak latency between Win and Avoid cues (in line with differences in reaction times between conditions) only arise in the theta band.

### S12: Correlation of EEG power with head motion

Recently, Fellner and colleagues (2016) reported that lower-frequency oscillations in simultaneous EEG-fMRI studies were affected by head motion: Realignment parameters strongly correlating with time-frequency power. These authors recorded simultaneous EEG and fMRI during encoding and retrieval phases of a memory task. They found difference in theta oscillations for remembered vs. forgotten items that had the opposite sign compared to when the same task was performed with EEG outside the MR scanner. Also, they found overall lower-frequency oscillations, especially in the theta range, to be strongly correlated with a summary measure of the six realignment parameters, casting doubt on the neural origin of theta effects measured in simultaneous EEG-fMRI recordings.

We leveraged our approach of fMRI-inspired EEG analysis approach (see main text) by computing the same summary statistic as Fellner and colleagues (2016) based on the six realignment parameters for each volume, upsampling this signal to a TR of 0.140 s, and then downsampling the signal again into epochs of 2 s length relative to trial onset, yielding an indicator of overall head motion during each trial.

First, similar to Fellner and colleagues (2016), we compared the head motion between trials with Go actions and trials with NoGo trials by computing the average head motion summary statistic for such trials for each participant and then performing a two-tailed paired-samples *t*-test. There was no significant difference in head motion between trials with Go and trials with NoGo actions, *t*(35) = 1.614, *p* = 0.116, *d* = 0.269. Similarly, there was no significant difference in head motion between Win and Avoid trials, *t*(35) = -0.467, *p* = 0.643, *d* = -0.078, nor between congruent and incongruent trials, *t*(35) = -0.304, *p* = 0.763, *d* = -0.051. These results suggest that head motion did not significantly differ between experimental conditions.

Next, we used the head motion summary statistic as a trial-by-trial predictor of time-frequency power in multiple regression, similarly to the BOLD signal extracted from neural regions. When head motion was used as a sole regressor, we indeed observed a significant positive correlation with theta/ delta power (*p* = 0.039). However, this correlation was not spread out in time and frequency space as in Fellner and colleagues (2016), but instead focused on theta/ delta power around 0.9–1.3 s. after cue onset—i.e. after the average response time (see Fig. S12A). When entering BOLD signal from the selected regions as additional regressors, this pattern remained similar but became non-significant (*p* = .089).

Given that participants were instructed to perform Go actions only while the respective cue was visible (0–1.3 s after cue onset), we restricted our analyses of task-related neural signals to this period. However, head motion-related signals might occur even after this period. When performing fMRI-inspired EEG analyses on a window of 0–2 s after cue onset, we indeed observed a strong correlation (*p* = 0.005) of head motion with broadband time-frequency power after cue offset (after 1.3 s, see Fig. S12B; especially so when including the five participants that were otherwise excluded from the fMRI-informed EEG analyses; see Fig. S12C). This finding suggests that head-motion (and associated artifacts) might predominantly occur during the inter-stimulus interval when participants have performed any cue-related action and wait for the outcome. Note that the trials in Fellner and colleagues (2016) were much longer (3 s) than in our paradigm. Hence, head motion might be a particular problem in EEG-fMRI studies with paradigms featuring long trial durations, unlike the cue presentation phase in our paradigm.

Correlations of time-frequency power with BOLD and task factors remained significant even when head motion was included in the regression: First, we still observed a significant correlation of striatal BOLD with late theta/ delta power 0.5–1.0 s after cue onset (*p* = .045), suggesting that striatal BOLD predicts theta power independently of any head motion-related signals (see Fig. S12D). Second, the correlations of left (*p* = .008) and right (*p* = .027) motor cortex (see Supplementary Material S11) and vmPFC (*p* = .035) BOLD with time-frequency power remained significant. Third, inspecting the beta-map of the additional regressor Go vs. NoGo responses (coded as 1 and 0, regressor demeaned), which we included in all these analyses by default, revealed correlations with broadband signal around the time of responses (*p* = .026), a pattern very similar to the EEG-only analyses (see Fig. S12E; compare to Fig. 3D in the main text). This finding suggests that the broadband signal associated with Go vs. NoGo responses in EEG-only analyses is not reducible to head motion artifacts either.

Finally, when using the trial-by-trial theta power as a regressor in an fMRI GLM (see Supplementary Material S12), we observed theta correlates in action-related regions such as ACC, motor cortices, opercula, putamen, and cerebellum, unlike Fellner and colleagues (2016) who observed correlates mostly in regions of the default-mode network. We thus conclude that the theta effects we observed constitute task-related neural signals rather than the head-motion related artifacts described by Fellner and colleagues (2016).

**Figure S12.**
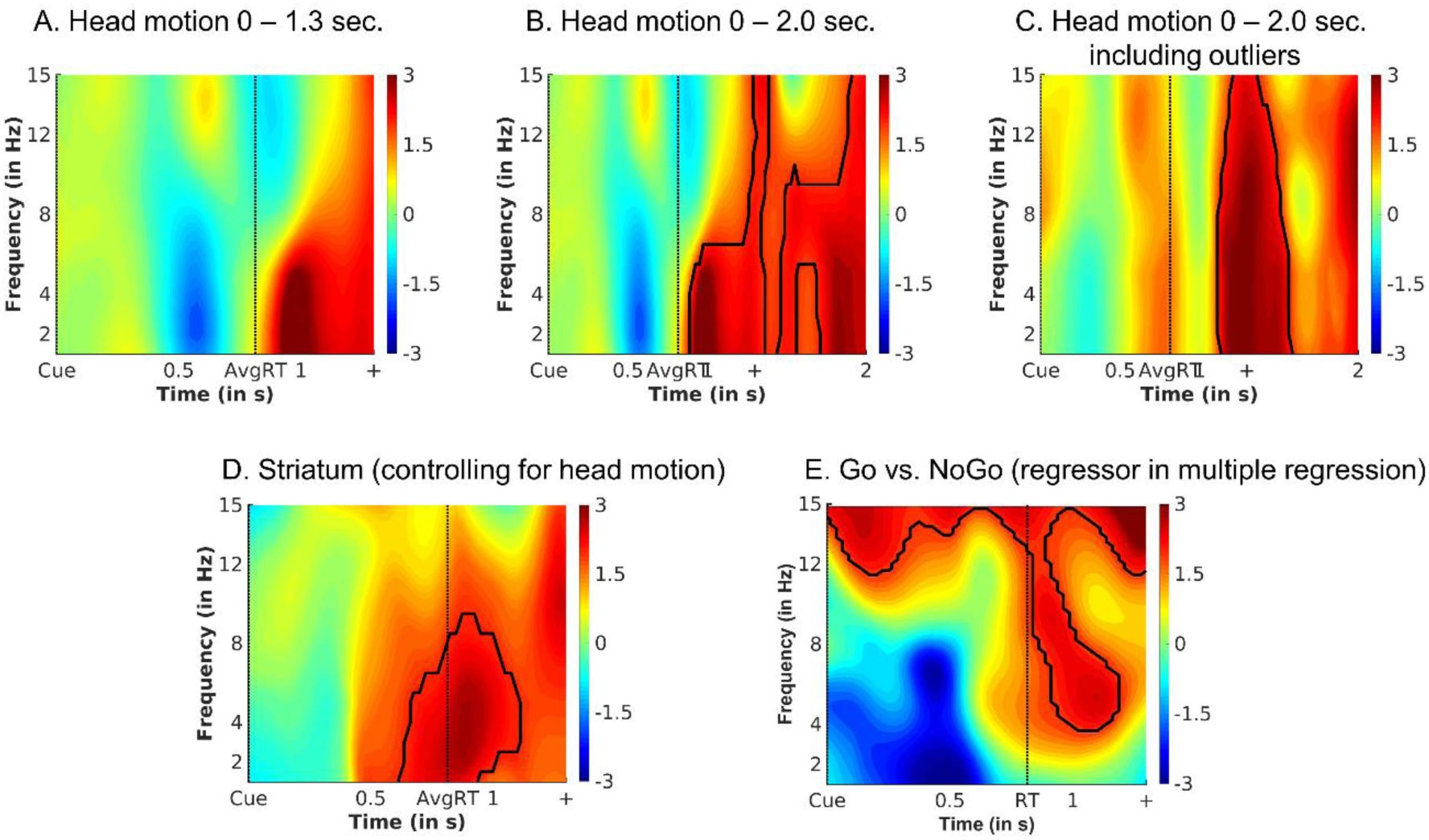
Time-frequency correlates of head motion and task effects when controlled for head motion. (A) Head motion was quantified via relative displacement as a summary statistic of the volume-by-volume realignment parameters, which was first upsampled, then downsampled into a single value per trial, and used to predict average time-frequency power over midfrontal electrodes. Head motion correlated (though not significantly) with theta/ delta power around 1 sec. after stimulus onset. (B) The same analyses on a time window of 0–2.0 revealed that head motion predominantly correlated with broadband time-frequency power after cue offset. (C) Same plot as (B), but with the four participants included that are typically excluded due to out-of-range regression weights and strong head motion. (D) The correlation of striatal BOLD with theta/ delta power around the time of responses remained unaltered when entering head motion as an additional regressor into the model. (E) Using Go vs. NoGo responses (coded as 1 and 0, demeaned) as an additional regressor yielded again an increase in broadband time-frequency power for Go compared to NoGo responses (see main text Figure 3D), even when entering head motion as an additional regressor into the model. Areas surrounded by a black edge indicate clusters of |*t*| > 2 with *p* < .05 (cluster-corrected).

### S13: Increase in time-frequency power relative to baseline for each cue valence x performed action pairing

**Figure S13.**
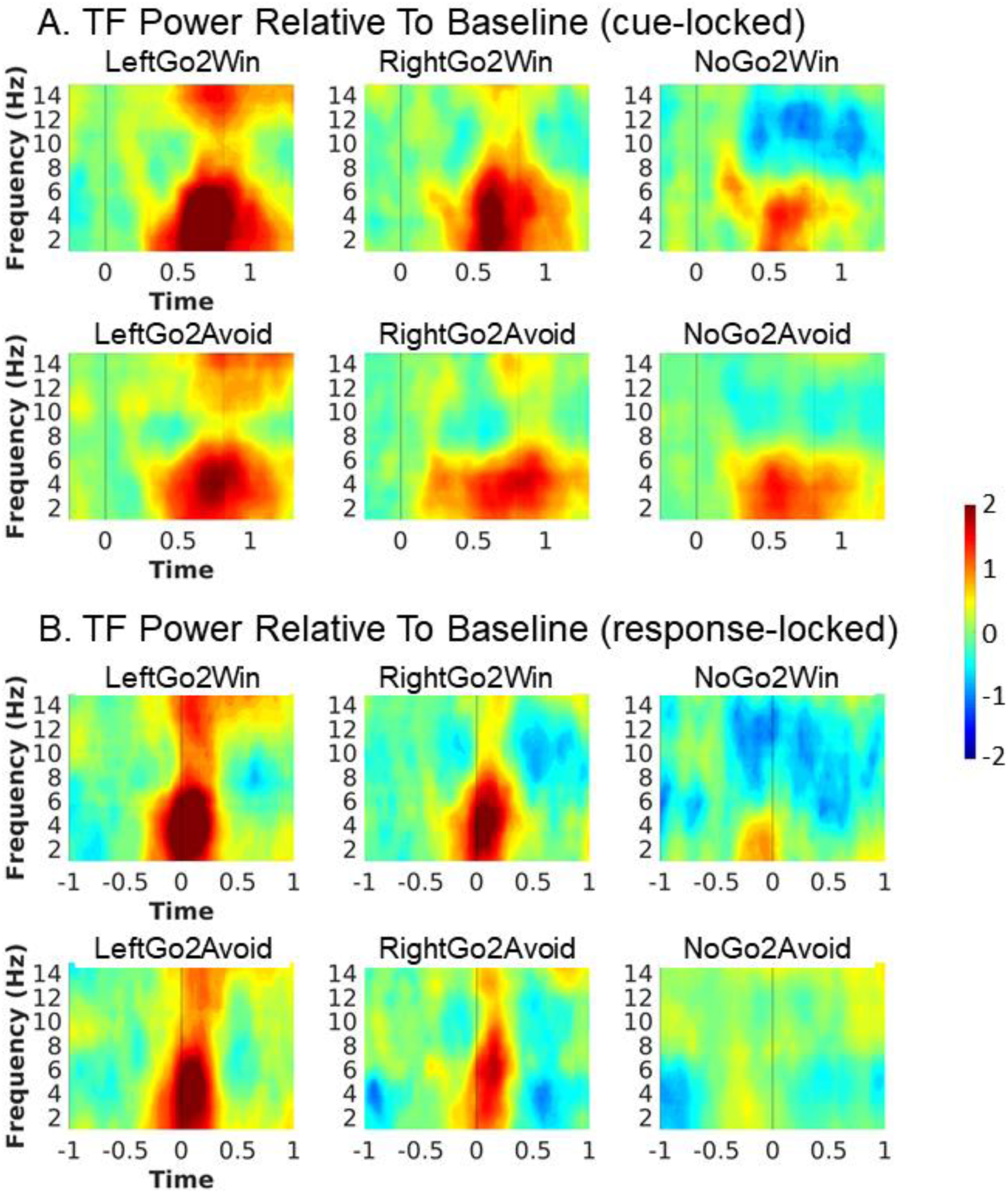
Average time-frequency power over midfrontal electrodes (Fz/FCz/Cz) relative to baseline (mean signal - 250–50 ms before cue onset), split up per cue valence (rows) and performed action (columns), both stimulus-locked (A) and response-locked (B). For stimulus-locked plots, vertical solid lines indicate cue onset, dashed vertical lines average response time. For response-locked plots, vertical solid lines indicate response time (i.e. 0 s). Time-frequency power in the delta/ theta range increases in every valence-action pairing, also for NoGo actions, which rules out that this signal is a mere motion artifact.

### S14: Theta and beta power for left vs. right hand responses

The strong association of broadband power with motor activity opened the possibility that this signal was potentially an artifact of EEG acquisition (e.g. head movement in the scanner) rather than a neural signal (Fellner et al., 2016). If the signal constituted an artifact, one would expect it to be symmetrical for both hands. One the other hand, if it was a neural signal reflecting the level of evidence accumulated before initiating a response, one might expect the signal to be sensitive to differences in evidence thresholds between hands. Given that all our participants were right-handed, one might expect that responses of the left (non-dominant) hand were less easily initiated and required a higher level of evidence to be selected than responses of the right (dominant) hand. A broadband permutation test (stimulus-locked: *p* = .020; response-locked: *p* = .006) indicated that power in the theta band (around -250–25 ms relative to responses, see Fig. S13) and in the beta-band (around -150– 675 ms relative to responses (see Fig. S13B and D) was in fact higher for left-hand than right-hand responses.

This modulation of the theta signal by handedness corroborates the interpretation that the observed theta synchronization is of neural origin. It might reflect a bias towards right-hand responses, such that left-hand responses require a higher level of evidence to be initiated than right-hand responses. Such a right-hand bias might also explain why synchronization in the beta band was higher for left than right hand responses: Given that beta synchronization is typically found for motor inhibition (Wessel et al., 2016; Wessel, Waller, & Greenlee, 2019), higher beta on trials with left-hand responses might reflect that the right hand needed to be actively suppressed on these trials (see also Supplementary Material S11). In fact, we also observed that left-hand responses (*M* = 0.772) were overall slower than right hand responses (*M* = 0.745), χ^2^(1) = 6.709, *p* = .010, which further corroborates the interpretation that left-hand responses might have been harder to perform than right-hand responses.

**Figure S14.**
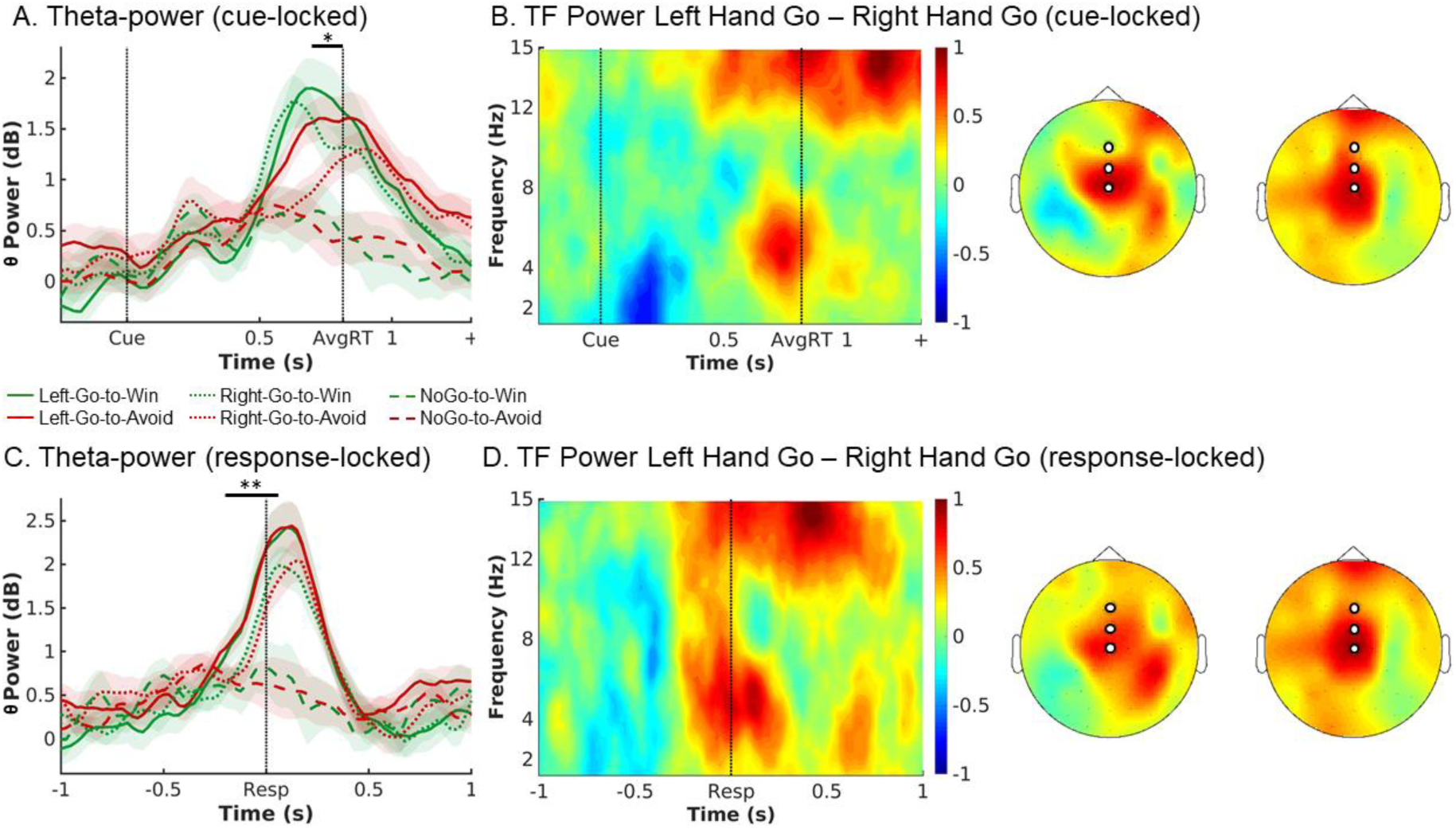
Theta and beta power split up for left vs. right hand responses. (A) Trial time course of average (±SEM) theta power (4–8 Hz) over midfrontal electrodes (Fz/FCz/Cz) per split up for left- and right-hand responses (correct-trials only; stimulus-locked). Theta increased to a higher extend for left-hand than right-hand responses. * *p* < 0.05. (B) Left: Time-frequency power over midfrontal electrodes for left-hand minus right-hand responses trials. Left-hand responses were associated with high theta and beta power compared to right-hand responses. Right: Topoplot for left-hand minus right-hand responses in the theta (left) and beta (right) range. (C-D) Same data when response-locked. Differences in theta peak latency between Go2Win and Go2Avoid trials disappear. ** *p* < 0.01.

### S15: Supplementary fMRI-inspired EEG results in time-frequency space

In addition to EEG correlates of BOLD signal in the striatum, ACC and vmPFC (see main text), we also observed correlates for BOLD in left and right motor cortex. Both left motor cortex (two separate clusters, both *p* = .030 and *p* = .030, cluster-corrected) and right motor cortex (*p* = .001 cluster-corrected) exhibited overlapping, but oppositely signed correlates in the alpha/beta band, with left motor cortex correlating negatively with midfrontal beta power (around 12–15 Hz, 0.6–1.3 s, Fig. S14B), while right motor cortex correlated positively with alpha/beta power (around 10–15 Hz, 0.6–1.3 s, Fig. S14C). These findings mirror the observation of higher theta and beta power for left hand (i.e. right motor cortex) compared to right hand (i.e. left motor cortex) responses (see Supplementary Material S9), again suggesting that executing a left hand response might have required an active suppression (associated with increased beta power) of the right hand. These results corroborate that theta power does not reflect motor preparation/ execution signals from the motor cortices, but signals from distinct regions. Furthermore, these associations replicate numerous intracranial and source-localization studies (Salmelin, Forss, Knuutila, & Hari, 1995; Salmelin, Hämäläinen, Kajola, & Hari, 1995; Sanes & Donoghue, 1993; Stolk et al., 2019) and previous EEG-fMRI studies (Jurkiewicz, Gaetz, Bostan, & Cheyne, 2006; Ritter, Moosmann, & Villringer, 2009) reporting beta oscillations in motor cortices. The presence of this well-established BOLD-EEG association corroborates the robustness of the data and analysis.

We performed a range of follow-up analyses to check for the robust of our results. We reached similar results and identical conclusions when a) performing regressions with each region as the sole predictor, b) including a summary measure of the realignment parameters as a proxy for head motion into the regression (Fellner et al., 2016), and c) when fitting HRFs for all trials of a certain block within a single GLM instead of separately for each trial.

We lastly aimed to test whether distinct striatal subregions with opposite valence coding, i.e. left putamen (Win > Avoid) and bilateral medial caudate (Avoid > Win), showed distinct time-frequency correlates. When using BOLD from those subregions instead of overall striatal BOLD as regressors, left putamen BOLD did not exhibit a significant association with time-frequency power (*p* = .218), while medial caudate BOLD did significantly correlate with delta/theta power around 825–1,2500 ms post-stimulus (*p* = .011). The cluster of significant correlations observed for medial caudate was highly similar to the cluster observed as a correlate of the entire striatum. Descriptively, both ROIs showed clusters of positive correlations with theta/ delta power around the time of responses, and slightly earlier so for the left putamen than for medial caudate. This finding would be in line with the idea of left putamen more strongly driving Go responses on Win trials, which showed shorter RTs, and medial caudate rather driving Go responses on Avoid trials, which showed longer RTs. However, as clusters associated with those regions were small and permutation tests not significant, this descriptive finding should be interpreted with caution.

**Figure S15.**
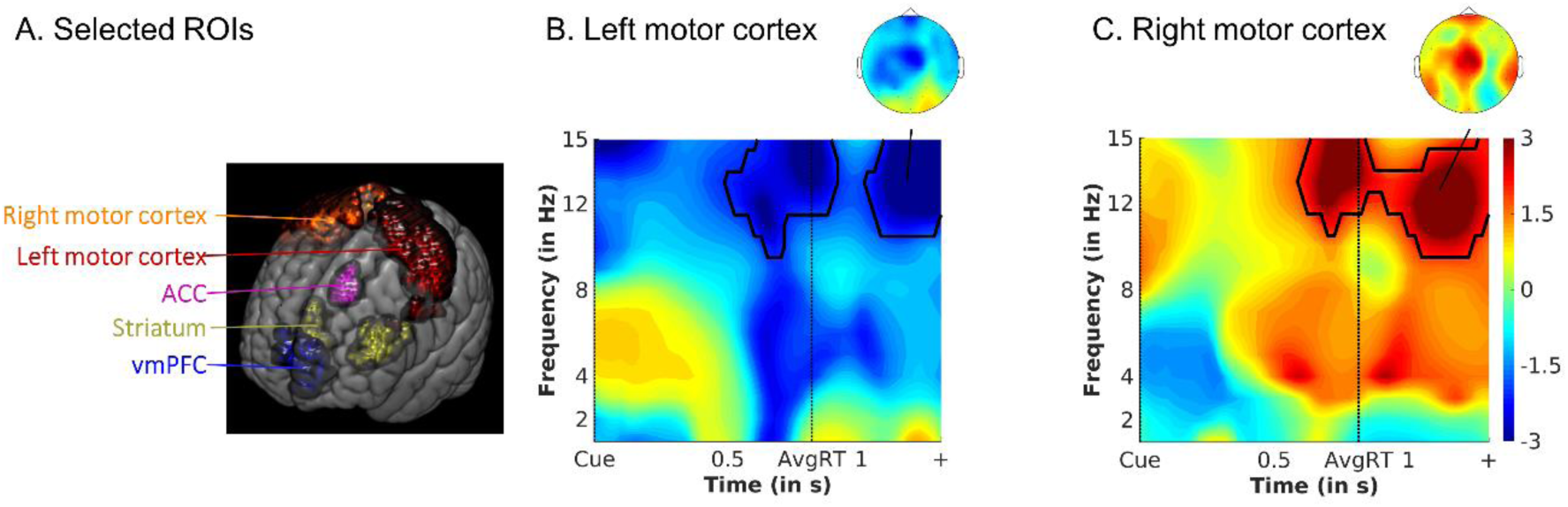
(A) Selected ROIs from which trial-by-trial HRF amplitudes were used as predictors in a multiple regression of midfrontal EEG time-frequency power. (B-C) Unique temporal correlation of BOLD signal in (B) left and (C) right motor cortex to average EEG time-frequency power over midfrontal electrodes (FCz/Cz). Group-level *t*-maps display the modulation of the EEG time-frequency power by trial-by-trial BOLD signal in the selected ROIs. Midfrontal beta power correlates negatively with BOLD in left motor cortex (more active for right hand responses), but positively with BOLD in right motor cortex (more active for left hand responses), putatively indexing response conflict and inhibitory processes when left hand responses were executed. Areas surrounded by a black edge indicate clusters of |*t*| > 2 with *p* < .05 (cluster-corrected). Topoplots indicate the topography of the respective cluster.

### S16: Supplementary fMRI-inspired EEG results in time space (ERPs)

Given that the time-frequency correlate of trial-by-trial vmPFC BOLD occurred very early after cue onset and was extended in frequency space, we hypothesized that vmPFC BOLD might be correlated with evoked rather than induced activity, which, when analyzed in time-frequency space, smeared across frequencies. We used the same approach for fMRI-informed EEG analyses as reported in the main text, but with the voltage signal (time-domain) instead of time-frequency power as dependent variable. We again used BOLD signal from striatum, ACC, left and right motor cortex, and vmPFC as simultaneous predictors in one single multiple regression.

When restricting analyses to midfrontal electrodes (FCz/ Cz), we found no significant modulation of EEG voltage by vmPFC BOLD (*p* = .260; see Fig. S15A and C). However, when considering a broader frontal ROI (F1/F3/FCz/FC1/FC3/ Cz/C1/C3), vmPFC appeared to attenuate the amplitude of the P2 component over left frontal electrodes (two clusters above threshold: *p* = .021 around 213–269 ms; *p* = .003 around 349 – 410 ms; see Fig. S15B and D). The topography of EEG voltage modulation did not exactly match with the topography of the time-frequency power modulation, but was rather restricted to left frontal electrodes (see Fig. S15C and E). Interestingly, these electrodes also showed the peak modulation of the P2 by Go compared to NoGo actions (see Supplementary Material S4). In conclusion, we found inconclusive evidence regarding whether broadband power decreases associated with vmPFC BOLD were reducible to evoked activity (modulation of the P2) or not.

**Figure S16.**
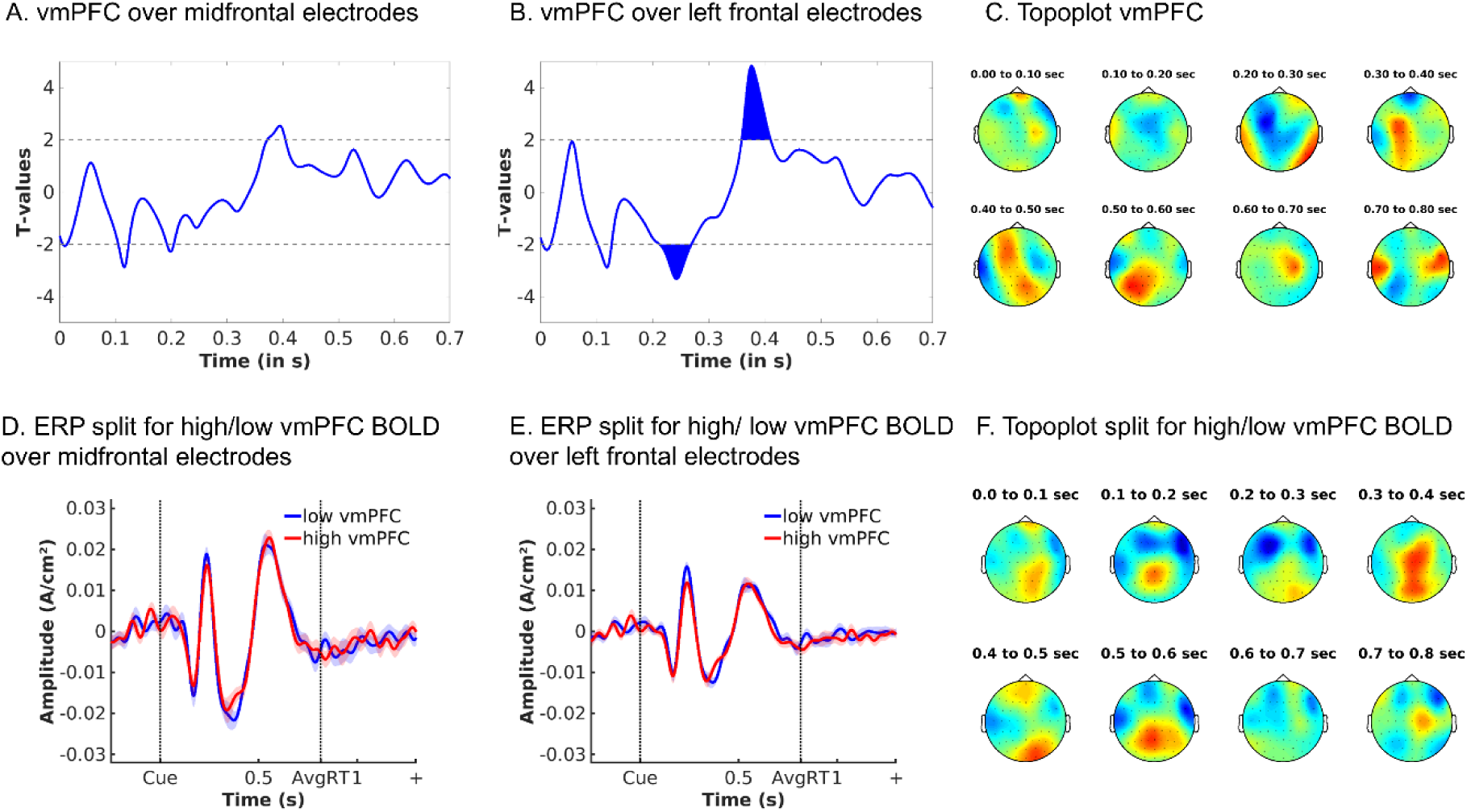
Modulation of EEG voltage by vmPFC BOLD signal. (A) Average EEG voltage over midfrontal electrodes (FCz/ Cz) was not significantly modulated by vmPFC BOLD, while (B) EEG voltage over left frontal electrodes (F1/ F3/ FCz/ FC1/ FC3/ FC5/ Cz/ C1/ C3) was. Filled areas indicate clusters of |*t*| > 2 with *p* < .05 (cluster-corrected). (C) Topoplots displaying *t*-values over the entire scalp in steps of 100 ms from 0 to 800 ms. The strongest modulation of frontal EEG voltage by vmPFC BOLD occurred over left frontal electrodes. This pattern does not fully match the topography of broadband power decreased associated with vmPFC BOLD (see Fig. 4C main text). (D) For plotting purposes, we sorted trials according to the trial-by-trial HRF amplitude in the vmPFC ROI and plotted the 33% of trials with highest vmPFC BOLD signal vs. the 33% trials with lowest vmPFC BOLD signal. This contrast indicates no strong difference a midfrontal electrodes (FCz/ Cz), but (E) does indicate an attenuation of the P2 component through high vmPFC BOLD over left frontal electrodes (F1/ F3/ FCz/ FC1/ FC3/ FC5/ Cz/ C1/ C3). (F) Topoplots displaying voltage for high vmPFC BOLD minus low vmPFC BOLD trials over the entire scalp in steps of 100 ms from 0 to 800 ms. The strongest modulation of frontal EEG voltage by vmPFC BOLD occurred over left frontal electrodes.

### S17: EEG-informed fMRI analyses

For the EEG-inspired fMRI analyses, we added trial-by-trial summary measures of conflict-related alpha power and action-related theta power to our GLM. These measures were created by using the 3-D (time-frequency-channel) *t*-map obtained when contrasting incongruent vs. congruent actions (Mask 1; stimulus-locked) and Go vs. NoGo actions (Mask 2; response-locked) over midfrontal channels (Fz/ FCz/ Cz) as a linear filter. We extracted those maps and retained all voxels with *t* > 2. We did not enforce strict frequency band cutoffs, but rather extracted the entire cluster of *t*-values above threshold. Restricting the action contrast *t*-map to the theta range or using the stimulus-locked rather than the response-locked map led to highly similar results and identical conclusions. These masks were applied to the trial-by-trial time-frequency data to create weighted summary measures of the average power in the identified clusters in each trial. Both resultant time series correlated only weakly (mean correlation across participants: *r* = .105). They were entered as parametric modulators on top of the task regressors as described above, with each regressor entering a separate parametric contrast.

The EEG alpha regressor correlated significantly negatively with BOLD in two clusters in left middle frontal gyrus/ lateral frontal pole and in right supramarginal gyrus. As Fig. S16A indicates, sub-threshold, the same areas in the respective other hemisphere also correlated negatively with trial-by-trial alpha, as did extensive areas in medial parietal/ occipital cortex. Overall, midfrontal alpha appeared to correlate negatively with extended areas that were part of the fronto-parietal and dorsal attention resting-state networks. Notably, no region correlated positively with midfrontal alpha.

The EEG theta regressor correlated significantly positively with BOLD in pre-SMA, ACC, bilateral precentral and postcentral gyrus, superior parietal lobule, precuneous, bilateral operculum, bilateral putamen, and bilateral cerebellum (see Fig. S16B). These regions also tended to be more active for Go than NoGo responses, corroborating the notion of theta reflecting evidence for active responses.

Notably, correlations with trial-by-trial theta power were not restricted to the striatum, but also occurred for other motor regions such as the ACC and motor cortices. These differences to the results of the fMRI-inspired EEG analyses might be attributable to methodological differences between both approaches: First, if theta power reflects global trial-by-trial brain activity associated with motion, this signal property will lead to correlations with BOLD in several motor regions in EEG-inspired fMRI analyses. In contrast, in fMRI-inspired analyses, such variance will be shared among regressors and thus be attributed to neither of them. Second, for EEG-inspired fMRI analyses, we created trial-by-trial indicators of theta power within a broad time window, which potentially mixes distinct events in theta that reflect activity in different brain regions. Thus, this approach might lead to correlations with BOLD in several brain regions that actually perform different computations at different time points. Following this reasoning, fMRI-inspired analyses have the advantage of a) identifying which regions uniquely predict time-frequency power beyond variance shared among regions, and b) unmixing different regions predicting time-frequency power at different time points.

**Figure S17.**
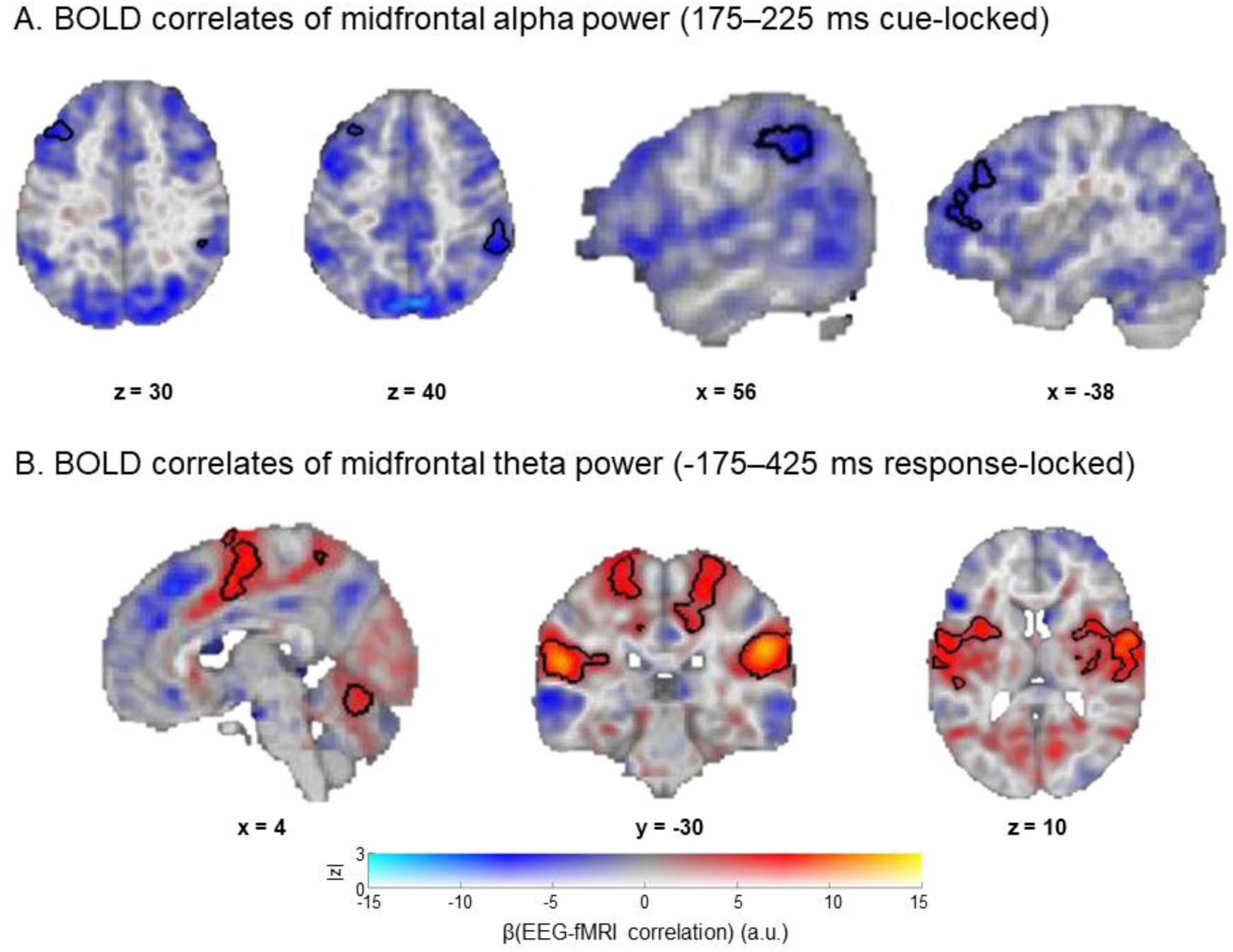
Trial-by-trial time-frequency power as a predictor of BOLD in an fMRI GLM. (A) Trial-by-trial midfrontal alpha power correlated significantly negatively with BOLD in two clusters in left middle frontal gyrus and in right supramarginal gyrus. Sub-threshold, it correlated negatively with extended regions in fronto-parietal and dorsal attention network. (B) Trial-by-trial midfrontal theta power correlated significantly positively with BOLD in pre-SMA, ACC, bilateral precentral and postcentral gyrus, superior parietal lobule, precuneous, bilateral operculum, bilateral putamen, and bilateral cerebellum.

## Notes

### Competing Interest Statement

The authors have declared no competing interest.

